# Building synthetic biosensors using red blood cell proteins

**DOI:** 10.1101/2023.12.16.571988

**Authors:** Taylor B. Dolberg, Taylor F. Gunnels, Te Ling, Kelly A. Sarnese, John D. Crispino, Joshua N. Leonard

## Abstract

As the use of engineered cell therapies expands from pioneering efforts in cancer immunotherapy to other applications, an attractive but less explored approach is the use of engineered red blood cells (RBCs). Compared to other cells, RBCs have a very long circulation time and reside in the blood compartment, so they could be ideally suited for applications as sentinel cells that enable *in situ* sensing and diagnostics. However, we largely lack tools for converting RBCs into biosensors. A unique challenge is that RBCs remodel their membranes during maturation, shedding many membrane components, suggesting that an RBC-specific approach may be needed. Towards addressing this need, here we develop a biosensing architecture built on RBC membrane proteins that are retained through erythropoiesis. This biosensor employs a mechanism in which extracellular ligand binding is transduced into intracellular reconstitution of a split output protein (including either a fluorophore or an enzyme). By comparatively evaluating a range of biosensor architectures, linker types, scaffold choices, and output signals, we identify biosensor designs and design features that confer substantial ligand-induced signal *in vitro*. Finally, we demonstrate that erythroid precursor cells engineered with our RBC protein biosensors function *in vivo.* This study establishes a foundation for developing RBC-based biosensors that could ultimately address unmet needs including non-invasive monitoring of physiological signals for a range of diagnostic applications.

## Introduction

Engineered cell-based therapies comprise a rapidly expanding field in medicine that has demonstrated remarkable clinical success, particularly as cancer therapeutics.^*1*^ An emerging potential frontier of this field is the engineering of red blood cells (RBCs) for therapeutic and diagnostic applications. RBCs are an attractive diagnostic platform for several reasons: they have an extensive circulatory range, exceptionally long circulation times (around 120 days in humans and 50 days in mice—far longer than synthetic vehicles), lack DNA (mitigating concerns of genetic alteration or abnormal growth in humans), and have a large surface area that could be used to house membrane-based sensors (160 µm^2^).^*2*^ Recent biotechnological advances have enabled the large-scale production of RBCs from precursor cells,^*3–5*^ which may potentially be harnessed to generate off-the-shelf RBC products to meet medical needs. Since such precursors can also be genetically engineered to include additional technology, this also opens the exciting possibility of building new classes of engineered RBC (eRBC) products.

Historically, eRBC research has focused on chemically and genetically modifying RBCs to function as drug delivery vehicles. RBCs have been engineered to incorporate therapeutic species (such as enzymes,^*6, 7*^ insulin,^*8*^ methotrexate,^*9*^ and magnetic nanoparticles^*10*^) in the plasma membrane,^*11–13*^ and the RBC surface has been engineered to display various agents through the use of chemical cross-linkers^*14–17*^ and conjugation to amino acids,^*18–20*^ sugars,^*21*^ and lipids.^*22, 23*^ However, modifying RBCs with such methods risks damage to the RBCs and may reduce circulation time, biocompatibility, and therapeutic utility *in vivo.*^*2, 6, 7, 24*^ To mitigate this problem, Lodish et al. reported two genetic strategies for covalently and site-specifically attaching probes to the RBC surface.^*25, 26*^ The first method entailed engineering an RBC protein, Kell, with sortase motifs to enable sortase-mediated attachment and display of disease-associated autoantigens. Such eRBCs were shown to exploit the natural method of RBC clearance to induce antigen-specific tolerance, both prophylactically and therapeutically, in a mouse model of multiple sclerosis.^*26*^ The second method genetically fused nanobodies (single domain camelid antibodies) targeting botulinum neurotoxin A to the RBC proteins glycophorin A (GPA) and Kell.^*25*^ When administered in an animal model, these eRBCs conferred sustained prophylactic protection against botulinum neurotoxin without provoking an immune response. These pioneering efforts demonstrate the promise of engineering RBCs as a new class of cell product.

Building on these promising foundations, we sought to genetically endow RBC precursors with the novel ability to report on the presence of a physiological cue. Ultimately, eRBCs generated from such precursors could enable entirely new capabilities, such as long-term monitoring of bioanalytes *in situ* and therapeutic and diagnostic applications beyond the reach of current cell-based technologies. Although the field of synthetic receptor engineering has evolved substantially in recent years,^*27*^ no existing technology has enabled such efforts in RBCs. A key unique challenge in RBC engineering is that during the process of erythropoiesis to generate mature RBCs, cells shed substantial portions of their membranes through remodeling events driven by both internal processes and external (e.g., macrophage-mediated) modification of the cell surface.^*28*^ Such considerations motivated the use of proteins such as GPA and Kell, which are retained through such remodeling, in the aforementioned eRBC engineering efforts. However, given the complex mechanisms through which GPA and Kell achieve retention through interactions with membrane and cytoskeletal components^*28*^, it was unclear whether such proteins are amenable to conversion into biosensors.

This study focused on evaluating the feasibility of building biosensor systems based upon RBC-resident membrane proteins which are retained during erythropoiesis and maturation (**Figure 1**). We built and evaluated a panel of receptor designs employing a general ligand binding-induced dimerization mechanism to reconstitute an intracellular split protein, derived design principles, and validated that such a strategy was feasible for generating either fluorescent or enzymatic (bioluminescent or bioluminescence resonance energy transfer, BRET) output modalities. These biosensors function in both workhorse model cell lines and in the erythroid precursor G1ER cell line, which can be matured into an RBC-like phenotype. Engineered G1ER cells conferred nearly real-time responses to ligand *in vitro*, and similar high performance was observed in mice in an *in vivo* validation. Overall, this study establishes the feasibility of RBC protein-based biosensors as a viable technology for eRBC development, which could ultimately address an unmet need for a range of applications.

**Figure 1.**
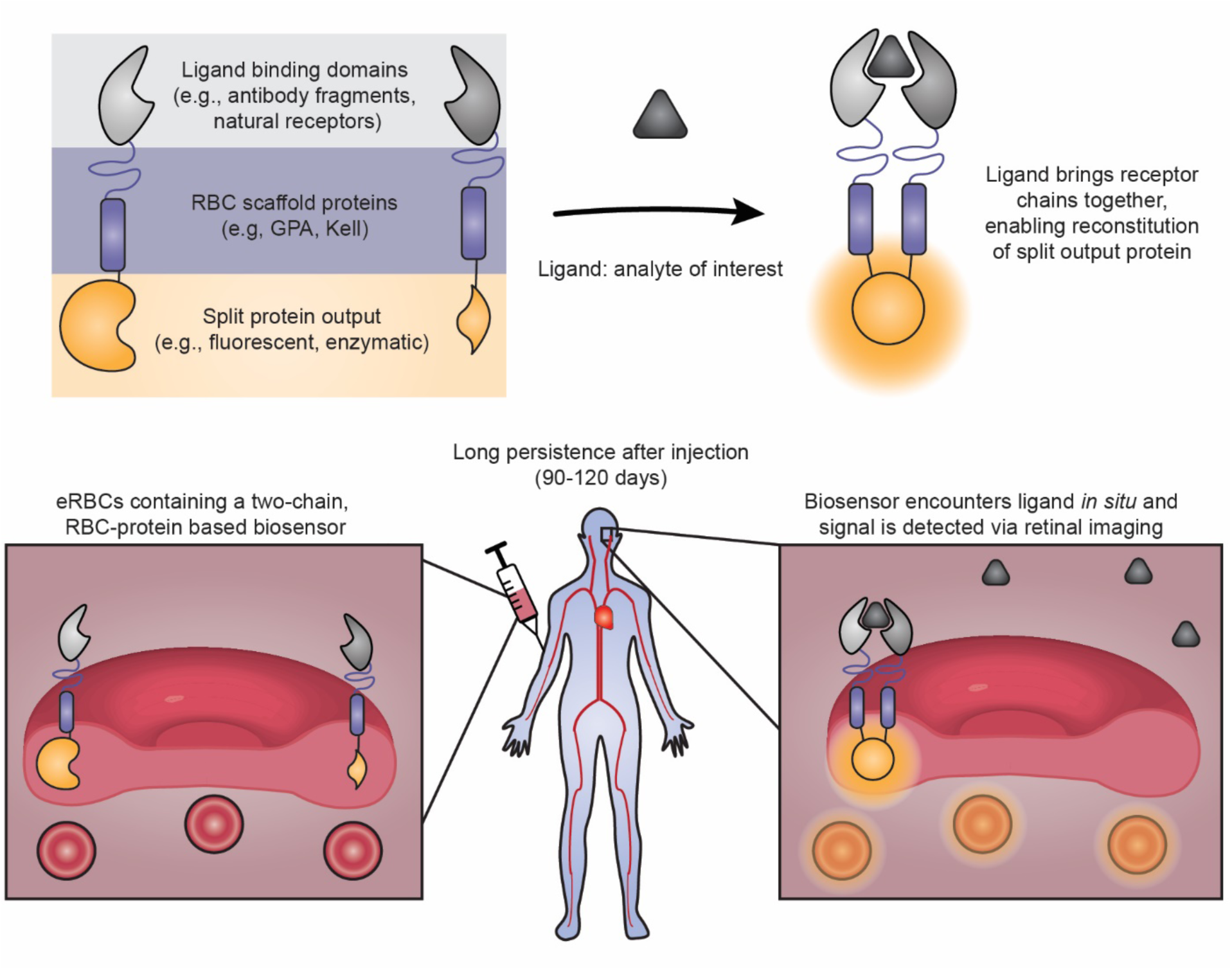
Conceptual vision of engineered red blood cell (eRBC) biosensors to enable non-invasive diagnostics. This cartoon describes a potential strategy through which RBC-based biosensors could enable a new class of capabilities—non-invasive, *in situ* diagnostics based upon long-lived “sentinel” cells. Top, schematic describing the approach used in this study to build an RBC-protein based biosensor. The receptors dimerize upon detecting their ligand, and this proximity promotes reconstitution of an intracellular split protein as the sensor’s output. Bottom, schematic describing a conceptual vision for how RBC-protein based biosensors could be implemented into eRBCs to support *in situ* diagnostics. We hypothesize that RBC precursors may be engineered to express biosensors and then differentiated, and biosensing eRBCs may then be administered to a patient for long-term monitoring (90 – 120 days). This illustration depicts a potential modality for routine, non-invasive monitoring of eRBCs in the bloodstream—retinal imaging to quantify biosensor output.

## Materials and methods

### DNA assembly and plasmid preparation

Plasmids encoding RBC biosensor proteins were assembled using standard molecular biology techniques of PCR and restriction enzyme cloning with Phusion DNA Polymerase (NEB), restriction enzymes (NEB; Thermo Fisher), T4 DNA Ligase (NEB), and Antarctic Phosphatase (NEB). GPA and Kell containing source plasmids were a generous gift of Harvey Lodish, Whitehead Institute, MIT. The NanoLuc, split NanoLuc,^*29*^ and CyOFP1^*30*^ genes were codon optimized for humans, synthesized (Thermo Fisher), and cloned into existing biosensor chains. EBFP2 was sourced from pEBFP2-Nuc, which was a gift from Robert Campbell (Addgene plasmid #14893).^*31*^ DsRed-Express2 was obtained by site directed mutagenesis of pDsRed2-N1, which was a gift from David Schaffer (University of California, Berkeley).^*32*^ Enhanced green fluorescent protein (eGFP)^*33*^, referred to as GFP throughout, was split according to prior reports.^*34, 35*^ The cooperative domains CZ and NZ were from Ghost et al.^*35*^ Firefly luciferase^*36*^ was a gift from Lonnie Shea. Plasmids were transformed into chemically competent TOP10 *E. coli* (Thermo Fisher) and grown at 37°C. Plasmid DNA used for transfection and transduction was prepared using the PEG precipitation method which has been extensively described previously.^*37*^ Plasmid maps cloned for this study are in **Supplementary Data 1**.

### HEK293FT cell culture

The HEK293FT cell line was purchased from ThermoFisher/Life Technologies (RRID: CVCL_6911). Cells were cultured in DMEM (Gibco #31600-091) with 3.7 g/L sodium bicarbonate (Fisher Scientific #S233), 4.5 g/L glucose (1g/L, Gibco #31600-091; 3.5 g/L additional, Sigma #G7021), 10% FBS (Gibco #16140-071), 6 mM L-glutamine (2 mM from Gibco 31600-091 and 4 mM from additional Gibco 25030-081), penicillin (100 U/μL), and streptomycin (100 μg/mL) (Gibco 15140122) in a 37°C incubator with 5% CO_2_. Cells were subcultured using Trypsin-EDTA (Gibco #25300-054).

### G1ER cell culture

The murine G1ER cell line was kindly provided by Drs. Mitchell Weiss and Stuart Orkin.^*38–40*^ Cells were maintained between 1 x 10^*5*^ and 2 x 10^6^ cells/mL by dilution into fresh medium or by centrifuging the cells at 125 g x 5 min and resuspending in fresh medium. G1ER cells were cultured in media containing 500 mL Iscove’s MDM (Gibco, 12440053), 75 mL ES-grade FBS (Gibco, 16141079), 10 mL penicillin/streptomycin stock (Gibco 15140122), 6.2 µL monothioglycerol (MTG) (Sigma, M6145), 100 µL erythropoietin (Procrit, 10,000 U/mL stock), and 50 µL puromycin (InvivoGen, ant-pr; ∼1 µg/mL). Fresh mouse stem cell factor was added when passaging cells at 20 ng / mL with a 1000X dilution (Sigma S9915, resuspended in 0.8X PBS).

### Transfection

Transient transfection of HEK293FT cells was conducted using the calcium phosphate method. Transfections were performed in 24-well plates seeded at 1.5 x 10^5^ cells in 0.5 mL of DMEM media (for functional experiments), in 12-well plates seeded at 3 x 10^5^ cells in 1 mL of DMEM media (for surface staining experiments), or in 6-well plates seeded at 7 x 10^5^ cells in 2 mL of DMEM media (for Western blot experiments). At 6-8 h post seeding, cells were transfected with DNA. All experiments included either a blue fluorescent protein (eBFP2) or a red fluorescent protein (DsRedExpress2) as a control to assess transfection efficiency. Luciferase assays in HEK293FT cells included constitutive Firefly luciferase (FLuc) as a normalization control. Plasmid DNA was diluted in H_2_O, and 2 M CaCl_2_ was added to a final concentration of 0.3 M CaCl_2_. Unless otherwise stated, typical amounts of DNA transfected in a 24-well plate were 500 ng of each receptor (or 500 ng of an empty pcDNA3.1 backbone plasmid, if only one receptor plasmid was transfected) and 250 ng of the eBFP2, dsRedExpress2, or firefly luciferase transfection control plasmid. BRET studies in a 24-well plate were transfected with 600 ng DNA total: 200 ng of each receptor or empty pcDNA3.1 backbone, and 200 ng of eBFP2 plasmid. For larger transfections, DNA doses were scaled with the volume. The DNA mixture was added dropwise to an equal-volume solution of 2x-HEPES-Buffered Saline (280 mM NaCl, 0.5 M HEPES, 1.5 mM Na_2_PO_4_) and gently pipetted up and down four times. After 2.5 min, the solution was mixed vigorously by pipetting ten times. 100 µL of this mixture was added dropwise to each well of a 24-well plate, 200 µL was added dropwise to each well of a 12-well plate, or 400 µL was added dropwise to each well of a 6-well plate. Approximately 12-14 h post-transfection, the medium was aspirated and replaced with fresh medium. For functional assays, fresh medium contained 100 nM rapamycin (Sigma 53123-88-9) or 0.05% DMSO as a control. At 24-30 h post media change, cells were harvested for flow cytometry, luciferase assay, Western blot, or surface staining.

### G1ER lentiviral construction and transduction

MigR1-based retroviral vectors were used to express the biosensors and positive controls (**Figs. 4.3a and 4.6a**). Retrovirus was packaged in Plat-E cells (Cell Biolabs) using the calcium phosphate transfection method. Virus was collected the day after transfection, then passed through 0.45 µm syringe filter and used to transduce G1ER cells through spinoculation. Briefly, approximately 2 x 10^6^ G1ER cells were resuspended in 0.8 to 1 mL medium in the presence of virus, 10 mM HEPES (Thermo Fisher Scientific, 15630-080) and 8 µg/mL polybrene (Millipore, TR-1003-G), and then centrifuged at 2,500 rpm for 90 min at 32 °C. Spinoculation was performed 2 times to improve efficiency. The medium was replaced the next morning, and 48 h after spinoculation, G1ER cells were sorted for GFP expression (NanoLuc biosensors) or CyOFP1 expression (BRET biosensors) with flow cytometry (BD FACSAria II Cell Sorter). Sorted cells were checked for biosensor expression by surface staining with Myc-Tag antibody (Myc-Tag 9B11 Mouse mAb, PE-conjugate, #3739, CST). Sorted, stable G1ER cell lines were expanded and cultured until used for experiments.

### Flow cytometry

For functional GFP reconstitution experiments in HEK293FT cells, cells were harvested for flow cytometry using FACS Buffer (PBS pH 7.4, 2 mM EDTA, 0.1% bovine serum albumin). Harvested cells were spun at 150 x g for 5 min, the FACS buffer was decanted, and fresh FACS buffer was added. Approximately 10^4^ single, live cells from each sample were analyzed using a BD LSRII flow cytometer (Robert H. Lurie Cancer Center Flow Cytometry Core) or a BD LSR Fortessa Special Order Research Product (Robert H. Lurie Cancer Center Flow Cytometry Core). Live, single cells were gated based on scatter, and transfected cells were gated based on the presence of a BFP (recorded in the DAPI channel, with excitation laser 405 nm and 450/50 filter set) or DsRed (recorded in the PE channel, with excitation laser 550 nm and 582/15 filter set) transfection control for all experiments (**Figure S1**). GFP signal was recorded in the FITC channel, with excitation laser 488 nm and 505LP, 530/30 filter set. Data reported in these plots are background subtracted, where the background was the average GFP fluorescence of transfected cells that did not receive full GFP.

For immunolabeling on the cell surface of HEK293FT cells, 10^6^ cells were harvested with FACS buffer and spun at 150 x g for 5 min. FACS buffer was decanted, 50 μL fresh FACS buffer was added, and 10 μL human IgG (Human IgG Isotype Control, ThermoFisher Scientific #02-7102, RRID: AB_2532958, stock concentration 1 mg/mL) was added. Cells were incubated at 4°C for 5 min. 2.5 μL Myc-Tag antibody (Myc-Tag 9B11 Mouse mAb, PE-conjugate, #3739, CST) was added and cells were incubated at 4°C for 30 min. After incubation, cells were washed three times by addition of 1 mL FACS buffer and 150 x g centrifugation at 4°C for 5 min. After the final wash, supernatant was decanted and 3 drops of FACS buffer was added to each sample. Approximately 10^4^ single, live cells from each sample were analyzed using a BD LSRII flow cytometer (Robert H. Lurie Cancer Center Flow Cytometry Core) or a BD LSR Fortessa Special Order Research Product (Robert H. Lurie Cancer Center Flow Cytometry Core). Live, single cells were gated based on scatter and transfected cells were gated based on the presence of a BFP (recorded in the DAPI channel, with excitation laser 405 nm and 450/50 filter set) transfection control for all experiments (**Figure S1**). PE fluorescence (conjugated to Myc-Tag antibody) was recorded in the PE channel, with excitation laser 550 nm and 582/15 filter set.

### NanoLuciferase assays

For functional NanoLuc reconstitution experiments in HEK293FT cells, Nano-Glo Dual-Luciferase Reporter Assay System (Promega N1610) was used. Cells were washed with PBS and harvested with 1X Passive Lysis Buffer (5X, Promega #E1941) diluted in water. 100 µL Passive Lysis Buffer was added to each well of a 24-well plate for cell harvest, incubated for 5 min, and then cell lysates were transferred to 1.7 mL Eppendorf tubes. 80 µL of cell lysate was added to each well of a 96-well white-walled plate (Thermo Fisher Scientific #655906). 80 µL ONE-Glo EX Reagent was then added and the plate was orbitally shaken at 300 rpm for 3 min in a microplate reader (BioTek Synergy H1, v2.04.11). Firefly luminescence was collected using the microplate reader’s luminescence fiber; the gain was adjusted to the positive control well, and the integration time set to 1 s. Next, 80 µL of NanoDLR Stop & Glo Reagent was added to each well. The plate was then orbitally shaken at 600 rpm for 3 min and further incubated with no shaking for an additional 10 min. NanoLuc luminescence was collected using the microplate reader’s luminescence fiber, with gain set to 200 and integration time of 1 s. Each condition was performed in biological triplicate, and the mean NanoLuc signal and mean Firefly signal were calculated for each condition. The NanoLuc signal was divided by the Firefly signal for each condition and reported as normalized luminescence. Error was propagated accordingly.

For functional NanoLuc reconstitution experiments in G1ER cells, the Nano-Glo Luciferase Assay System (Promega N1110) was used. G1ER cells were treated with 100 nM rapamycin or vehicle for 15 min, 1 hr, or 3 hrs, and then pelleted and resuspended in passive lysis buffer. 50 µL of G1ER cell lysates (corresponding to approximately 2.7 x 10^5^ cells) harvested with 1X Passive Lysis Buffer were transferred into white-walled 96-well plates (Thermo Fisher Scientific #655906). GFP fluorescence was collected in the microplate reader (BioTek Synergy H1, v2.04.11) using the GFP filter set with gain set to 35. Next, 50 µL of Nano-Glo Luciferase Assay Reagent was added to each well, and the plate was incubated in the microplate reader with 300 rpm orbital shaking for 3 min. NanoLuc signal was collected using the microplate reader’s luminescence fiber, with gain set to 200 and integration time of 1 s. Each condition was performed in technical triplicate unless otherwise stated, and the mean NanoLuc signal and mean GFP signal were calculated for each condition. The NanoLuc signal was divided by the GFP signal for each condition and reported as normalized luminescence. Error was propagated accordingly.

### Western blotting

For HEK293FT Western blots, cells were washed with PBS before harvest and lysed with 250 μL NP-40 buffer (150 mM NaCl, 50 mM Tris-HCl pH 8.0, 1% Triton X-100) with protease inhibitor cocktail (Pierce/Thermo Fisher #A32953). For G1ER Western blots, cell lysates were harvested with 1X Passive Lysis Buffer (5X, Promega #E1941) diluted in water. Next, NP-40 (HEK293FT) or Passive Lysis Buffer (G1ER) cell lysates were incubated on ice for 30 min and spun at 12,200 rpm at 4°C for 20 min. The supernatant was transferred to a new tube, and the pellet was discarded. A BCA assay (Pierce/Thermo Fisher #23225) was performed to determine protein concentration of each sample so that equal protein concentrations could be loaded per well. 2 to 10 ug of total protein was mixed with Lamelli buffer (final concentration 60 mM Tris-HCL pH 6.8, 10% glycerol, 2% sodium dodecyl sulfate, 100 mM dithiothreitol, and 0.01% bromophenol blue) and incubated at 70°C for 10 min. Samples were loaded into a 4-15% Mini-PROTEAN TGX Precast Protein gel (Bio-Rad) and run at 50 V for 10 min followed by 100 V for 1-1.5 h. Wet transfer was performed onto an Immuno-Blot PVDF membrane (Bio-Rad) for 45 min at 100 V. Protein transfer was confirmed with Ponceau-S stain before the membrane was blocked with 5% milk in TBST (pH 7.6, 50 nM Tris, 150 mM NaCl, 0.1% Tween) while rocking for 4 h at 4°C or 1 h at room temperature (20-25°C). Membranes were then incubated with primary antibody (Anti-Myc tag antibody, mouse monoclonal [9E10], Ab32, Abcam), diluted 1:1000 in 5% milk in TBST at 4°C overnight while rocking. Primary antibody was decanted, and the membrane was washed 3 times with TBST for 5 min each. Secondary antibody (HRP-anti-Mouse, CST 7076, RRID: AB_330924) was diluted 1:3000 in 5% milk in TBST and applied for 1 h at room temperature while rocking. The membrane was washed 3 times with TBST for 5 min each and then the membrane was incubated with Clarity Western ECL Substrate (Bio-Rad) for 2 min before exposure to film and development in a darkroom.

### HEK293FT BRET imaging

For BRET assays with HEK293FT cells, cells were washed with PBS before harvest and harvested with 1X Passive Lysis Buffer (5X, Promega #E1941) diluted in water. 200 µL Passive Lysis Buffer was added to each well of the 12-well plate and incubated for 5 min; cell lysates were then transferred to 1.7 mL Eppendorf tubes. 50 µL of cell lysate was added to each well of a 96-well black-walled, clear bottom plate, and 50 μL of Nano-Glo Luciferase Assay System Reagent (Promega N1110) to added to each well and mixed via pipetting. For conditions with less than 50 μL of cell lysate, extra passive lysis buffer was added to bring the volume of each condition to 50 μL before adding the Nano-Glo reagent. The plate was wrapped in foil, incubated in the dark for 3 min, and then imaged in an in vivo imaging system (IVIS) Spectrum (PerkinElmer; blocked excitation filter and open emission filter). To mimic tissue—which would absorb much of the intense blue-spectrum NanoLuc emission but would allow more of CyOFP1’s red shifted light to reach the detector—pieces of ham (Krakus sliced polish ham with natural juices, 3 mm thick, Mariano’s), were placed on top of the plate during IVIS imaging. Images were analyzed and quantified with Living Image Software (v4.5.2.19424).

### G1ER BRET imaging

For functional BRET experiments in G1ER cells, the Nano-Glo Luciferase Assay System (Promega N1110) was generally used. G1ERs were pelleted at 125 g x 5 min at 4°C, resuspended in fresh medium, and added to a 96 well, black-walled plate in 100-120 µL. Exact cell numbers per well can be found in the relevant captions; biosensor (TD146/TD147) cells were generally plated between 5 x 10^5^ – 5 x 10^6^ cells per well, and positive control (TD142) cells were generally plated between 750 – 3000 cells per well in triplicate. Rapamycin (or equivalent vehicle) was added at a final concentration of 100 nM in 1-1.2 µL and mixed via pipetting. Plates were incubated at 37°C for approximately 30 min and allowed to cool to room temperature. Nano-Glo was then added 1:1 with the original volume of cell suspension in the well (approximately 1.96 µL or 10.3 nanomoles substrate/well)^*41*^ and mixed via pipetting. When used, fluorofurimazine (FFZ, Promega, CS320501 (Nano-Glo® In Vivo Substrate, 4.6 µmoles) was resuspended in 525 µL of PBS. FFZ was further diluted in PBS, added to the well plate in 100 µL (approximately 10.3 nanomoles/well), and mixed via pipetting. Plates were incubated in the dark for 3 min before imaging on an IVIS spectrum using an open emission filter and a blocked excitation filter. Ham (Krakus, sliced polish ham, 3 mm) were placed over the well plate to simulate tissue during imaging. Images were analyzed and quantified with Living Image Software (v4.5.2.19424).

### G1ER differentiation studies

G1ER cells were harvested, pelleted at 125 g x 5 m at 4°C, and resuspended at 1 x 10^4^ cells/mL in fresh G1ER medium. 100 uL of cell suspension was added per well in a 96-well plate. 100 uL of G1ER medium containing either no additional treatment, 100 nM β-estradiol (MilliporeSigma, E8875-250MG), or an equivalent amount of ethanol vehicle were added to wells containing cells and mixed. An initial 5 mM stock of β-estradiol was made by dissolving in 100% ethanol, and intermediate stocks were made by further diluting in ethanol to 100 uM; aliquots were stored at -20°C and diluted 1000X upon addition to cells. Cells were incubated at 37°C, 5% CO_2_ for two days. Cells were then imaged on a Keyence BZ-x800 microscope (BZ Series Application software v01.01.00.17) with a PlanApo 4X (NA 0.2) objective. Wells were then mixed and two aliquots (one to be treated with rapamycin and one for vehicle) of 59 uL of cell suspension was transferred to a white walled, 96-well plate and 20 uL to a microcentrifuge tube. Rapamycin at 100 uM in 50% DMSO/50% nuclease free (NF) water was diluted to 6 uM with NF water, and 1 uL of the rapamycin or vehicle was added to cells in the 96-well plate. The plate was spun at 500 g x 1 min at 4°C to remove bubbles, and then incubated at 37°C for 30 min. Nano-Glo reagent was diluted 51X in Nano-Glo buffer, 60 uL was added per well, and samples were pipetted to mix. The plate was shaken at 300 rpm (orbital) for 3 min and BRET biosensor performance was recorded using the plate reader’s luminescence filter (BioTek Synergy H1, v2.04.11) with an integration time of 1 s, full emission light, top optics, and a gain of 200. The cell aliquot put into a microcentrifuge tube was mixed 1:1 with trypan blue and counted on a manual hemocytometer. Luminescent values from the plate reader were normalized to the matched cell count.

### G1ER kinetic studies

G1ER cells were harvested, pelleted at 125 g x 5 min at 4°C, and plated into a white-walled 96-well plate in fresh G1ER medium at 10^5^ cells/well in 80 µL. Cells were treated with 100 nM rapamycin or vehicle in 2 µL for 60, 30, 15, or zero min at 37°C before addition of Nano-Glo substrate (Promega N1110) in 80 µL. The well plate was then shaken at 300 rpm for 3 min before being read using a Synergy H1 plate reader (Aglient, v2.04.11). Luminescent values were recorded using a gain of 200 and an integration time of 1 s.

### In vivo mouse experiments

Northwestern University’s Institutional Animal Care and Use Committee approved all animal protocols. 8–10 week-old, male BALB/c mice, weighing 25-30 g, were obtained from The Jackson Laboratory. Mice trunks were shaved prior to the study. The day of the study, 10 x 10^6^ BRET biosensor G1ER cells suspended in 100 µL PBS (or a PBS control) were injected via tail vein. 20 mg / kg rapamycin (Selleck Chemicals) or equivalent vehicle (10% DMSO (Protide Pharmaceuticals), 40% PEG-300 (Spectrum Chemical, PO108), 5% Tween 80 (Sigma), 45% saline) were injected intraperitoneally (IP) at 10 mL/kg. Animals were then anesthetized with 1-3% isoflurane, and a baseline image (t = 0) was captured via an IVIS Spectrum (PerkinElmer). Animals were removed from anesthesia after the image. 30 min after rapamycin or vehicle injection, animals were again anesthetized, injected IP with FFZ at 44 µmole/kg in PBS, and imaged at approximately 2, 7, and 12 minutes after FFZ injection. For all *in vivo* images, the camera was in luminescent mode (excitation filter closed and emission filter open), with medium binning, and the exposure time was set to 60 s. All images were taken from the dorsal perspective. Black plastic separators were placed between animals to prevent signal bleed over from one mouse to the next. Animals referred to with an identifier “L” (Ligand) were administered biosensor cells, rapamycin, and FFZ; animals with an identifier “V” (Vehicle) were administered biosensor cells, vehicle, and FFZ; animals with an identifier “S” (Substrate) were administered PBS, rapamycin, and FFZ. To quantify luminescent flux from each animal, a region of interest (ROI) was manually drawn for each image as described in **Figure S14**. Image capture and analysis were performed using Living Image software v4.5.2.19424. A one-tailed t-test was performed in GraphPad; this test was justified because of a priori knowledge from the *in vitro* studies suggesting that rapamycin-treated, BRET biosensing G1ERs have a higher ON state than vehicle-treated cells.

## Results

### Evaluating eRBC biosensor feasibility

As a first step toward developing an eRBC biosensor, we explored designs which use two native RBC membrane proteins—Glycophorin-A (GPA) or Kell—as biosensor scaffolds (**Figure 2a)**. GPA and Kell are present on both precursor cells and mature erythrocytes, suggesting that precursor cells engineered with GPA and Kell-based receptors might retain the biosensors after erythrocyte maturation.*^2,^* ^*28, 42*^ Employing a synthetic receptor design widely used by us and others,^*27, 37, 43–45*^ we elected to pursue a mechanism whereby two receptor chains dimerize upon extracellular ligand binding to promote intracellular reconstitution of a split protein as the output signal (**Figure 1**, **2a)**. This topology was motivated by the expectation that many analytes of interests (e.g., cytokines, toxins) may be restricted to the extracellular compartment, while retention of the biosensor output domain in the intracellular compartment shields it from proteases and host antibodies. We selected rapamycin as our model ligand and the rapamycin binding domains FKBP (FK506-binding protein) and FRB (FKBP rapamycin-binding)^*46*^ as model exterior sensing domains because these well-characterized modules have been widely used to study protein-protein interactions.^*43, 47–52*^ This approach has generally proven useful for driving biosensor development before subsequent adaptation to sensing other ligands of interest.^*27*^ As the initial output module, we employed split eGFP—wherein eGFP has been split into two fragments: GFP-Large (GFPL, consisting of amino acids 1-157) and GFP-Small (GFPS, consisting of amino acids 158-238).^*35, 53*^ Neither half of GFP fluoresces independently, but GFP fluorescence is restored upon reconstitution of the two fragments. We anticipated that some geometric flexibility may be required to enable the split GFP parts to reconstitute when fused to their scaffolds. Therefore, we included a small (0, 6, or 10 amino acid) unstructured linker between the scaffold and the split GFP. These various choices were used to generate a pilot library of biosensor candidates in order to explore this design space.

**Figure 2.**
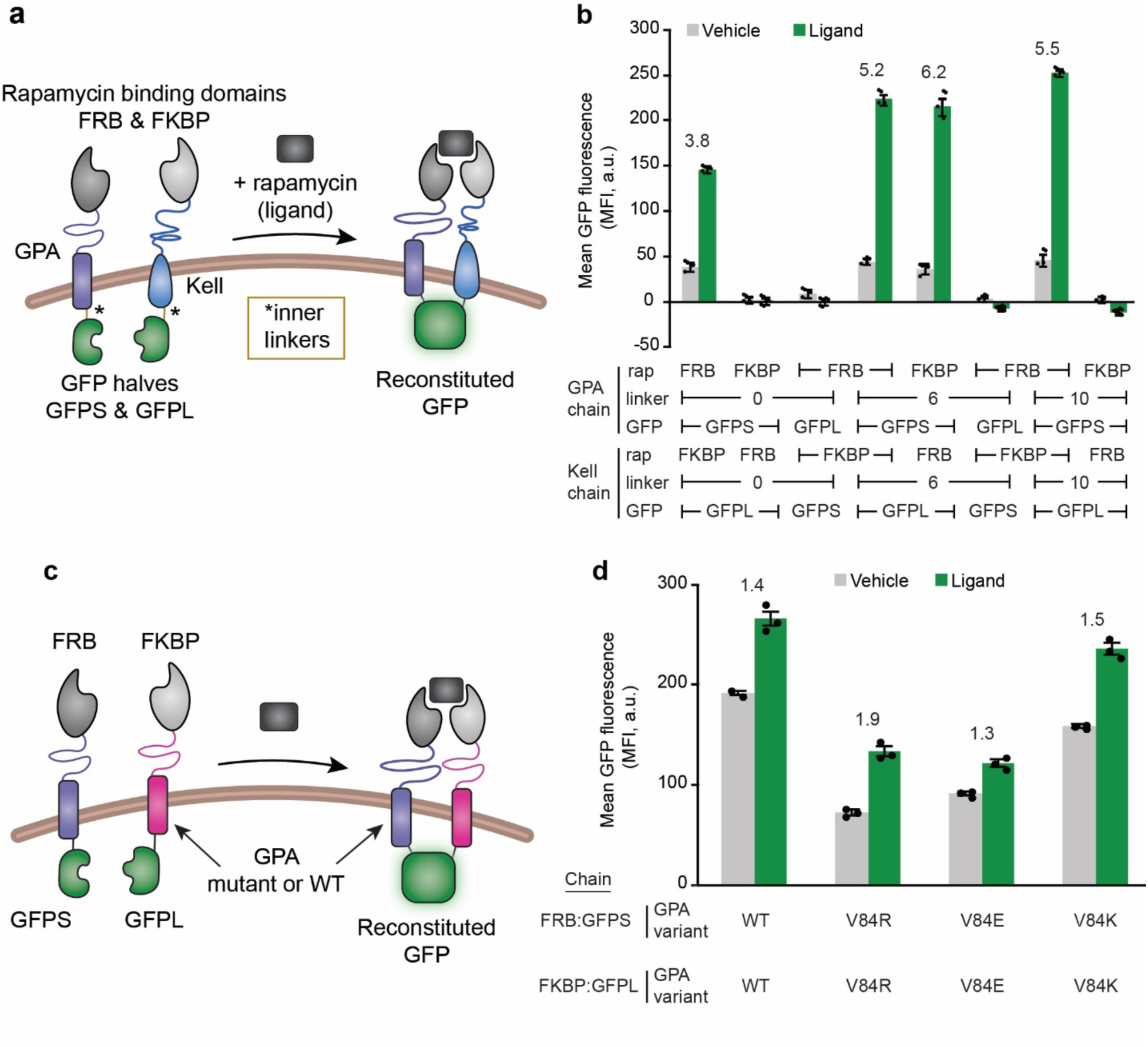
Performance of eRBC biosensors change as a function of scaffold and linker length choices. **(a)** This schematic depicts the design for an eRBC biosensor built on two native RBC membrane protein: GPA and Kell. Rapamycin binding domains, FKBP and FRB, are tethered to the extracellular domains of GPA and Kell, and two halves of GFP, GFPS and GFPL, are tethered to the intracellular domains of GPA and Kell via a small inner linker. Upon the addition of rapamycin, the two chains come into close proximity which drives the two halves of GFP to reconstitute and fluoresce. All studies in this figure were performed in HEK293FT cells. **(b)** Functional analysis of the first generation of Kell and GPA-based biosensors generated by varying choice of rapamycin binding domains (FRB, FKBP) and GFP halves (GFPS, GFPL) between Kell and GPA chains that contained a 0, 6, or 10 amino acid (glycine-serine) inner linker. Flow cytometry was used to measure GFP reconstitution. Four of the eight combinations exhibited a substantial increase in GFP fluorescence upon ligand treatment. **(c)** Schematic depicting the design of GPA-based biosensors where the scaffold can be WT or mutant GPA. **(d)** Functional analysis of GPA-based biosensors. Flow cytometry was used to measure GFP reconstitution. All V84 mutations to the transmembrane domain of GPA decreased both the background GFP reconstitution and the ligand-induced GFP reconstitution when compared to the WT GPA. The symbols represent biological replicates, the bars represent the mean, the error bars represent standard deviation, and the numbers above the bars are the fold induction (signal with ligand divided by signal with vehicle) for biosensors with a fold induction greater than 0. Data in this figure are representative of at least two experiments. MFI, mean fluorescence intensity.

We next evaluated our pilot library of biosensor candidates, with a focus on evaluating failure modes and fundamental feasibility. First, each biosensor chain was expressed by transient transfection in HEK293FT cells, and surface expression of all constructs was confirmed by flow cytometry (**Figure S1, S2**). Next, cells were transfected to express various pairs of biosensor chains, treated with rapamycin ligand or vehicle (DMSO) control, and interrogated for GFP fluorescence via flow cytometry (**Figure 2b**). We observed ligand-inducible GFP reconstitution for four of the eight combinations tested. This high degree of success in this pilot evaluation demonstrated the fundamental feasibility of engineering biosensor chains on RBC membrane proteins.

Towards improving biosensor performance, we next explored strategies for improving the biosensor ON state by modulating biosensor design. The magnitude of GFP signal in the initial biosensor panel was relatively weak (for example, unlike a full GFP tethered to a GPA or Kell, reconstituted GFP was not observable by fluorescence microscopy), and therefore we hypothesized that GFP reconstitution may be limited by geometric constraints. We observed that three of the inducible pairs from the **Figure 2b** had the same chain architecture: FRB and GFPS domains on the GPA chain, and FKBP and GFPL domains on the Kell chain. All exhibited increasing GFP signal for increasing length of inner linkers. Therefore, we built a second set of biosensors based on this promising architecture and further extended the length of the inner linkers to potentially alleviate geometric constraints or steric hindrance limiting split GFP reconstitution. Ligand-induced reconstituted GFP signal increased when the inner linker domain was extended from 10 amino acids to 20 amino acids (**Figure S3a**), but this improvement in ON state came at the expense of increased background (ligand-independent) GFP reconstitution, and the ON state was still well below the signal magnitude of full GFP control chains (**Figure S3b**). We hypothesized that GFP reconstitution might be increased by genetically fusing engineered leucine zipper domains to the receptors’ inner linkers, which interact with one another at a range of affinities and thus may modulate receptor-receptor affinity.^*35, 54*^ We found that these designs indeed enhanced split GFP reconstitution, but this effect was independent of rapamycin treatment and ultimately diminished the biosensor’s inducibility for nearly all cases (**Figure S4a**). When compared to control chains containing full GFP, the reconstituted GFP signal magnitude was still substantially lower (**Figure S4b**). Together, these results suggest that biosensor performance can be modulated by tuning the inner linker architecture, but that GFP reconstitution with the current biosensor geometry was still limited.

We next explored how changing the RBC-protein scaffold would affect biosensor geometry and overall biosensor performance. This effort was motivated by the observation that GPA and Kell have significant differences in size (72 aa extracellular domain and 665 aa extracellular domain, respectively) and structure (type I vs type II transmembrane protein, respectively).^*24, 55, 56*^ We hypothesized that Kell’s large extracellular region might allow Kell-based biosensors to bind rapamycin without bringing the intracellular split GFP close enough to a partner GPA-based receptor to reconstitute GFP. Because Kell is retained on mature erythrocytes at a lower copy number than is GPA (3 – 7 x 10^3^ copies of Kell per RBC compared to 500 – 1000 x 10^3^ copies of GPA per RBC^*55*^), we next pursued a biosensor architecture where both chains used a GPA scaffold. Biosensor performance data show that the GPA-only biosensors had a much higher maximum GFP signal than did the GPA-Kell biosensors; however, the background level was also higher for GPA-only designs (**Figure S3c**). The substantially higher signal may indicate that GFP reconstitution is more favorable, or less constrained, when using the GPA-only architecture compared to the GPA-Kell biosensors, at the expense of lower fold induction. Such observations motivated a refinement of this GPA-only design.

Since high background signal from GPA-only biosensors may derive from GPA homodimerization, we next explored strategies for mutating GPA to reduce background. GPA homodimerization is mediated by seven amino acids in the transmembrane domain (TMD),^*57*^ and mutating one or more of these residues decreases the propensity for homodimerization.^*58, 59*^ Since dimerization-deficient mutant GPA variants are still displayed at the RBC surface and, like wild-type (WT) GPA, facilitate Band 3 translocation^*60*^ (one indicator of preserved GPA structural integrity), we anticipated that mutating GPA’s TMD would be unlikely to ablate our biosensor’s display on an eRBC’s surface. We generated biosensors with three mutations to one TMD residue (valine 84) that decreases GPA dimerization,^*57*^ and expressed these biosensors in HEK293FT cells (**Figure S5a**). Via western blot, we found that all constructs were expressed, and we observed dimerized GPA chains for biosensors employing WT GPA but not the mutant GPA chains (**Figure S5b**). All variants were surface expressed in HEK293FT cells (**Figure S5c**). In functional evaluations, mutant GPA biosensors exhibited decreased background GFP reconstitution as well as decreased ligand-induced GFP reconstitution (**Figure 2d**). We speculate that the lower overall expression of the mutant GPA chains (**Figure S5b**) may contribute to the lower GFP reconstitution signal, especially for poorly expressed chains (e.g., FRB-V84R-GFPS). We observed no improvement in ligand-induced GFP reconstitution or fold induction when we extended the inner linkers between the V84R GPA and GFP domain or added outer linkers between the rapamycin binding domain and the V84R GPA compared to our base case (**Figure S6**). Together, these data demonstrate that functional RBC-protein based biosensors can be built with a number of different architectures, and that choice of scaffold and linker can help to tune receptor performance, while also motivating the exploration of additional choices—beyond the scaffold—to further improve performance.

### Adaptation of GPA-based biosensors to yield enzymatic bioluminescent output

Having explored design handles hypothesized to affect biosensor performance through altering receptor geometry, we next investigated changing the output from GFP reconstitution to luciferase reconstitution. We anticipated that because an enzymatic reporter amplifies signal output, this pivot could improve receptor performance. High output may be particularly important for ultimate deployment as a diagnostic, and adaptation of our biosensors to yield an enzymatic output—in general—may be useful for validating the potential to employ output modalities beyond the scope of those contemplated here. We chose to use a well characterized version of split NanoLuc termed NanoBiT,^*29, 61, 62*^ and we implemented this system by replacing the GFPS and GFPL fragments on the biosensor interior with the 11S and 114 fragments of the split NanoLuc system (**Figure 3a**). We built a panel of biosensor chains by varying the rapamycin binding domain (FRB or FKBP), GPA scaffold (WT or V84R mutant, termed mutant GPA), and NanoLuc fragment (114 or 11S), and we evaluated biosensor performance via transient transfection of HEK293FT cells. We observed ligand-inducible NanoLuc reconstitution for all of the biosensor chain pairs tested (**Figure 3b**). The magnitude of ligand-induced luminescence was higher for the WT GPA biosensors than for the biosensors containing mutant GPA; this pattern concords with initial observations with split GFP (**Figure 2c**). These data demonstrate that our biosensor strategy can be employed to regulate reconstitution of an enzyme and produce ligand-induced luminescence. We carried forward the receptor pair that produced higher ligand-induced luminescence—FRB and 114 domains on one chain and the FKBP and 11S domains on the second chain—for further development.

**Figure 3.**
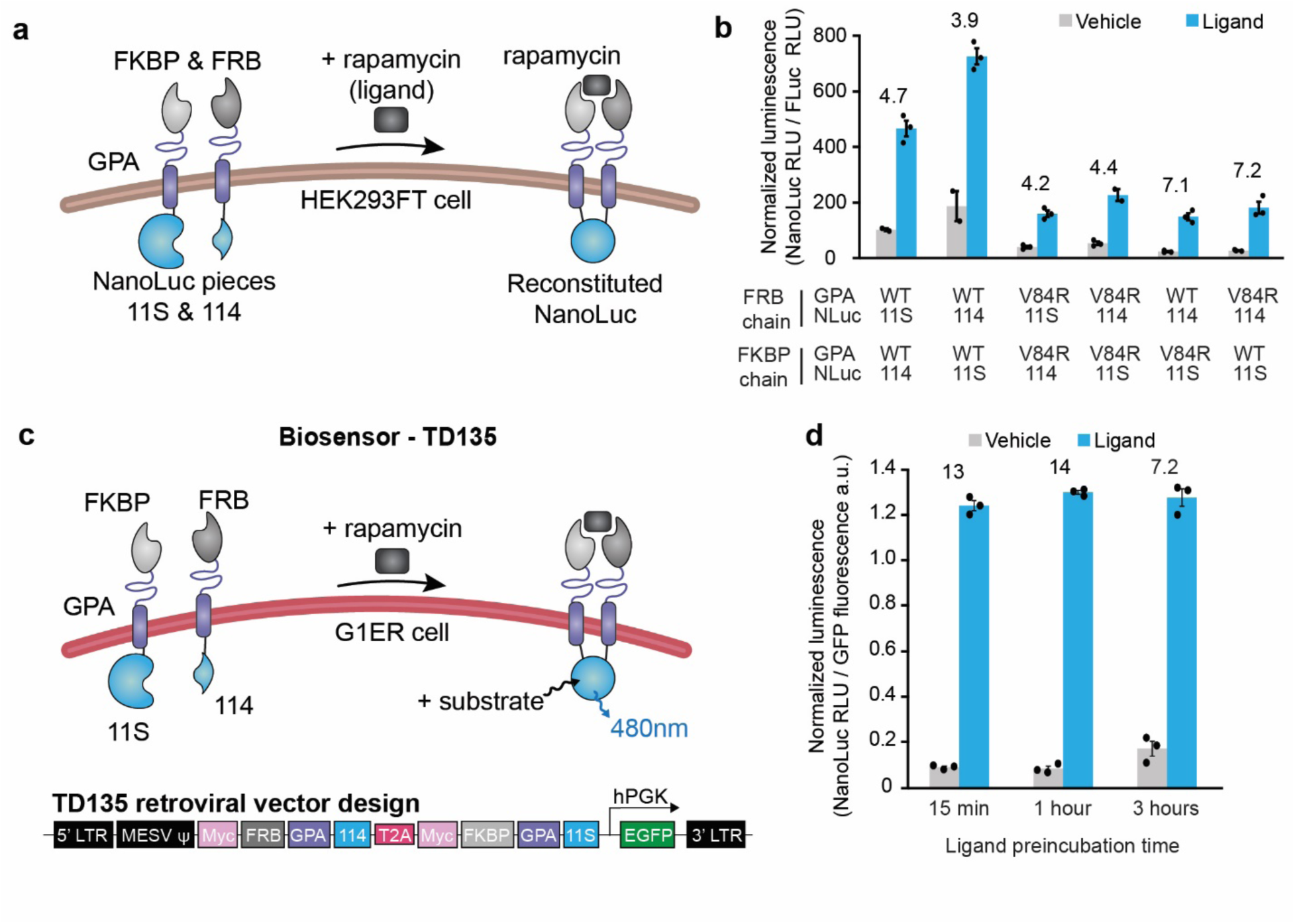
Adaptation of GPA-based biosensors to yield bioluminescent output and validation in erythroid precursor cells. **(a)** This schematic depicts the design for a GPA-based biosensor with a luciferase output. NanoLuc was split into two fragments, 11S and 114, for the output domain. Rapamycin binding domains FKBP and FRB were tethered to the extracellular domains of GPA, and NanoLuc fragments 11S and 114 were tethered to the intracellular domains of GPA. Ligand-induced reconstitution of NanoLuc yields luminescence upon the addition of substrate, furimazine. **(b)** Functional evaluation of GPA-NanoLuc biosensors expressed by transient transfection in HEK293FT cells. Biosensor variants were constructed by swapping NanoLuc fragments 11S and 114 between FRB and FKBP chains and evaluating WT GPA and V84R GPA. Reconstituted NanoLuc luminescence was measured with a plate reader and normalized to the signal from co-transfected Firefly luciferase (Fluc) and reported as normalized luminescence. All architecture combinations tested show increased luminescence with ligand treatment. **(c)** Schematic of biosensor TD135 evaluated in G1ER cells. The two receptor chains in TD135 are separated by a T2A peptide in the retroviral vector, and the hPGK-GFP is used for identifying and sorting transduced cells. **(d)** Functional analysis of GPA-NanoLuc biosensor chains in mouse G1ER cells as a function of ligand exposure time. Reconstituted NanoLuc luminescence was measured with a plate reader after addition of furimazine substrate and normalized to GFP fluorescence (expressed from retroviral vector). Throughout the figure, the bars represent the mean, the error bars represent SEM, and the numbers above the bars are the fold induction (signal with ligand divided by signal with vehicle). The symbols represent biological replicates for **b** and technical replicates for **d**. Data are representative of two experiments for panel **b** and one experiment for panel **d.**

### Validation of GPA-based biosensors in erythroid precursor cells

We next investigated whether our GPA/NanoLuc-based biosensor could be implemented in an RBC-precursor cell. We chose to use G1ER cells which are an engineered murine erythroid precursor cell line, most resembling erythroblasts, that is capable of undergoing erythroid differentiation as early as 24 h after the addition of β-estradiol.^*38–40*^ Retroviral vectors were built encoding the biosensor chains separated by a T2A “self-cleaving” peptide and a GFP gene driven by an hPGK promoter to enable cell sorting (**Figure S7a**). Four G1ER cells lines were created that expressed FRB/NanoLuc 114 on one biosensor chain and FKBP/NanoLuc 11S on the second chain. Cell lines differed by the choice of mutant or WT GPA as the scaffold: TD135 (WT GPA on both chains), TD136 (mutant GPA on both chains), TD137 (WT GPA on FRB/114 chain, mutant GPA on FKBP/11S chain), TD138 (mutant GPA on FRB/114 chain, WT GPA on FKBP/11S chain). These cell lines were sorted for GFP expression via flow cytometry to select for cells transduced with the retroviral construct, and surface expression of the biosensor chains was evaluated. All four constructs were expressed on the cell surface, and the construct with mutant GPA on both chains (TD136) showed the lowest amount of surface expression (**Figure S7b**). Next, we investigated biosensor performance in G1ER cells. In a pilot study, the biosensor with two WT GPA chains (TD135) exhibited detectable and inducible NanoLuc luminescence; the other three constructs (TD136, TD137, TD138) did not confer detectable NanoLuc luminescence (**Fig S7c**), and thus TD135 was retained for detailed characterization. To evaluate how quickly the biosensor can be activated, we performed a time course study and observed that maximum signal had already occurred within 15 min of exposure to ligand, and the output was steady up to 3 h (**Figure 3c,d**). These data demonstrate the GPA-based biosensors prototyped in HEK293FT cells retain functionality when implemented in RBC-like cells, validating the overall engineering strategy employed here.

### Adaptation of GPA-based biosensors to yield in vivo-optimized output

Toward the goal of validating our biosensors *in vivo*, we next explored whether biosensor output could be refined to incorporate a modality that is well-suited to that application. While many features of NanoLuc make it an attractive output for the RBC protein-based biosensor—small size, well characterized protein split, and bright signal (∼150x brighter than Fluc or Renilla luciferase [Rluc])^*63*^—the blue shifted output (NanoLuc: 460 nm, compared to Rluc: 480 nm and Fluc: 565 nm)^*64*^ does not penetrate tissue well and may impair detection *in vivo*.^*65*^ To address this potential barrier, we explored the use of bioluminescence resonance energy transfer (BRET) for biosensor output, whereby a longer-wave fluorescent protein acceptor is excited, in the presence of substrate, by a bioluminescent luciferase donor that normally emits shorter wavelength light, such as NanoLuc. BRET occurs naturally in jellyfish and has been employed in literature to study protein-protein interactions.^*66*^ Many BRET systems have been developed using NanoLuc as the luciferase donor and incorporating a range of fluorescent protein acceptors.^*63, 67–69*^ Biosensor emission of light with wavelength greater than 600 nm is preferred for *in vivo* imaging because, compared to bluer photons, red photons exhibit longer mean free paths, enabling deeper imaging in tissue.^*70*^ One system, named Antares, utilizes NanoLuc and CyOFP1 (a bright, engineered, orange-red FP that is excitable by cyan light), which shifts the emission from 460 nm to over 600 nm, and in one study, Antares improved *in vivo* detection of reporter systems 84x over NanoLuc alone.^*30*^ For these reasons we selected Antares as an output for our biosensors.

We prototyped a series of GPA-based biosensors to evaluate the feasibility of incorporating Antares-based BRET as an output. We first explored whether BRET could occur while fused to a membrane with our GPA chain architecture. We built a BRET positive control, consisting of an outer FRB rapamycin binding domain, a WT GPA scaffold, a full NanoLuc on the interior, and CyOFP1 tethered to the C-terminus of the NanoLuc (**Figure 4a, left**). We compared the BRET positive control construct to the previous NanoLuc positive control construct (**Figure 4a, right**), via transient transfection of HEK293FT cells. Pieces of thin sliced (3 mm thick) ham were placed on top of a plate containing transfected cell lysates during imaging to mimic tissue (i.e., to mimic the tissue barrier experienced with imaging in mice).^*71*^ The BRET positive control chain was brighter than the NanoLuc positive control at all densities of cell lysate tested through 3 mm ham (**Figure 4b**). The BRET positive signal persisted through multiple layers of tissue (up to 15 mm of ham) at the cell densities tested, whereas the NanoLuc positive signal was minimally detectable past 6 mm of tissue (**Figure S8a-b**). Next, we investigated linking split NanoLuc reconstitution to the BRET output by adding the CyOFP1 domain onto the C-terminal fragment (114) of split NanoLuc. We hypothesized that this position of the CyOFP1 would be least likely to interfere with the reconstitution and folding of NanoLuc because it is attached to the C-terminus of the reconstituted NanoLuc construct rather than in the middle of the two fragments. We also tested different orientations of the chains and multiple linkers (also tested in the development of Antares)^*30*^ between the 114 and CyOFP1 domains because we suspected that particular configurations and geometries might enable different propensities of NanoLuc reconstitution and thus different amounts of BRET signal (**Figure 4c**). In transient transfection of HEK293FT cells, ligand-inducible BRET signal was observed from all biosensor constructs tested (**Figure 4d**). Rapamycin-induced BRET biosensors exhibited detectable signal through many layers of tissue mimetic where NanoLuc biosensor signal was no longer visible (**Figure S8b**). The biosensor with FRB and 114 domains on one chain and FKBP and 11S domains on the second chain, with a two-residue arginine-histidine amino acid linker between 114 and CyOFP1 proteins (orientation 1 and linker D), showed high BRET signal with low background through many layers of tissue mimetic; therefore, this construct was chosen for further evaluation. Together, these observations demonstrate that: (i) the BRET output mechanism is feasible when tethered to full NanoLuc on a membrane; (ii) reconstituted split NanoLuc is amenable to use for BRET; and (iii) the Antares BRET output greatly improves signal penetrance through tissue mimetic.

**Figure 4.**
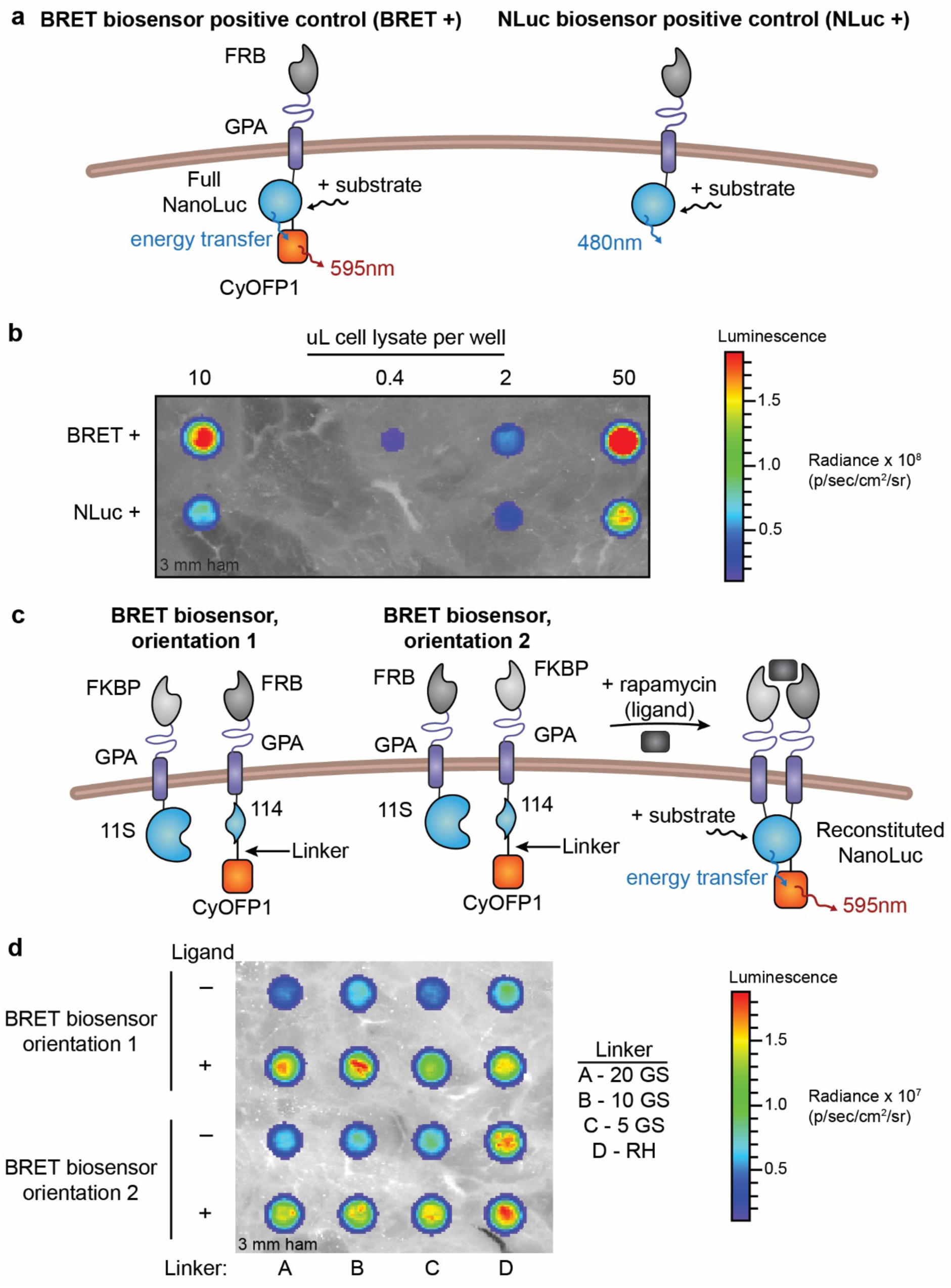
GPA-based biosensors adapted to yield bioluminescence resonance energy transfer (BRET) output improve detection through tissue mimetic. **(a)** This schematic depicts the strategy for modifying biosensor output via BRET to shift the emitted light from 460 nm to 595 nm to ultimately improve *in vivo* detection. The WT GPA-NanoLuc positive control chain (“NLuc+, right) was altered to include the fluorescent protein CyOFP1 on the interior of the chain (“BRET+”, left). When the substrate furimazine is added, NanoLuc excites the CyOFP1 fluorescent protein, resulting in emission of 595 nm light. **(b)** Positive control (BRET +) signal detection in transiently transfected HEK293FT cells imaged with an in vivo imaging system (IVIS) and one piece of ham. Signal detected for BRET+ construct was higher than NanoLuc+ construct signal for all cell densities. **Figure S9b** shows that with increasing thickness of ham, the NanoLuc+ signal was no longer detectable, but the BRET+ signal penetrated up to 15 mm of ham. Data shown are representative of two experiments. **(c)** This schematic illustrates biosensor design wherein split NanoLuc reconstitution drives BRET signal. The CyOFP1 fluorescent protein was fused to the 114 fragment of split NanoLuc (C-terminal fragment) with a linker. Upon ligand treatment and in the presence of NanoLuc’s substrate, the two chains associate, drive NanoLuc reconstitution, and excite CyOFP1 to emit red-shifted, 595 nm light. **(d)** BRET biosensor signal detection in HEK293FT cells with ligand treatment. Two orientations of biosensor parts and four linkers between the 114 NanoLuc domain and CyOFP1 fluorescent protein were examined. GS refers to glycine and serine-containing linkers, and RH is a two amino acid arginine-histidine linker. Signal was detected for all constructs, with higher signal detected for samples treated with ligand. Orientation 1, linker D was chosen for further analysis. **Figure S9c** shows that the biosensor signal was detectable up to about 15 mm of ham. This experiment was performed once.

### Validation of GPA-based BRET biosensor in G1ER cells

Having developed a functional GPA-based BRET biosensor, we next evaluated its use in G1ER cells. Retroviral vectors were constructed to express a BRET positive control (TD142) or biosensor constructs (TD146: chain1-T2A-chain2, TD147: chain2-T2A-chain1), varying the chain orientation with the T2A peptide (**Figure 5a, S9a**). We transduced G1ERs with these vectors and sorted the cells for CyOFP1 fluorescence. Biosensor expression was confirmed by Western blot (**Figure S9b**), and all three stable cell lines showed CyOFP1 expression and stained positive for surface expression of the biosensor chains (**Figure 9c**). Both TD146 and TD147 exhibited ligand-induced signaling (**Figure S9d,e**), although the background and signal for TD146 were nearly two orders of magnitude greater than for TD147. We elected to focus on TD146 (termed BRET biosensor, hereafter) because we anticipated higher signal being an important feature for *in vivo* studies. The BRET biosensor exhibited up to 15 fold induction with ligand addition and could be detected using an IVIS through 18 mm of tissue mimetic (**Figure 5b,c, S10).** Biosensor induction kinetics were rapid—no pre-incubation with rapamycin was required to achieve maximum luminescent signal (**Figure 5d**). These observations confirm BRET biosensor function in G1ER cells.

**Figure 5.**
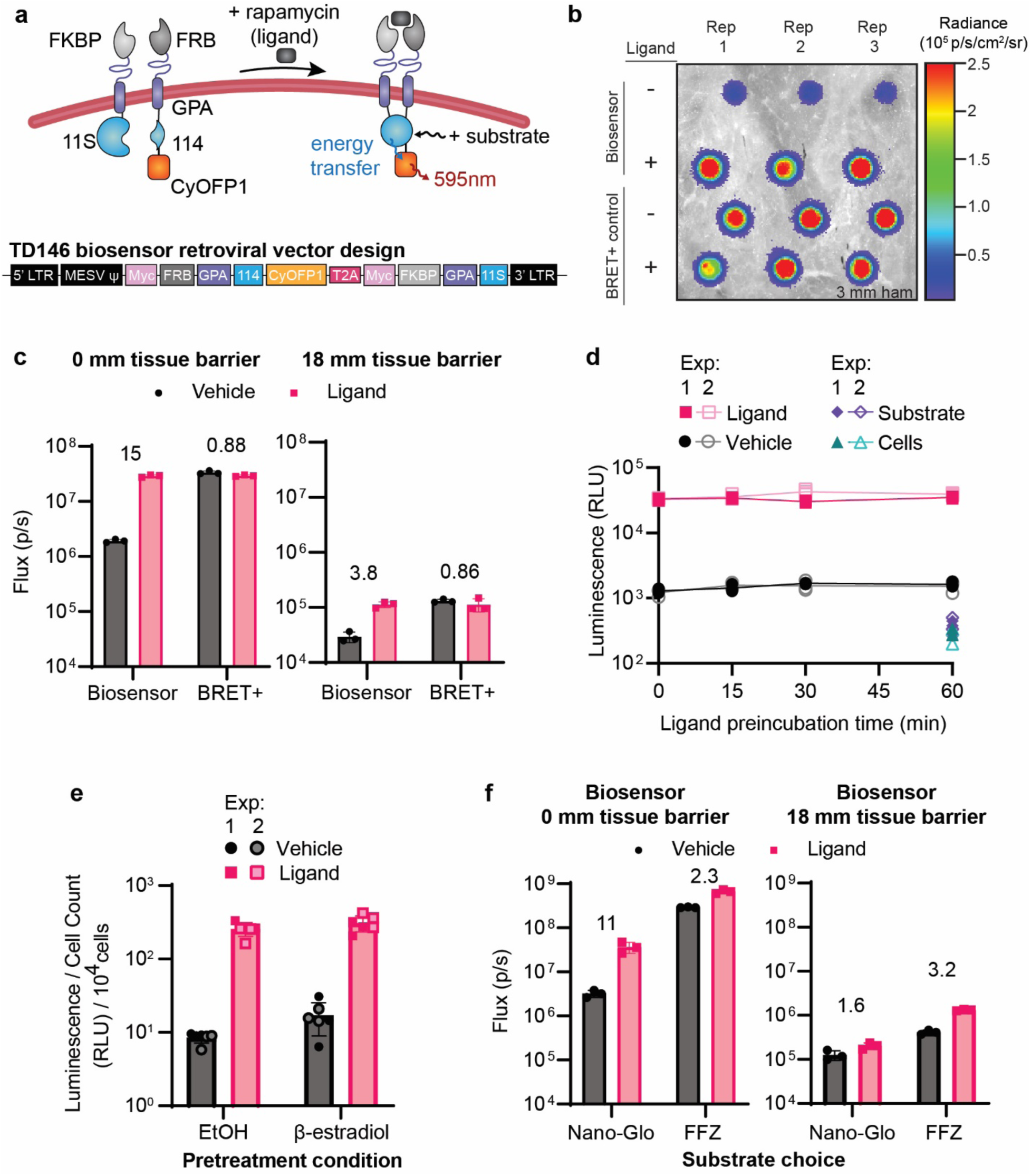
GPA-based BRET biosensors function in G1ER cells. **(a)** Schematic illustrating the structure and proposed mechanism of a GPA-based, BRET output biosensor implemented in G1ER cells. **(b)** Representative IVIS data showing the luminescent signal from BRET biosensors in G1ER cells penetrating 3 mm of tissue mimetic. 1 x 10^6^ BRET biosensor cells were plated and 7.5 x 10^2^ BRET positive control cells were plated to facilitate comparison on the same plate. Three replicates are shown in the columns. The biosensor condition is TD146 from panel (**a**), and the BRET+ control is TD142 from **Figure S9**. The uncropped image is presented in **Figure S10**. (**c**) Quantified IVIS data in luminescent flux from (**b**) and **Figure S10** through either 0 mm (left) or 18 mm (right) tissue mimetic. Each symbol is a biological replicate, the bars represent the mean, the error bars are standard error of the mean (SEM), and the numbers above the bars are the fold induction (signal with ligand divided by signal with vehicle). Trends are representative of two independent experiments. (**d**) Kinetics of biosensor activation, evaluated by plate reader, for BRET biosensor cells pretreated with rapamycin or vehicle prior to addition of the Nano-Glo substrate. “Substrate” refers to conditions that received substrate but not cells or rapamycin; “Cells” refers to conditions that received cells but not rapamycin or substrate. Each symbol represents a biological replicate, the lines connect the mean at each time point, and the error bars are SEM. Two independent experiments were performed with 3 biological replicates in each. (**e**) BRET biosensor function in differentiated G1ER cells. Cells were treated with 100 nm β-estradiol (to induce differentiation towards RBCs) or ethanol vehicle (EtOH) for 2 d before biosensor evalution by plate reader. Each symbol represents a biological replicate, the bars represent the mean, and the error bars are SEM. Two independent experiments were performed with 3 biological replicates in each. (**f**) Quantified IVIS data from **Figure S12** evaluating how choice of NanoLuc substrate (Nano-Glo or fluorofurimazine [FFZ]) affects BRET biosensor performance through 0 mm (left) or 18 mm (right) of tissue mimetic. 1 x 10^6^ BRET cells were plated per well, each symbol is a biological replicate, the bars represent the mean, the error bars are SEM, and the numbers are the fold induction (signal with ligand divided by signal with vehicle). The trends are representative of two independent experiments.

We then investigated whether the BRET biosensor retained activity after treating the G1ER cells with 100 nM β-estradiol, which drives differentiation towards a mature RBC phenotype.^*72*^ The β-estradiol treated cells exhibited greatly reduced growth and developed a different morphology characterized by puncta-like structures (consistent with a normoblast morphology), which differed from vehicle-treated controls (**Figure S11**).^*72*^ In β-estradiol treated cells, BRET biosensor output was similar to non-treated cells when correcting for cell count differences (**Figure 5e**), suggesting that such GPA-based biosensors retain functionality when pushed towards an erythroid state.

Finally, we explored how choice of NanoLuc substrate affected biosensor performance. The substrate used up until this point was furimazine (sold as Nano-Glo), which is challenging to use *in vivo* due in part to low solubility and bioavailability.^*73*^ A newly developed analogue, fluorofurimazine (FFZ), has higher solubility which enables higher dosing and therefore has shown promise for *in vivo* applications.^*73, 74*^ When treating G1ER cells with either vehicle or rapamycin and either Nano-Glo or FFZ, we found that FFZ generated a higher ON state than Nano-Glo, differing by approximately an order of magnitude, for all conditions evaluated (**Figure 5f, S12**). Nano-Glo conferred a higher fold induction than did FFZ in the absence of tissue mimetic material (11 vs 2.3, respectively), but this trend reversed when evaluated in the presence of 18 mm tissue mimetic (1.6 vs 2.3, respectively) (**Figure 5f**). We speculate that this reversal can be explained by the generally lower output when using Nano-Glo compared to FFZ, a difference which increases with thickness of tissue mimetic (**Figure S12**); low output can yield a higher fold induction but generally yields a low ON state as well. We anticipated low signal being a challenge for *in vivo* applications and therefore chose to use FFZ as our *in vivo* substrate for subsequent studies. Taken together, these results suggest that BRET biosensors can be implemented in RBC precursor cell lines and motivate *in vivo* validation of this candidate eRBC.

### In vivo performance of GPA-based BRET biosensors in G1ER cells

As a final validation of the GPA-based BRET biosensors developed in this study, we evaluated whether these biosensors could report on environmental stimuli detected *in vivo*. We injected 10 x 10^6^ engineered G1ER cells via tail vein into BALB/c mice, followed by rapamycin or vehicle administration via IP, and finally FFZ injection via IP (**Figure 6a**). Excitingly, we found that luminescence from mice treated with the BRET biosensors and rapamycin was significantly greater than luminescence from mice treated with biosensors and vehicle at several time points (**Figure 6b,c**). The fold induction observed *in vivo* (between 2.7-4) was similar to the fold induction observed *in vitro* through tissue mimetic (**Figure 6c**, **Figure 5f**), validating our overall biosensor design pipeline, including the use of the tissue mimetic as a useful way to prototype designs *in vitro*. We note that a large portion (> 50%) of the luminescent signal from the vehicle condition could be attributed to background luminescence from the substrate alone. This suggests that substrate choice is also an important design consideration, and metrics such as fold induction or magnitude of the “ON” state can be tuned by choosing alternate substrates and conditions. Ultimately, our results demonstrate that GPA-based BRET biosensors implemented in red blood cell precursors can detect and report on stimuli in their environmental milieu in essentially real time, and these data suggest an exciting path forward in the development of RBC-based products for diagnostic applications.

**Figure 6.**
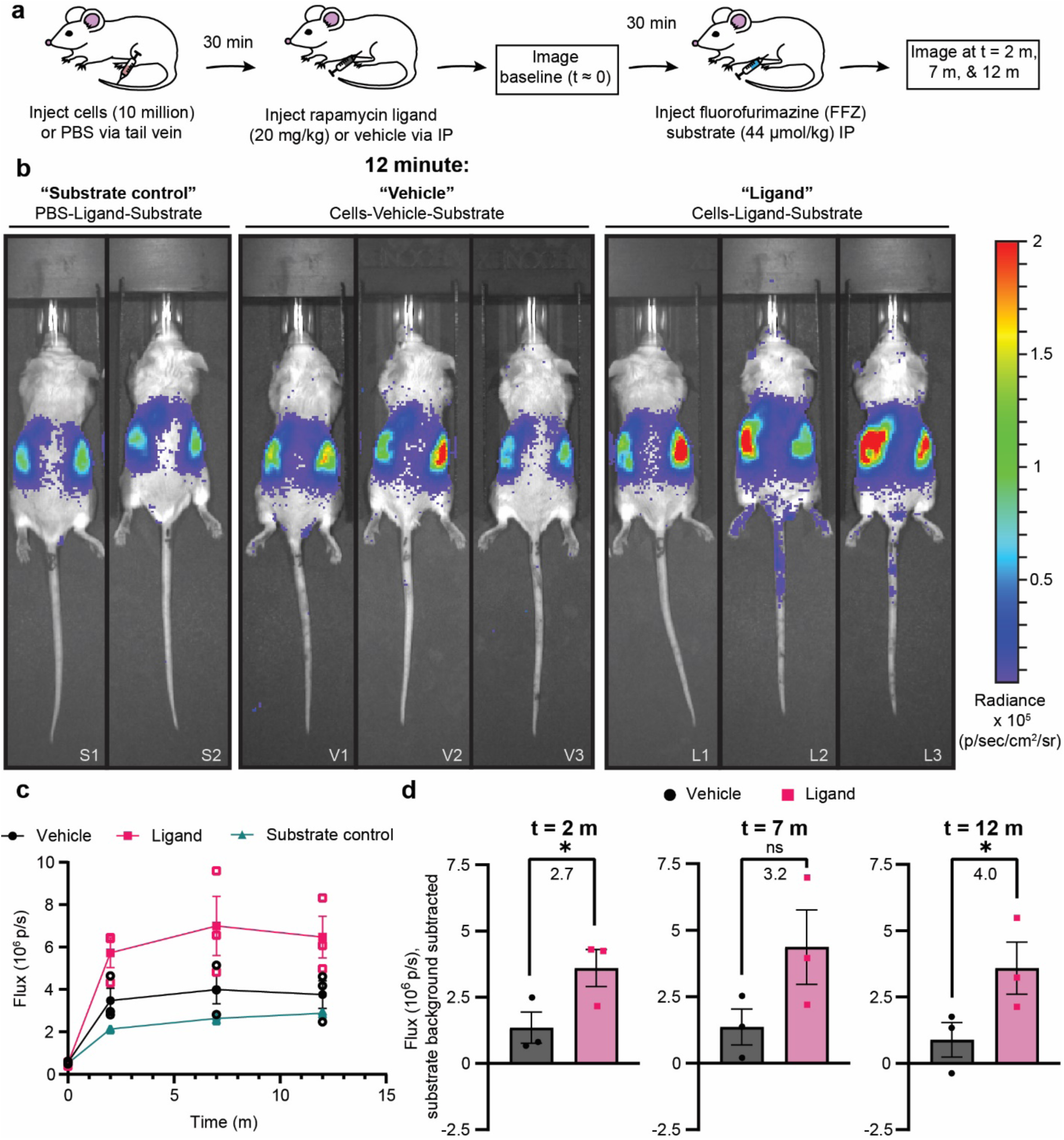
G1ER cells engineered to express GPA-based biosensors achieve detectable ligand sensing in vivo. **(a)** Schematic summarizing the design of the *in vivo* biosensor evaluation study. The ligand condition received cells, rapamycin, and the fluorofurimazine (FFZ) substrate. The vehicle condition received cells, the rapamycin vehicle, and the FFZ substrate. The substrate only condition received PBS instead of cells, rapamycin, and the FFZ substrate. **(b)** Biosensor performance evaluated by in vivo imaging system (IVIS). This figure shows all mice in this study at 12 minutes after substrate injection. Luminescence data are represented as a colored heatmap and are scaled equally for all images. Uncropped images for all time points are available in **Figure S13**. Unique mouse identifiers are present in the lower right hand corner of each image and correspond to the identifiers in **Figure S13**: Substrate “S”, Vehicle “V”, and Ligand “L”. **(c)** Quantified luminescence data from IVIS images presented in (b) and **Figure S13**. A region of interest was drawn around each mouse as presented in **Figure S13** and the flux in photons per second are reported. Open symbols represent individual mice, closed symbols represent the mean of all mice in a given condition at a particular time point. Error bars represent SEM, and the connecting lines are drawn as a visual aid. **(d)** For each time point after substrate injection in (c), the signal attributable to the substrate was subtracted from the vehicle and ligand conditions and plotted. The bars represent the mean, error bars represent the SEM, and each symbol represents an individual mouse. Fold inductions, or the signal with ligand divided by the signal with vehicle, are reported for each plot. An unpaired, one-tailed t-test was performed to determine significance (p values from left to right: 0.0350, 0.0625, 0.0421).

## Discussion

In this study, we developed a strategy for converting RBC membrane proteins into surface-displayed, self-contained biosensors capable of sensing and reporting on the presence of a ligand. Both Kell and GPA are suitable scaffolds for our two chain biosensing architecture, and receptor performance characteristics (e.g., background and inducible signal) could be tuned by exploring alternate chain architectures, inner linker compositions and flexibilities, and scaffold types and mutants. This biosensor output can comprise multiple split protein reconstitution modes including GFP, NanoLuc, and NanoLuc-based BRET for shifting the output light to better enable *in vivo* deployment. Many aspects of receptor performance mapped fairly well between deployment in workhorse HEK293FT cells and erythroid precursor G1ER cells, with some important exceptions (discussed below), which together provide useful insights to guide future receptor development. The final BRET biosensor developed here generated up to 15 fold induction with ligand, was detectable through 18 mm of tissue mimetic, reached maximum signal intensity within 5 min of ligand addition, and retained signaling capability after treatment with an erythroid differentiation signal. Finally, we demonstrated that our RBC-protein based biosensor, implemented in G1ER cells, successfully reported on the presence of our model ligand *in vivo.* Altogether, this work demonstrates the feasibility of building biosensors using RBC membrane proteins.

The biosensors explored here used model inputs (ligand) and outputs (i.e., rapamycin and fluorescence/luminescence, respectively), and these studies helped to identify principles that should be useful for adapting this technology to engineer eRBCs for new applications of interest. First, optimal linker design and receptor geometry are difficult to predict a priori, yet these are very important choices when building synthetic receptors and biosensors.^*27, 37, 43, 75, 76*^ We found that linker tuning and rearranging receptor/scaffold geometry could change both background and inducible signaling in our split GFP implementation (**Figure 2, S4, S6)**. Second, implementing the biosensors in an RBC-like membrane environment required addressing two key considerations: (i) chain expression levels and (ii) chain association propensity. These two factors greatly affected biosensor inducibility and differed between the characterization cell line (HEK293FTs) and the murine erythroid cell line (G1ERs). For example, the mutant V84R GPA chains capable of inducible signaling in HEK293FTs (**Figure 3b**) were not able to signal in G1ERs (**Figure S7c**) and this effect can be partially explained by poor surface expression in the G1ER cell line (**Figure S7b**). Chains that do not express well on the cell surface in the desired cellular context are unlikely to produce useful biosensors. Third, detecting the biosensor *in vivo* was enabled by using the red-shifted BRET output rather than the more blue-shifted NanoLuc luminescence (**Figure 6)**. We anticipate that these lessons will benefit future bioengineering efforts.

Building on the fundamental proof of feasibility established in this study, several technical hurdles remain for evaluating and developing eRBCs for diagnostic use. Although rapamycin is a useful model ligand, building RBC-protein biosensors that respond to health and disease-associated cues (e.g., growth factors, cytokines) is an important next step. One validated strategy for such extensions is fusing affinity reagents (e.g., antibody fragments, nanobodies, or natural receptor domains) onto the GPA and Kell scaffolds, as we and others have done for other receptor systems.^*27, 37, 75, 76*^ Another technical challenge comprises choosing and developing a split protein output compatible with human monitoring. Interesting avenues include reconstitution of a protease to modulate the ultrasound profile of encapsulated gas vesicles,^*77*^ or to release a surface-bound synthetic biomarker that is excreted in a patient’s urine.^*78, 79*^ There exist many approaches for choosing protein split sites, and we recently developed a computation-guided method for tuning reconstitution propensity to achieve high performance when split proteins are used in applications such as the engineered biosensors described here.^*45*^ Finally, we validated our BRET biosensor in G1ER cells over a short period of time (< 90 min). It will be important to subsequently evaluate whether translationally viable differentiated erythrocytes (e.g., generated from primary cells or suitable immortalized sources^*80*^) function *in vivo* over a clinically relevant timescale (ideally, the native RBC lifespan of 90 – 120 days, in humans). These future research avenues are exciting potential advances in eRBC technology that are enabled by the foundations established in this study.

We envision myriad opportunities for using eRBC biosensors in clinical applications. We expect that eRBC biosensors would benefit patients most when non-invasive, frequent monitoring is needed. As an illustrative example, a potentially fatal side effect after chimeric antigen receptor (CAR) T cell cancer therapy is a rapid hyper-inflammatory state termed cytokine release syndrome (CRS).^*81*^ A main driver of CRS’s toxicity is high interleukin-6 (IL-6) in circulation; interventions to block IL-6 or suppress the immune system can alleviate the severe side effects but require timely intervention and in some cases impact the efficacy of the cancer therapy.*^1,^* ^*81*^ In this example, early *in situ* detection of the physiological cue of disease, IL-6, via eRBCs could provide clinicians with actionable information to drive changes in treatment regimens. One could envision similar use cases for eRBC biosensors in detecting early indications of graft vs. host disease following allogeneic stem cell transplant;^*82*^ monitoring patients on blood thinners for d-dimer levels to guide dosing and duration of anticoagulant drugs;^*83*^ and monitoring natriuretic peptide levels in patients at risk of congestive heart failure.^*84*^ Broadly, we envision eRBC diagnostics as tools that could provide previously inaccessible information to physicians and patients, helping to reduce morbidity and mortality for a wide range of conditions. The RBC-protein based biosensors developed in this study comprise a key first step towards realizing this potential.

## Data Availability

The data generated in this study are available in the published article and its Supporting Information (**Supplementary Data 2**). Any additional information requested will be made available upon reasonable request.

## Supporting Information

Supplementary Information: flow cytometry gating strategy (Figure S1); surface staining for initial Kell and GPA-based biosensors (Figure S2); GFP-based, scaffold and linker evaluation for WT scaffold proteins (Figure S3); evaluation of cooperative domains on biosensors (Figure S4); expression of mutant GPA biosensors in HEK293FT cells (Figure S5); functional evaluation of mutant GPA biosensors with varying linker lengths (Figure S6); evaluation of split NanoLuc in G1ER cells (Figure S7); BRET output evaluation in HEK293FTs (Figure S8); characterization of BRET-based biosensors in G1ER cells (Figure S9); TD146 evaluation through tissue mimetic (Figure S10); morphology of TD146 cells after exposure to a differentiation signal (Figure S11); functional evaluation of TD146 through tissue mimetic for Nano-Glo and FFZ substrates (Figure S12); uncropped IVIS images for the *in vivo* studies presented in Figure 6 (Figure S13); example region of interest used when quantifying biosensor performance *in vivo* (Figure S14) (PDF)
Supplementary Data 1: Plasmid maps for all plasmids generated in this study; table describing which plasmids were used in which figures (.Zip)
Supplementary Data 2: Raw and analyzed source data for Figures 2b, 2d, S3a-c, S4a-b, S6, 3b, 3d, S7c, s9e, 5c-f, S10-S12, 6c-d; source images for westerns blots (Figures s5b, s9b), and IVIS data (Figures 4b, 4d, 6, S8, S9e, S10, S12 (.Zip)

## Acknowledgements

This work was supported in part by the National Institute of Biomedical Imaging and Bioengineering of the NIH under award number R01EB026510 (J.N.L.), by the Northwestern University Clinical and Translational Sciences Institute (NUCATS) and by a gift from Kairos Ventures. T.B.D. was supported by the Department of Defense (DoD) through the National Defense Science & Engineering Graduate Fellowship (NDSEG) Program. T.F.G. was supported by an NSF Graduate Research Fellowship (DGE-1842165). Any opinion, findings, and conclusions or recommendations expressed in this material are those of the authors(s) and do not necessarily reflect the views of the National Science Foundation. This work was also supported in part by the Northwestern University Flow Cytometry Core Facility supported by Cancer Center Support Grant (NCI 5P30CA060553), and the NUSeq Core of the Northwestern Center for Genetic Medicine. IVIS imaging work was performed at the Northwestern University Center for Advanced Molecular Imaging (Evanston) and Northwestern University Center for Advanced Microscopy (Chicago), both supported by NCI CCSG P30 CA060553 awarded to the Robert H Lurie Comprehensive Cancer Center. Animal studies were performed by the Northwestern University Developmental Therapeutics Core supported by NCI CCSG P30 CA060553 awarded to the Robert H Lurie Comprehensive Cancer Center. We would like to thank Nayereh Ghoreishi-Haack for her assistance planning the animal studies, and Elizabeth Dempsey for her expertise in both planning and executing the animal studies. We would also like to thank Malini Rammohan provided guidance on *in vivo* studies, Dr. Alexandra De Lille who offered invaluable input informing the method of IVIS detection of BRET, and Dr. Chad Haney who assisted with IVIS imaging.

