## Supplementary Information for "Building synthetic biosensors using red blood cell proteins"

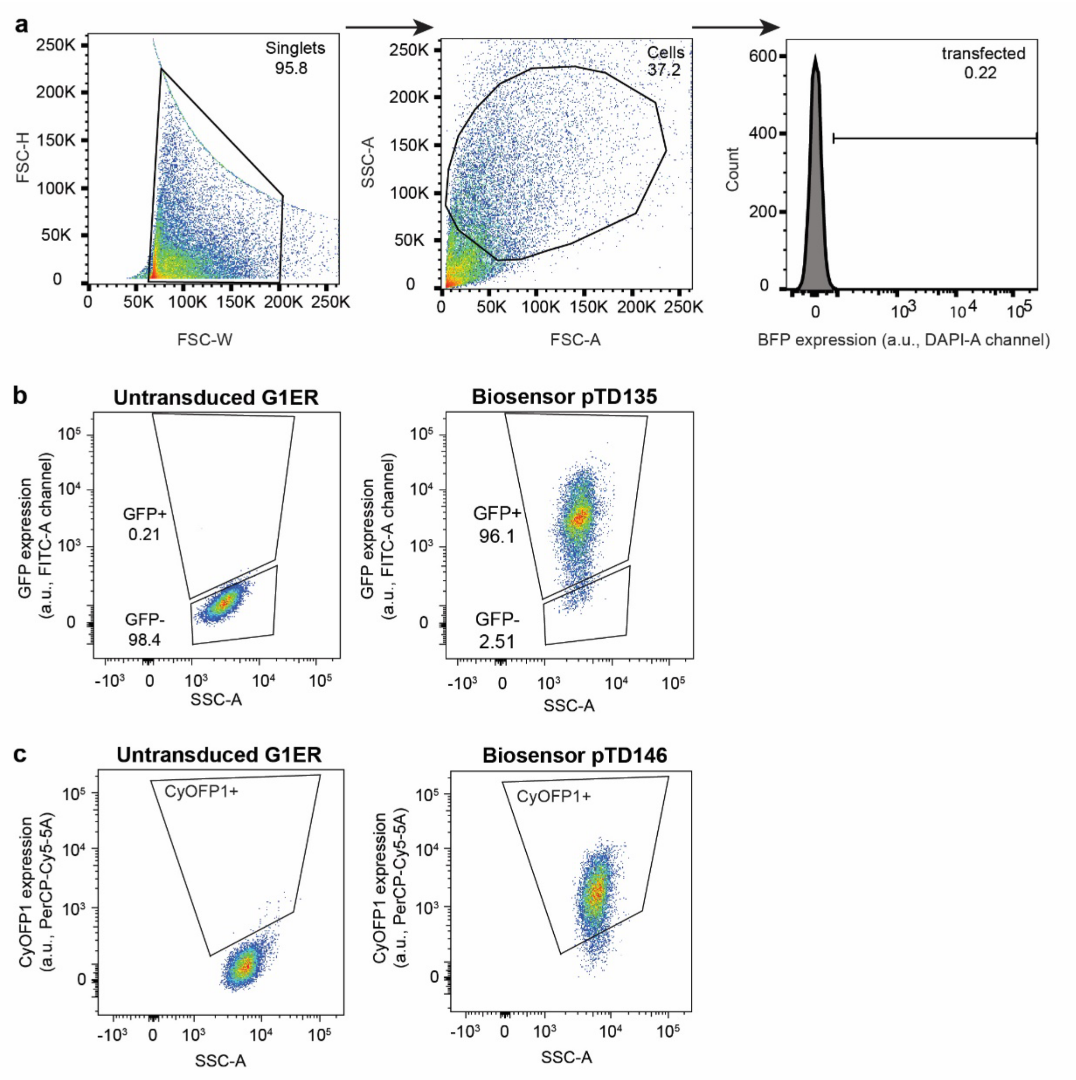


**Figure S1. Flow cytometry gating strategy for experiments in this study. (a)** Representative flow cytometry gating for transfected HEK293FT studies involving GFP reconstitution or surface staining. Events were first gating for singlets using forward scatter height (FSC-H) against FSC width (FSC-W). Singlets were then gated for cells using FSC-area (FSC-A) against side scatter area (SSC-A). Single cells were then gated for expression of a co-transfected fluorescent protein (here, BFP). DsRed was also used in certain cases. Single, transfected cells were then analyzed further depending on the experiment. **(b)** Representative gating strategy for sorting of G1ER cells expressing NanoLuc-based GPA constructs, which co-express GFP from a co-integrated promoter. **(c)** Representative gating strategy for sorting of G1ER cells expressing BRET-based GPA constructs, which contain a CyOFP1 fluorescent protein.

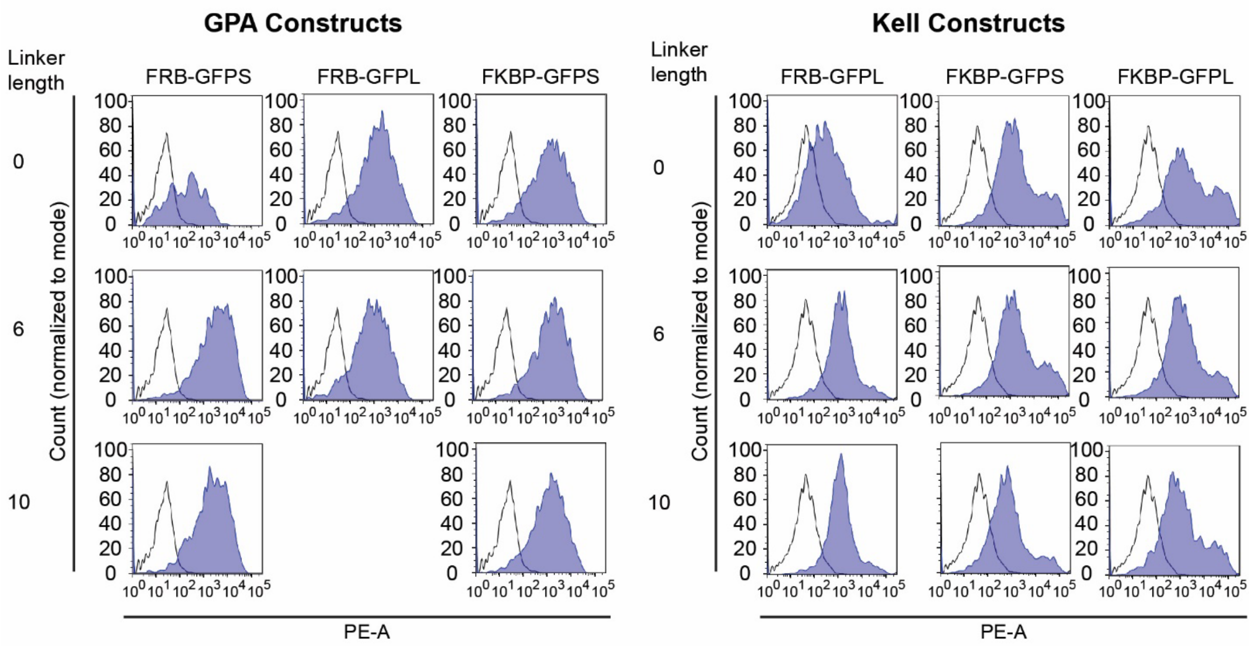


**Figure S2. Surface staining of initial Kell and GPA-based biosensors with various inner linkers demonstrate similar surface expression for most constructs.** Surface staining profiles of HEK293FT cells expressing myc-tagged GPA and Kell-based biosensor chains, shown in each plot as the purple histograms. A stained vector-only control sample is shown in each plot as the black-lined histogram. All constructs investigated with various inner linkers (0, 6, 10 glycine-serine linkers), either rapamycin binding domain (FRB or FKBP), and either GFP half (GFPL or GFPS) showed surface expression. This experiment was performed once.

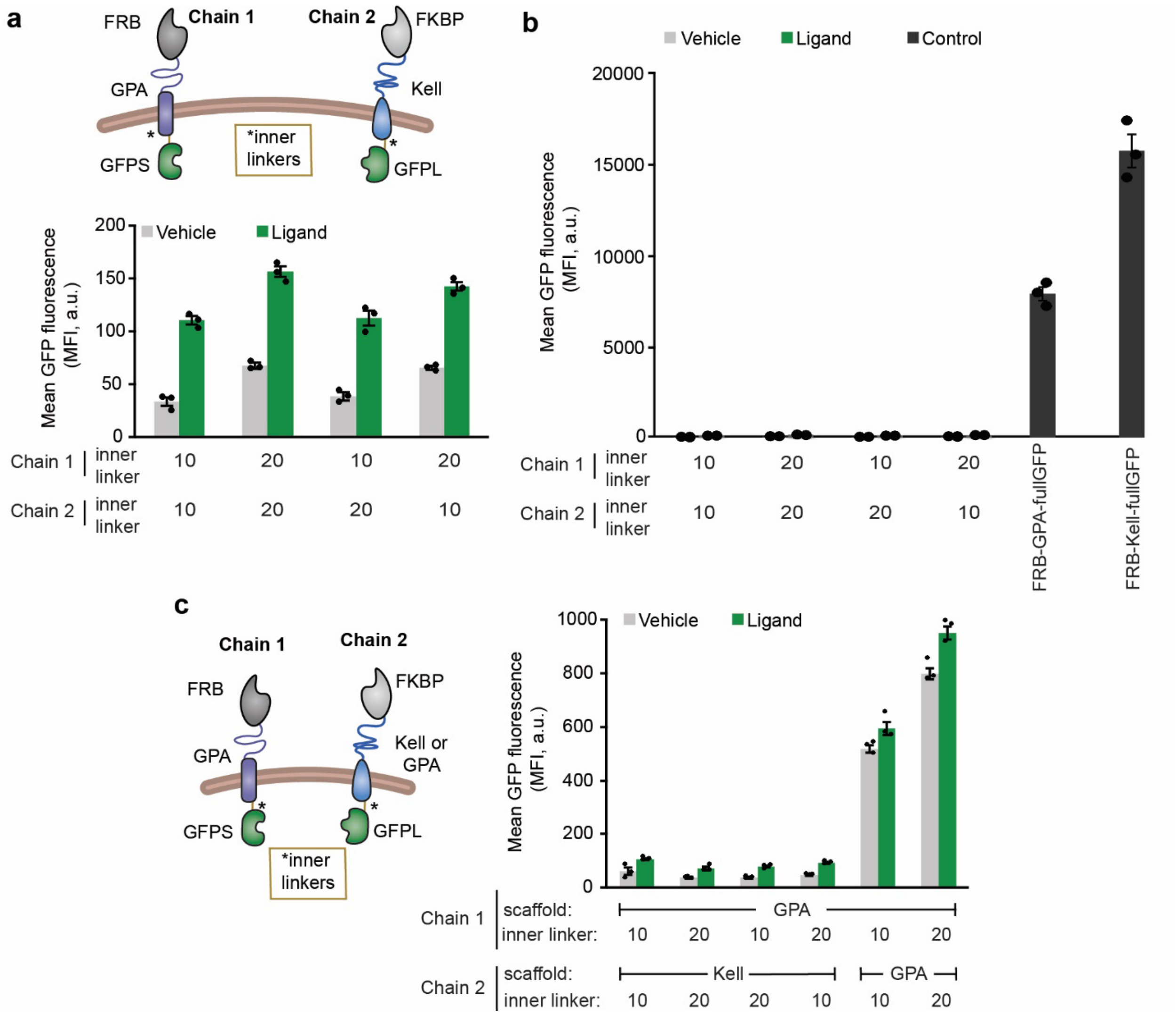


**Figure S3. An eRBC-based biosensor reconstitutes split GFP in the presence of ligand, although the magnitude of fluorescence is substantially reduced compared to a full GFP control. (a)** From **Figure 2b**, the architecture with the highest GFP fluorescence upon ligand treatment (FRB and GFPS on the GPA chain and FKBP and GFPL on the Kell chain) was chosen to investigate extended and mixed inner linkers. Biosensor function was evaluated by flow cytometry. Extending the inner linkers from 10 to 20 residue glycine-serine linkers increased the GFP reconstitution with ligand treatment. **(b)** Data reproduced from panel (a) alongside signals from two controls shown in black bars on the right: 1) a GPA chain with full GFP and 2) a Kell chain with full GFP. The magnitude of the reconstituted GFP in comparison to the full GFP suggested that split GFP reconstitution was limited or impaired. **(c, left)** Schematic depicting an alternate biosensor strategy wherein the scaffold for "Chain 2" may be Kell or GPA. **(c, right)** Functional analysis of first generation Kell-GPA biosensor with 10 and 20 inner linkers compared to a GPA only biosensor with 10 and 20 inner linkers. The GFP fluorescence for the GPA biosensors was substantially higher than the GFP fluorescence for GPA-Kell biosensors. The symbols represent biological replicates, the bars represent the mean, and the error bars represent SEM. Trends throughout are representative of two similar experiments. MFI, mean fluorescence intensity.

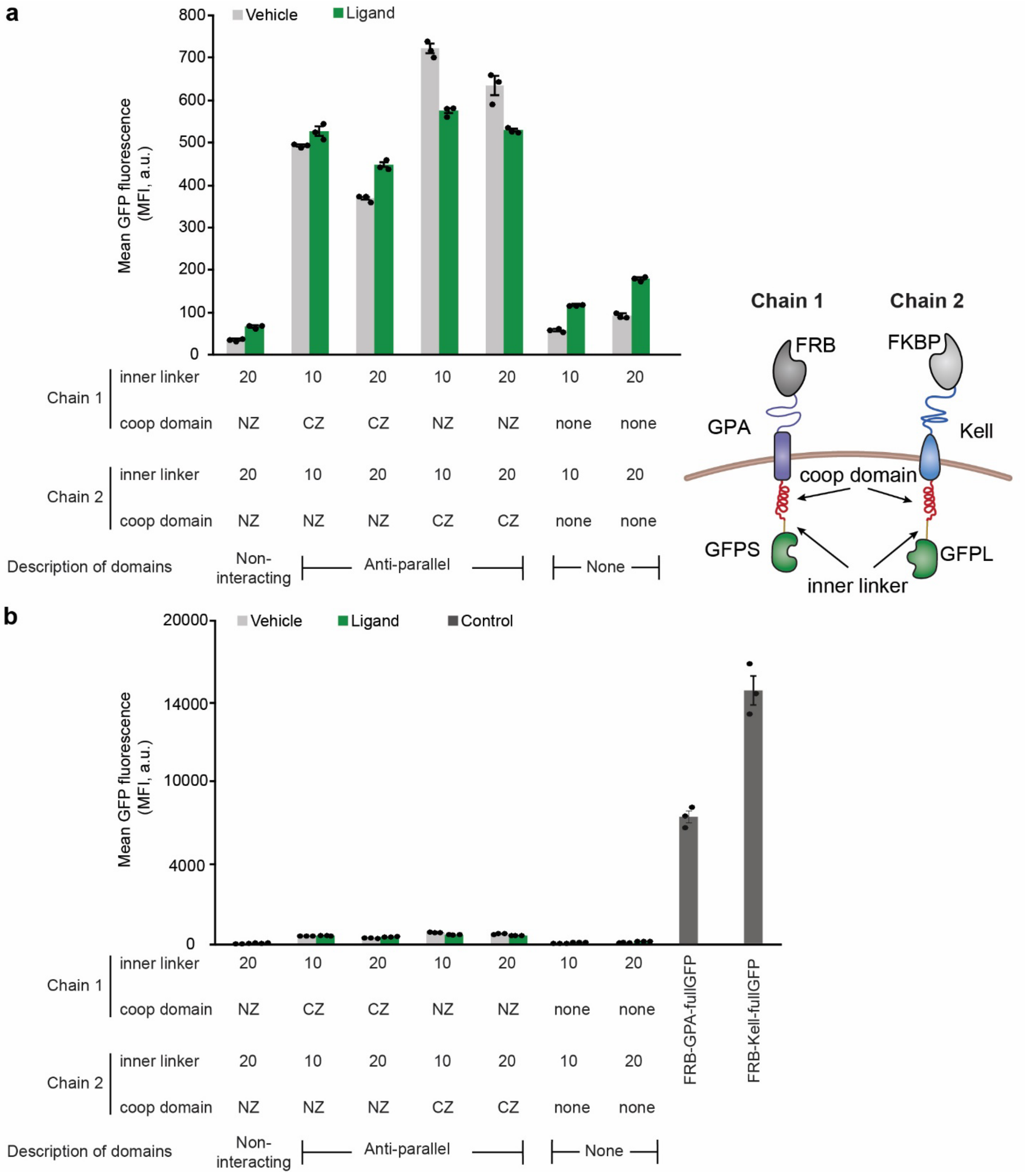


**Figure S4. Introducing cooperative binding domains into Kell and GPA-based biosensors generated substantial ligand-independent signaling and did not improve biosensor performance. (a)** Cooperative binding domains (shown in red) genetically fused to the interior of GPA chain 1 (containing FRB and GFPS domains) and Kell chain 2 (containing FKBP and GFPL domains) biosensor chains to facilitate GFP reconstitution upon ligand addition. Biosensor performance was evaluated by flow cytometry. Anti-parallel leucine zippers (CZ/NZ) were tested, where CZ and NZ are known dimerizing pairs. The NZ paired with NZ condition was used as a non-interacting control. The addition of cooperative binding domains increased magnitude of both background and ligand-mediated GFP reconstitution, however most combinations of anti-parallel cooperative binding domains with 10 and 20 glycine-serine linkers produced little or no ligand-induced signaling. **(b)** Same data as shown in panel (a), with controls of GPA chain with full GFP and Kell chain with full GFP shown in the black bars on the right. The magnitude of the reconstituted GFP in comparison to the full GFP showed the GFP reconstitution was severely limited. This experiment was run on the same day as the study in Figure S3b, and therefore the full GFP bars represent the same samples between the two figures. The symbols represent biological replicates, the bars represent the mean, and the error bars represent SEM. This experiment was performed once. MFI, mean fluorescence intensity.

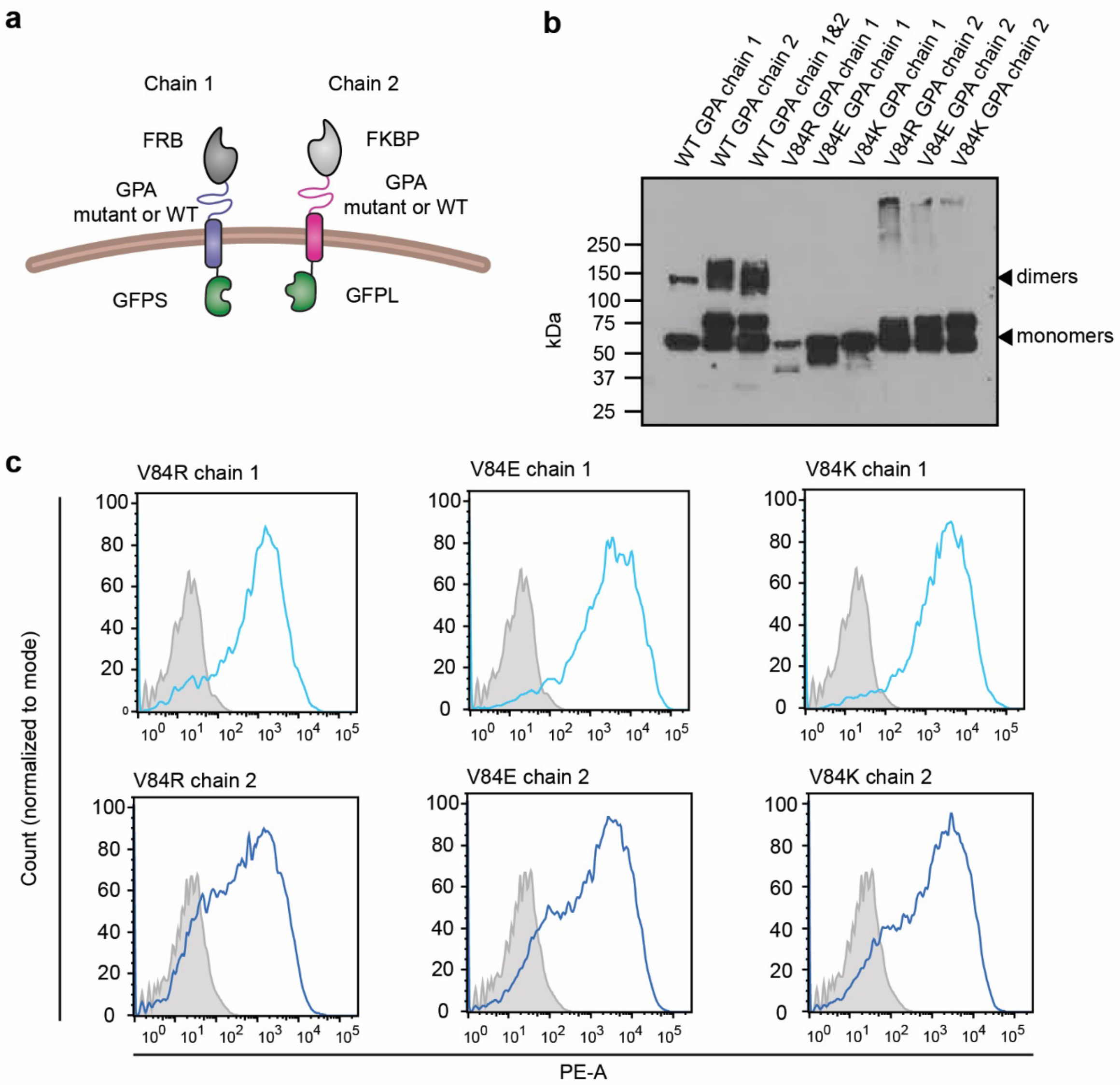


**Figure S5. WT and mutant GPA-based biosensors are expressed in HEK293FTs and traffic to the cell surface. (a)** The schematic illustrates the GPA-based biosensor chains examined here. Chain 1 contains FRB rapamycin binding domain and GFPS with WT or mutant GPA, and chain 2 contains FKBP rapamycin binding domain and GFPL with WT or mutant GPA. **(b)** Western blot analysis of WT and mutant GPA biosensor chains. Cells were transfected with a single myc-tagged chain 1 or myc-tagged chain 2 containing WT or a mutant GPA (V84R, V84E, V84K), or paired chains (3rd column). Equal masses of protein (3 ug/lane) were loaded into each lane to investigate chain expression level and presence of dimers. The WT GPA chains show larger size bands, corresponding to chain dimers. The mutant GPA chains do not show the existence of dimer bands, supporting the hypothesis that the V84 mutations diminish GPA dimerization. **(c)** Surface staining profiles of three different mutant GPA biosensor chains. HEK293FT cells expressing myc-tagged GPA chains are shown in each plot as the blue outlined histograms. A stained vector-only control sample is shown in each plot as the gray histogram. All GPA mutant chains investigated showed surface expression. Each experiment was performed once.

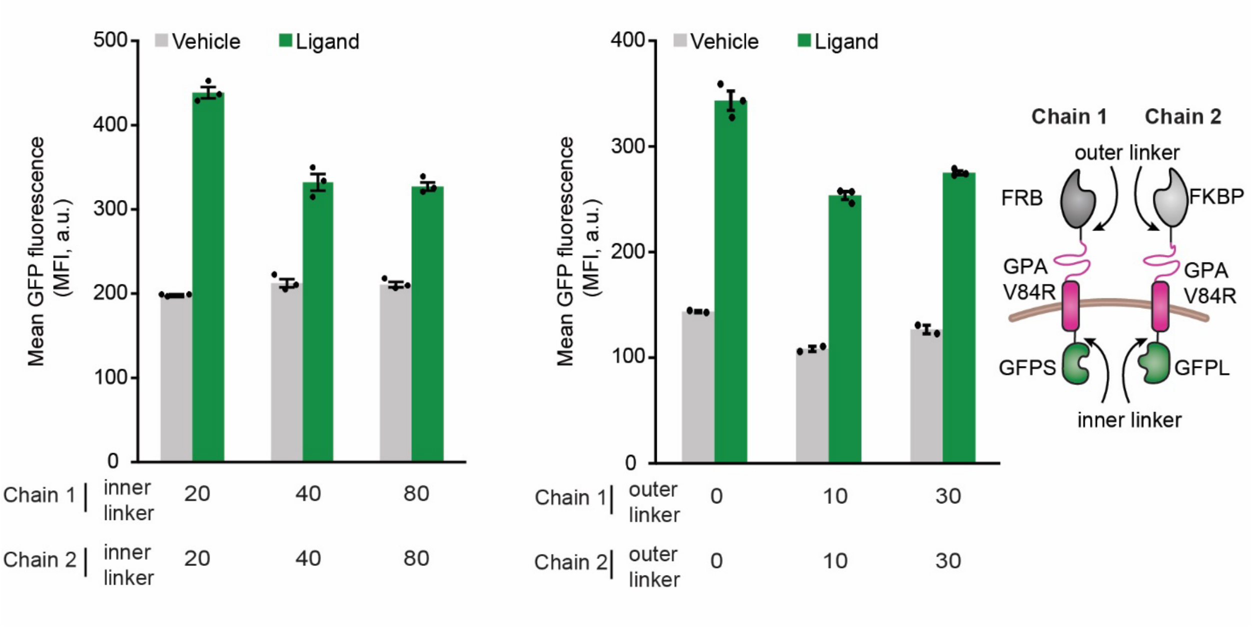


**Figure S6. Changing the length of the inner and outer linkers of GPA-based biosensors did not substantially improve receptor performance for the lengths evaluated.** Experimental analysis of increasing inner and outer flexibility on V84R GPA-based biosensors with addition of glycine-serine linkers, depicted in the cartoon on the right. Biosensor function was evaluated by flow cytometry. Increasing inner linkers beyond 20 amino acids did not increase the signal of rapamycin mediated GFP reconstitution. Adding outer linkers to the GPA chains also did not increase rapamycin mediated GFP reconstitution. An architecture with 20 inner glycine-serine linkers and 0 outer linkers was carried on for further investigation. The symbols represent biological replicates, the bars represent the mean, and the error bars represent SEM. This experiment was performed once. MFI, mean fluorescence intensity.

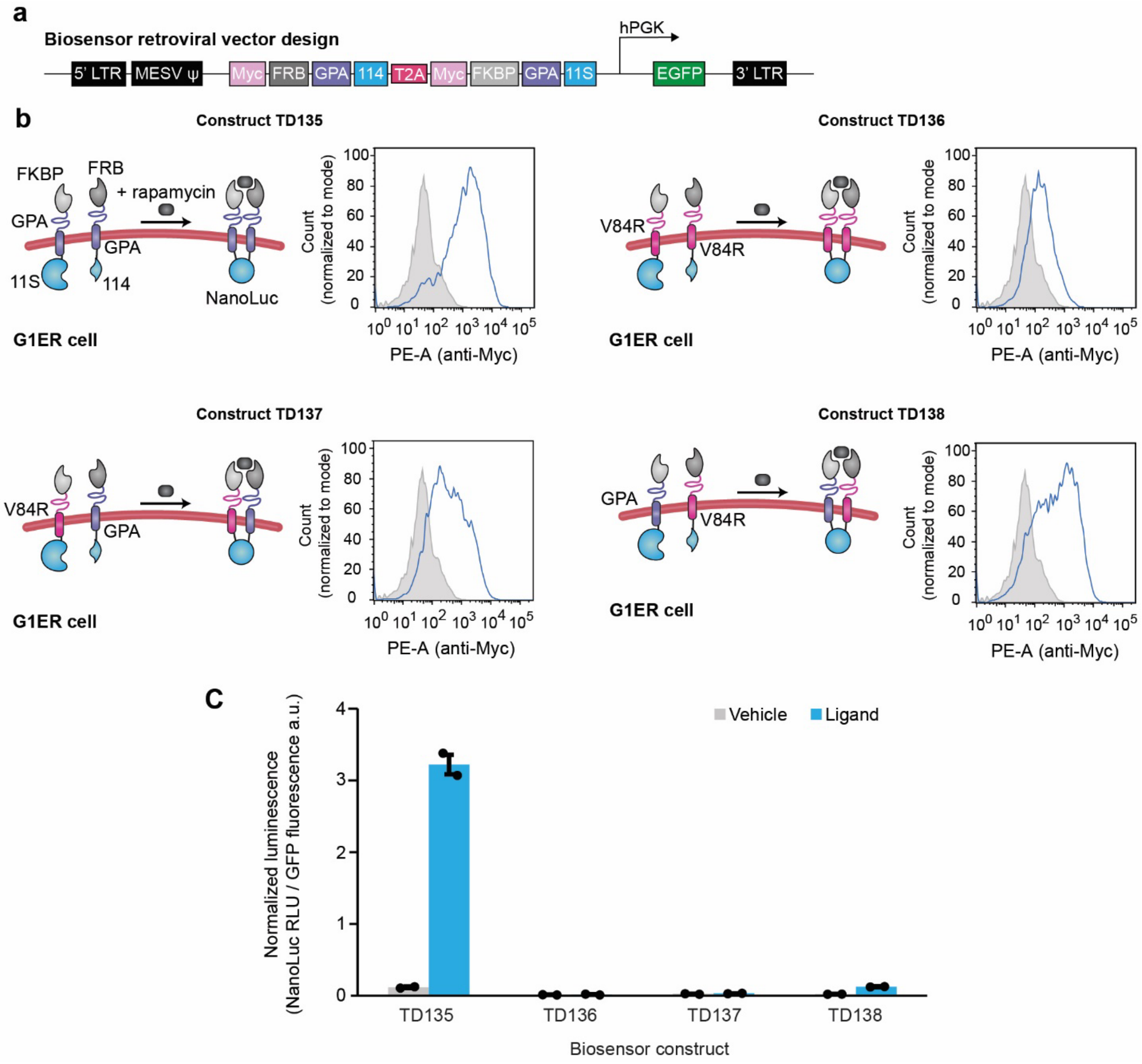


**Figure S7. GPA-based biosensors with a split Nanoluciferase output are surface expressed in G1ER cells. (a)** Retroviral vector design for transducing G1ER cells. Within each vector, there are two biosensor chains separated by a T2A peptide and GFP under an PGK promoter for selection and sorting. **(b)** Four retroviral constructs (TD135, TD136, TD137, TD138) were evaluated for surface expression in G1ER cells. The four constructs differ in their choice of WT or Mut GPA on each biosensor chain; the specific choices are presented in the corresponding cartoon. Surface staining profiles of G1ER cells expressing myc-tagged biosensor chains are shown in each plot as the blue histograms. Non-transduced G1ER cells were used as a stained control sample shown in each plot as the gray histogram. All four constructs showed surface expression of biosensor chains, with the double mutant biosensor showing the lowest surface expression (TD136). **(c)** Functional analysis of GPA-NanoLuc biosensor chains in G1ER cells for the constructs in **b**. Reconstituted NanoLuc luminescence was measured with a plate reader after addition of furimazine substrate and normalized to GFP fluorescence (expressed from retroviral vector). Constructs containing a mutant GPA chain (TD136, TD137, and TD138) exhibited minimal ligand-induced reconstitution of NanoLuc and were not evaluated further. Construct TD135, built on two WT GPA domains, was taken forward (**Figure 3e-f**). The symbols represent technical duplicates, the bars represent the mean, and the error bars represent SEM. Each experiment was performed once.

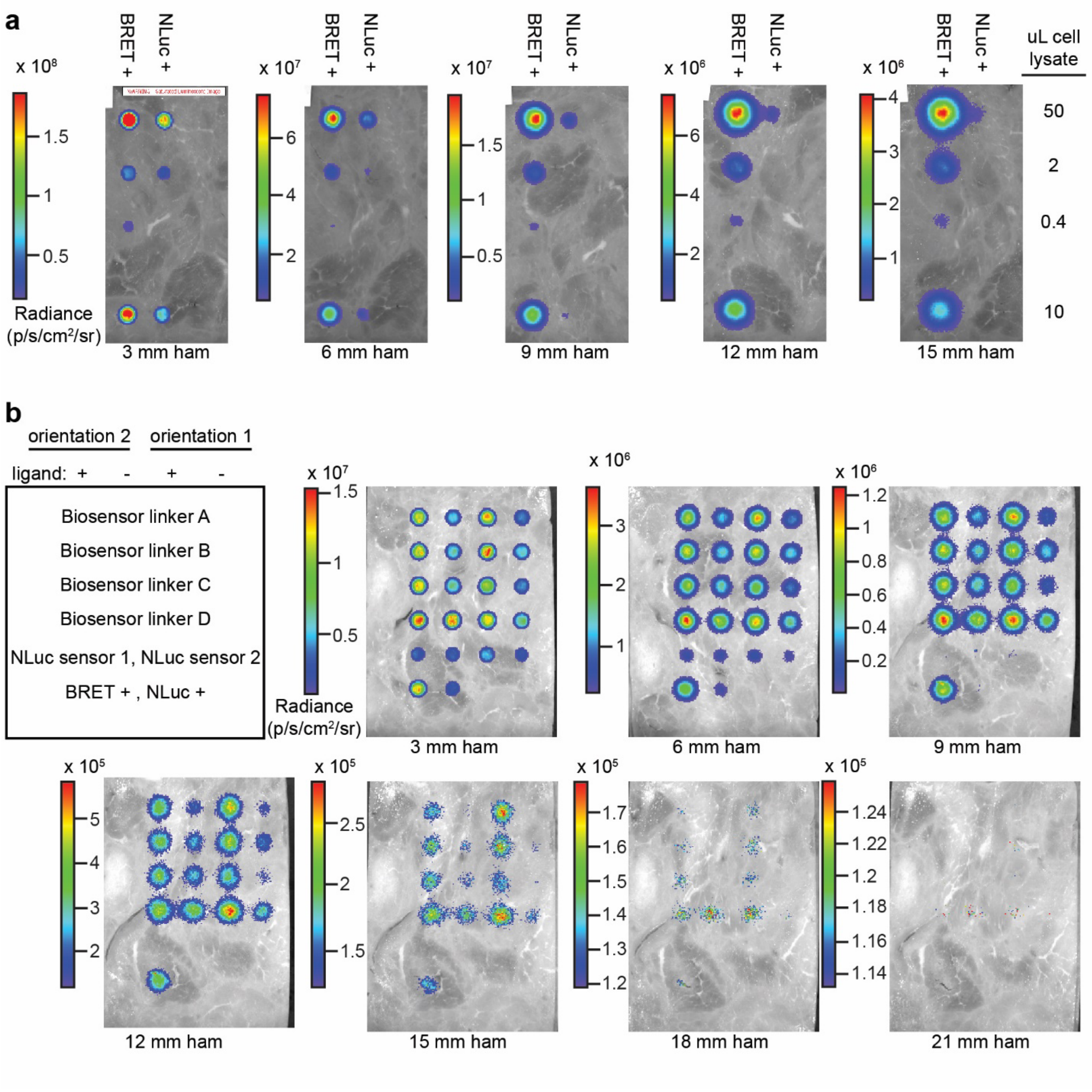


**Figure S8. GPA-based biosensors generate detectable BRET signal through up to 15 mm of tissue mimetic when implemented in HEK293FT cells. (a)** Positive control BRET and NanoLuc signal detection in transiently transfected HEK293FT cells imaged with IVIS and covered with ham from **Figure 4b**. With increasing thickness of ham, the BRET + signal was still detectable and the NanoLuc + signal was undetectable. **(b)** BRET biosensor signal detection in HEK293FT cells imaged with IVIS and covered with ham from **Figure 4d**. Two orientations and four different linkers between 114 NanoLuc domain and CyOFP1 fluorescent protein were examined. GPA biosensors with NanoLuc only (no CyOFP1) were included to compare BRET and NanoLuc signal. NanoLuc sensor 1 has an FRB-GPA-114 and FKBP-GPA-11S architecture, and NanoLuc sensor 2 has an

FKBP-GPA-114 and FRB-GPA-20 architecture. BRET + control and NanoLuc + control lysates (diluted 1:125) were also included. Signal was detected for all samples, with BRET samples being detectable with increasing thickness of ham. Orientation 1, linker D was carried on. Each experiment was performed once.

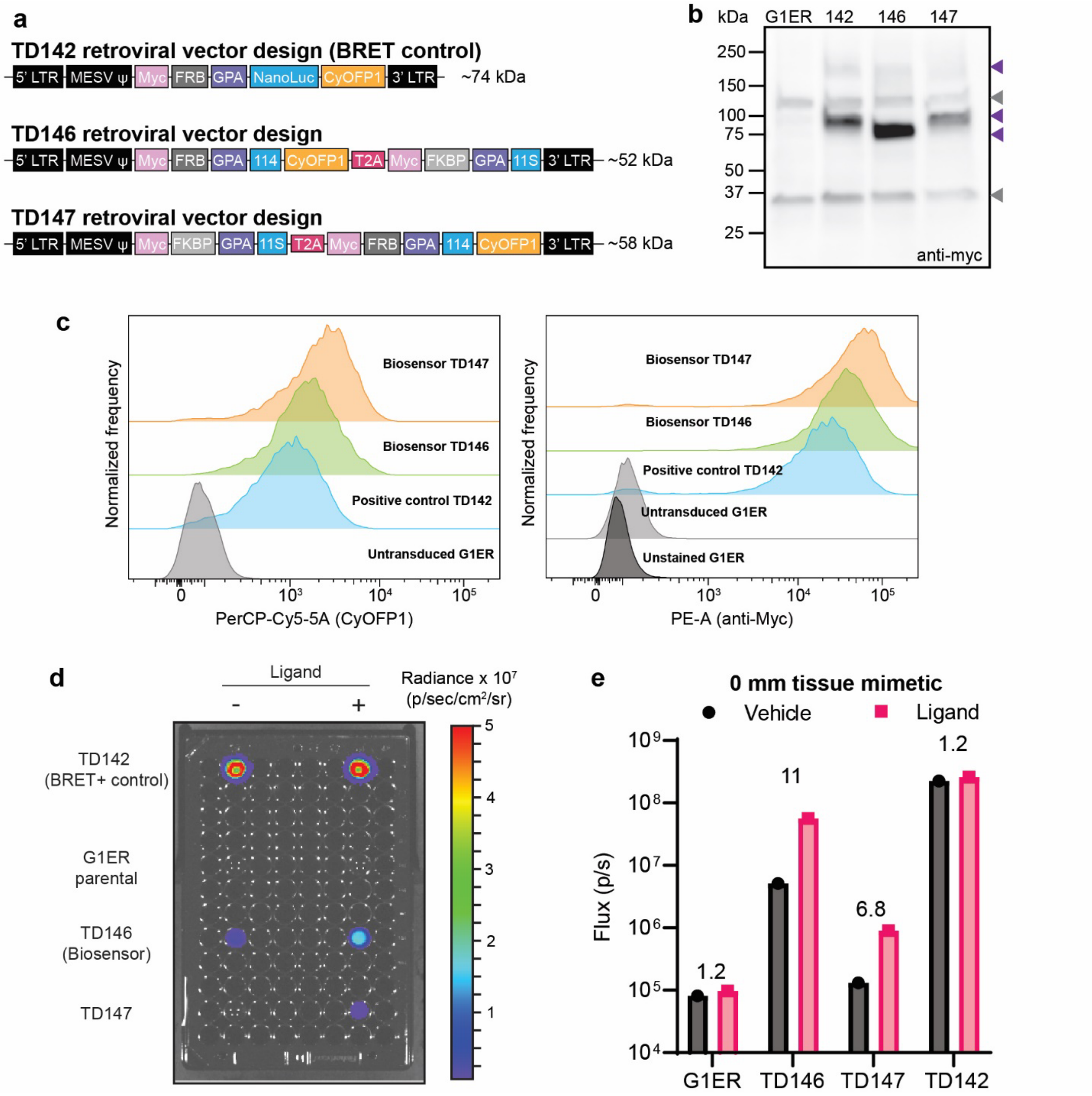


**Figure S9. BRET biosensors TD146 and TD147 are surface expressed in G1ER cell lines after sorting but TD146 produces higher signal in response to rapamycin. (a)** Schematic of positive BRET control TD142 and biosensor constructs TD146 and TD147 built for evaluation in G1ER cells. The positive control consists of FRB rapamycin binding domain, WT GPA, full NanoLuc, and CyOFP1. Biosensor constructs TD146 and TD147 consist of two chains built using WT GPA as a scaffold; chain 1 has FRB and 114 domains, with CyOFP1 tethered to 114 with RH linker, and chain 2 has FKBP and 11S domains. The two chains are separated by a T2A peptide in the retroviral vector. TD146 has chain1-T2A-chain2 orientation and TD147 has chain2-T2A-chain1 orientation. CyOFP1 expression is used for identifying and sorting biosensor expressing cells. **(b)** Anti-myc western blot analysis of G1ER cells transduced with TD142, TD146, and TD147 show detection of biosensor chains in cell lysates, compared to untransduced G1ER cells.

Equal amounts of cell lysate were added to each well. The gray arrows highlight non-specific background bands in G1ERs. The purple arrows highlight expected sizes of the various biosensors as monomers (bottom two) or a dimer (top). **(c)** Flow cytometry profiles of untransduced G1ER cells, positive control vector TD142, biosensor vector TD146, and TD147 transduced G1ER cells. Left, histogram profiles of G1ER cells showing CyOFP1 expression (PerCP-Cy5-5A). Right, G1ER cells stained with anti-Myc-PE antibody evaluated for surface expression of myc tagged biosensor and positive control chains. TD142, TD146, and TD147 showed surface expression in G1ER cells. **(d)** IVIS data showing the luminescent signal from BRET biosensor constructs TD146 and TD147 in G1ER cells. $4 \times 10^6$ BRET biosensor cells were plated and $3.3 \times 10^3$ BRET positive control cells were plated to facilitate comparison on the same plate **(e)** Quantification of the data in **(d)**, showing that TD146 has greater ligand-induced signal than TD147; therefore, TD146 was used going forward. In this experiment, a single biological replicate was performed; therefore, the symbols and bars represent the single value quantified, and there are no error bars. This result is representative of two independent experiments.

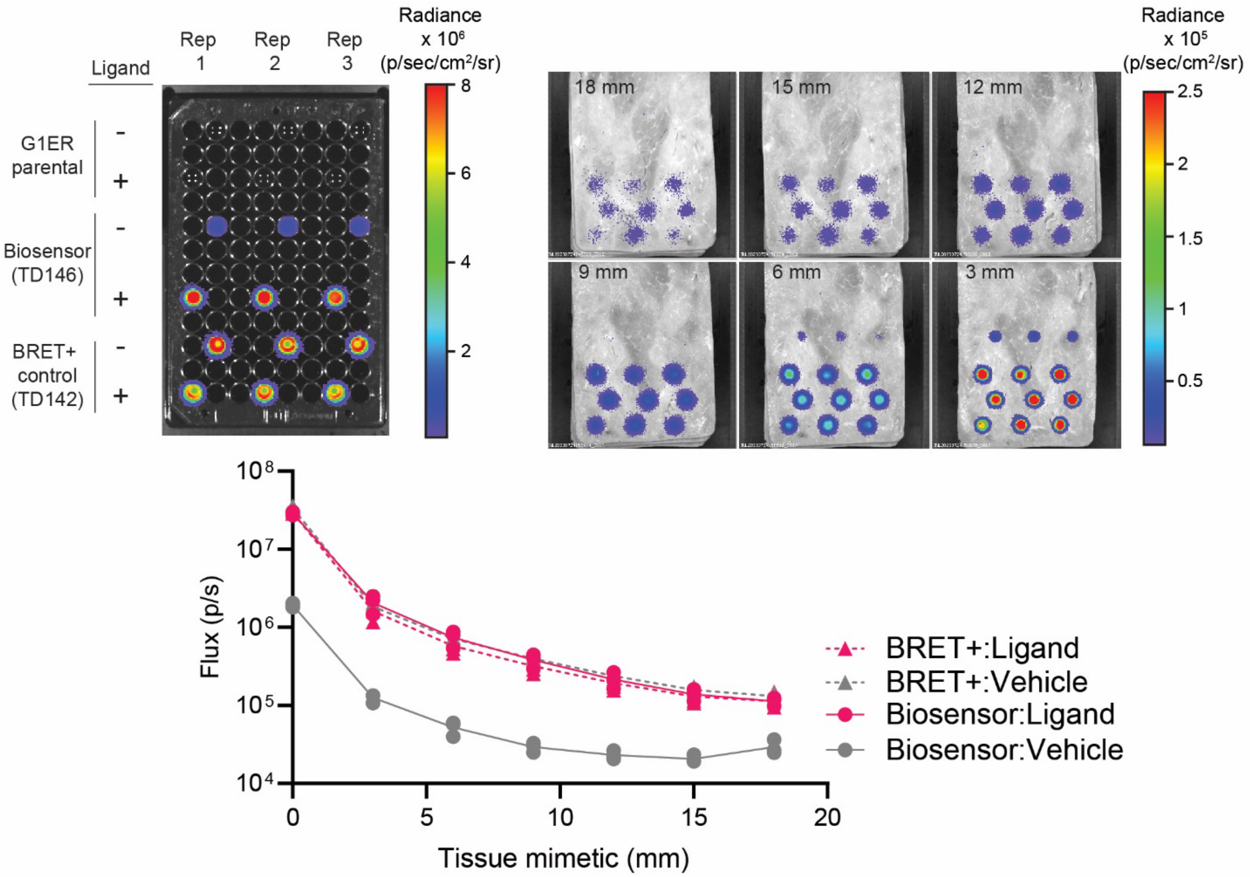


**Figure S10. Uncropped IVIS data from Figure 5b and 5c demonstrate that TD146 BRET biosensor can signal through many layers of tissue mimetic. (a, top)** IVIS data showing the luminescent signal from BRET biosensor construct TD146 in G1ER cells through no ham or up to 18 mm of ham. 1 x $10^6$ TD146 BRET biosensor cells were plated and 7.5 x $10^2$ TD142 BRET positive control cells were plated to facilitate comparison on the same plate. Three replicates are shown in the columns. The 3 mm tissue mimetic condition is also shown in **Figure 5b**. **(a, bottom)** Quantification of the data in the top of the figure, showing that TD146 (Biosensor) retains ligand-induced signal through all thicknesses of tissue mimetic evaluated. The symbols represent the three biological replicates shown, the lines connect the means of the replicates at each tissue thickness, and the error bars represent SEM. The trends are representative of two independent experiments.

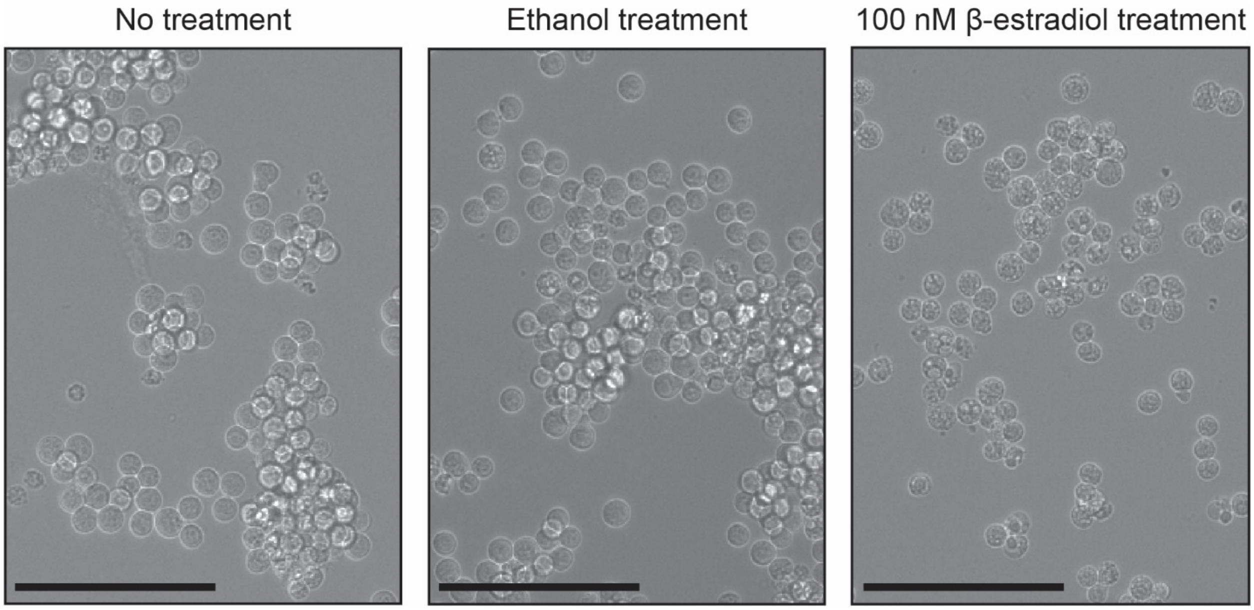


**Figure S11. β-estradiol treatment of G1ERs induce morphological differences of cells after two days in culture.** G1ERs were plated in a 96 well plate and treated with no addition chemical species (left), ethanol vehicle (middle) or 100 nM β-estradiol (right) for two days prior to imaging. β-estradiol treatment is expected to cause differentiation of the G1ER cells into erythroid-like cells under these conditions. We observed that β-estradiol treatment for 48 h reduced the cell count relative to the no treatment or ethanol treatment conditions. Cell count in each well determined by manual counting is reported in the **Supplementary Data 2** for **Figure 5e**. G1ER cells treated with β-estradiol also developed a cellular phenotype characterized by dark puncta which may be a result of cellular or membrane remodeling, consistent with the expected differentiation process. Scale bar is 100 μm.

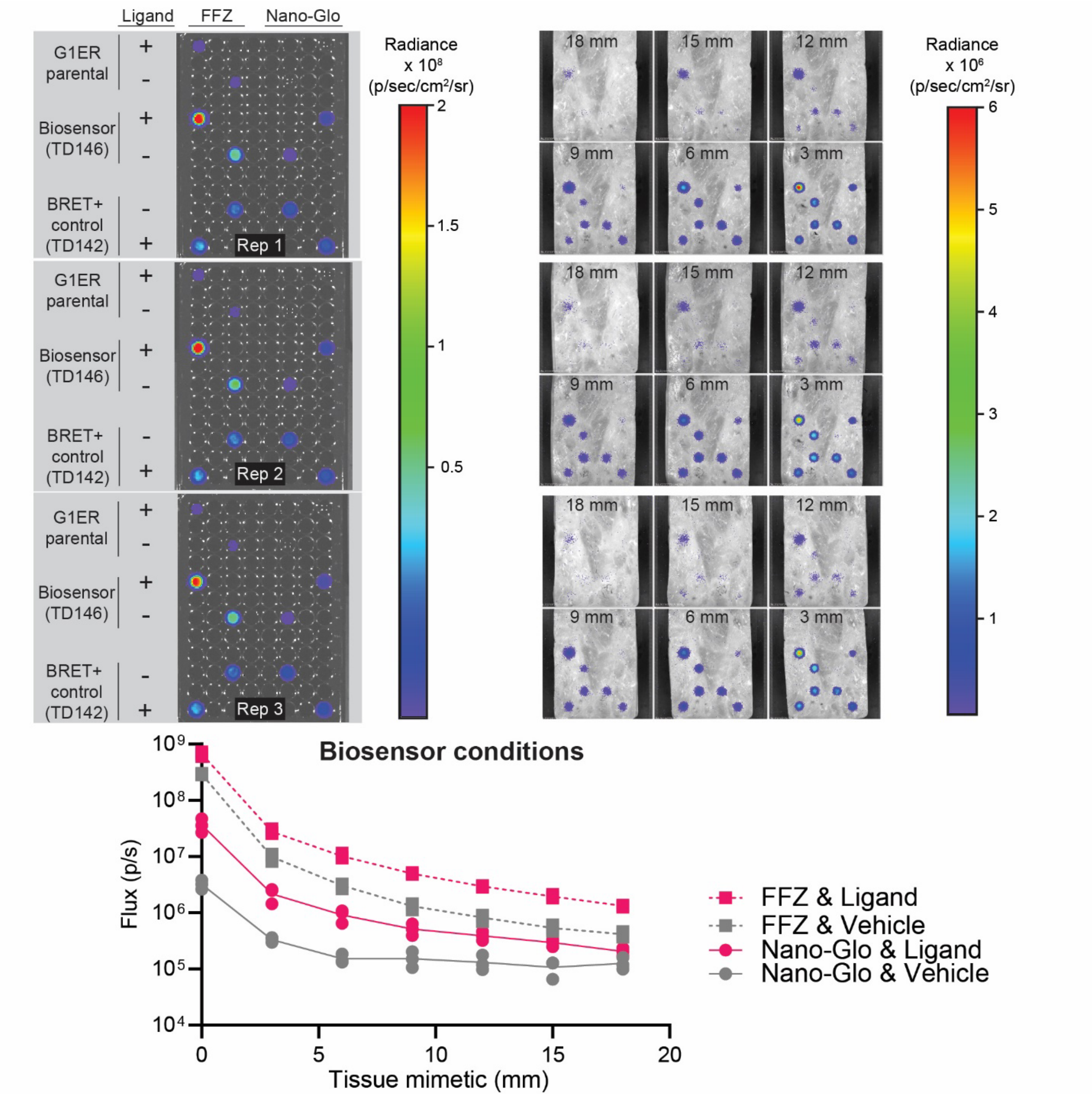


**Figure S12. Uncropped IVIS data from Figure 5f demonstrates that BRET signal from BRET biosensors is brighter when using fluorofurimazine (FFZ) as a substrate instead of furimazine (Nano-Glo). (a, top)** IVIS data showing the luminescent signal from BRET biosensor construct TD146 in G1ER cells through no ham or up to 18 mm of ham. Two substrates were evaluated: FFZ and Nano-Glo. 1 x $10^6$ TD146 BRET biosensor cells were plated and 2 x $10^3$ TD142 BRET positive control cells were plated to facilitate comparison on the same plate. Three replicates are shown on three separate plates. **(a, bottom)** Quantification of the data in the top of the figure, showing that FFZ generated brighter BRET signal through many layers of tissue mimetic. FFZ was selected as the substrate for subsequent *in vivo* studies. The symbols represent the three biological replicates shown, the lines connect the means of the replicates at each tissue thickness, and the error bars represent SEM. The trends are representative of two independent experiments.

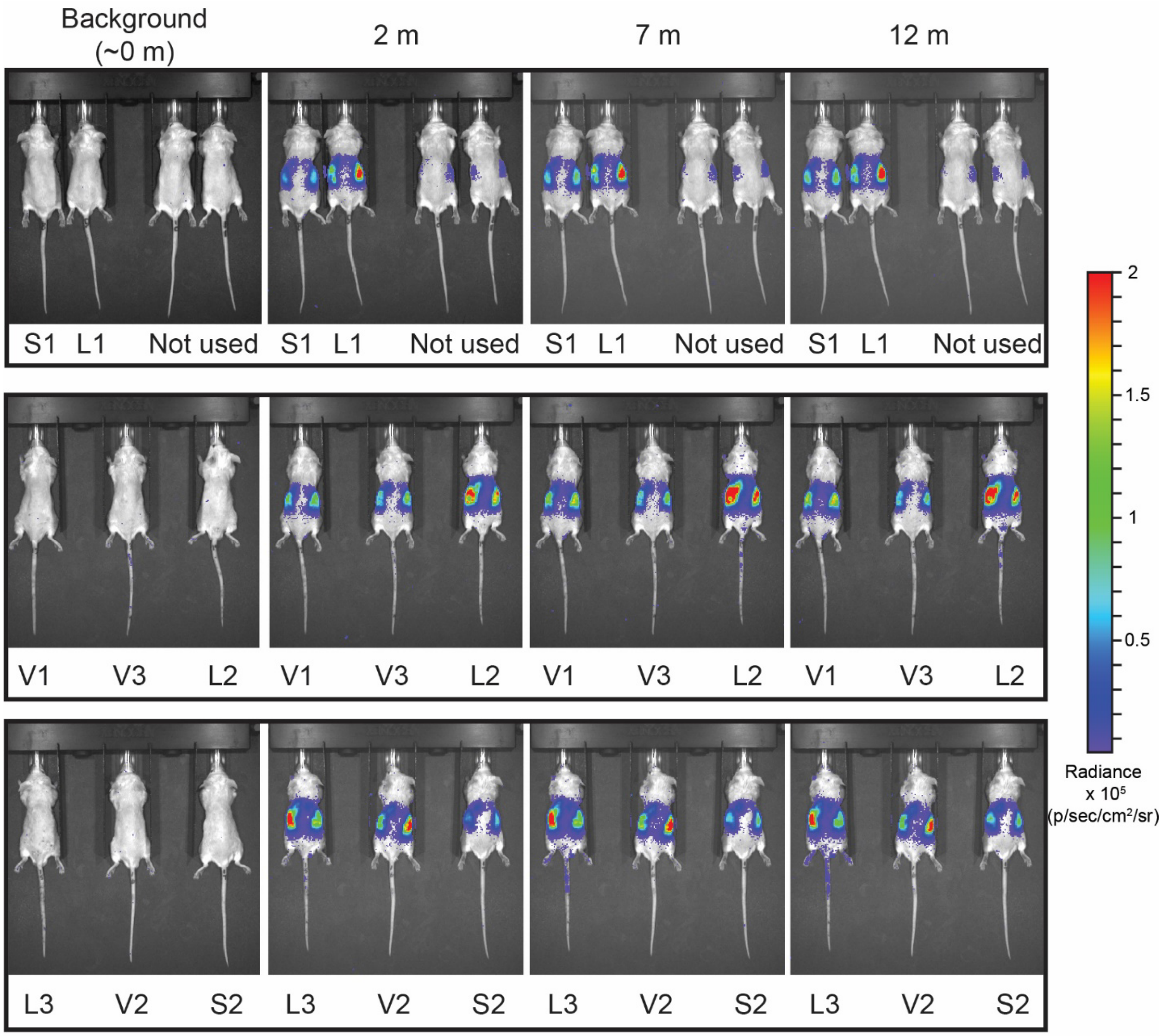


**Figure S13. G1ER cells engineered to express GPA-based biosensors detect cognate ligand in vivo.** Uncropped IVIS data for all mice in this study prior to ("baseline"), 2 m, 7 m, and 12 m after substrate administration. The images above correspond to the data points presented in **Figure 6c-d**. The 12 m time point images are also presented in **Figure 6b**. Luminescence data are represented as a colored heatmap and are scaled equally for all images. Unique mouse identifiers are present below each mouse at each time point and correspond to the identifiers in Figure 6b: Substrate "S", Vehicle "V", and Ligand "L". Mice listed at "Not used" are paired conditions with S1 and L1 (left to right) but were injected with Nano-Glo (diluted 20x in PBS and dosed at 1 mg/kg, IP) instead of fluorofurimazine as an initial evaluation to determine whether Nano-Glo was a suitable substrate in this study. The Nano-Glo signal was weak and therefore not pursued further.

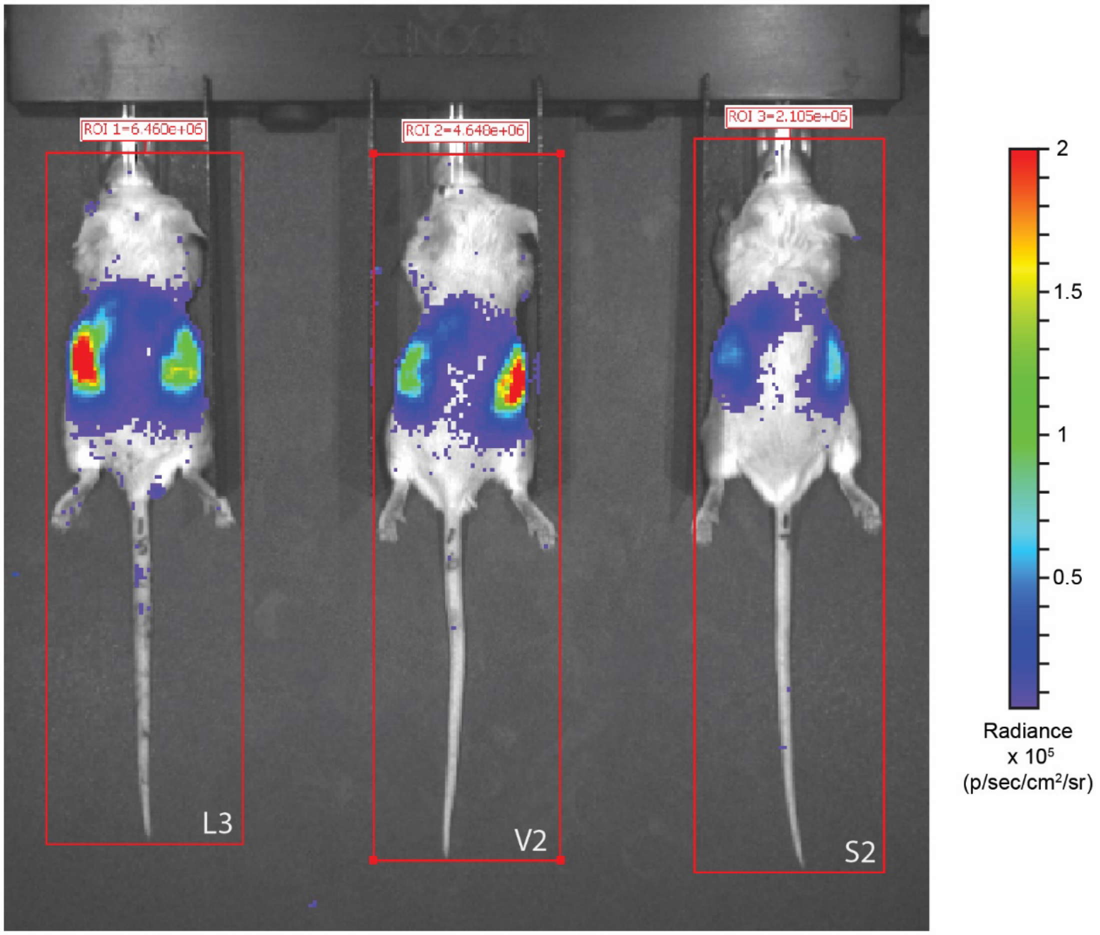


**Figure S14. Drawing a region of interest (ROI) around the entire body of each enabled quantification of the luminescent signal from each animal.** Sample in vivo imaging system (IVIS) data to demonstrate how the ROI was drawn to quantify luminescent signal from each mouse. The red box represents the ROI, and the numbers above each box represent the flux (photons per second) through the ROI. The colored heatmap presents luminescence data in units of radiance. With the exception of the ROI box, these images are identical to those presented in Figure S13 at t = 2 m for animals L3 (ligand #3), V2 (vehicle #2), and S2 (substrate #2). The unique identifiers for each animal are provided below each animal.
