## Supplementary figures and images for "Building synthetic biosensors using red blood cell proteins"

### 1-5s-1ham.png

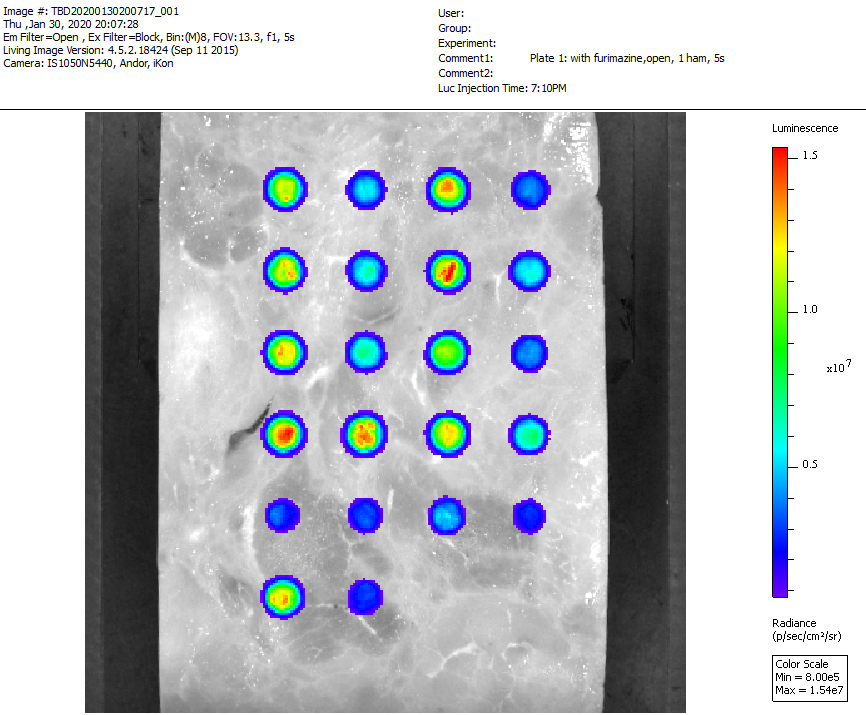

### 1-5s-2ham.png

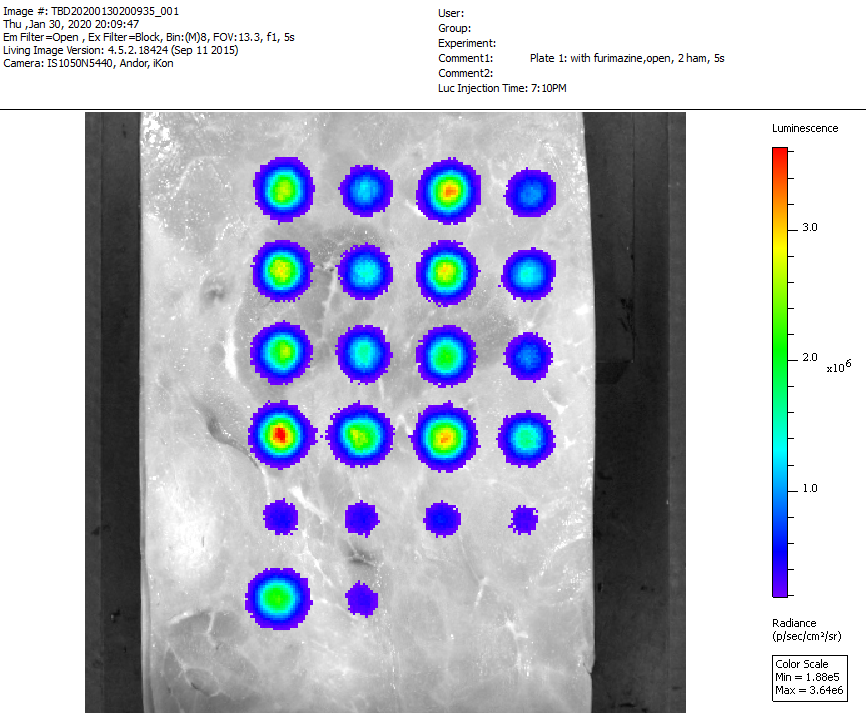

### 1-5s-3ham.png

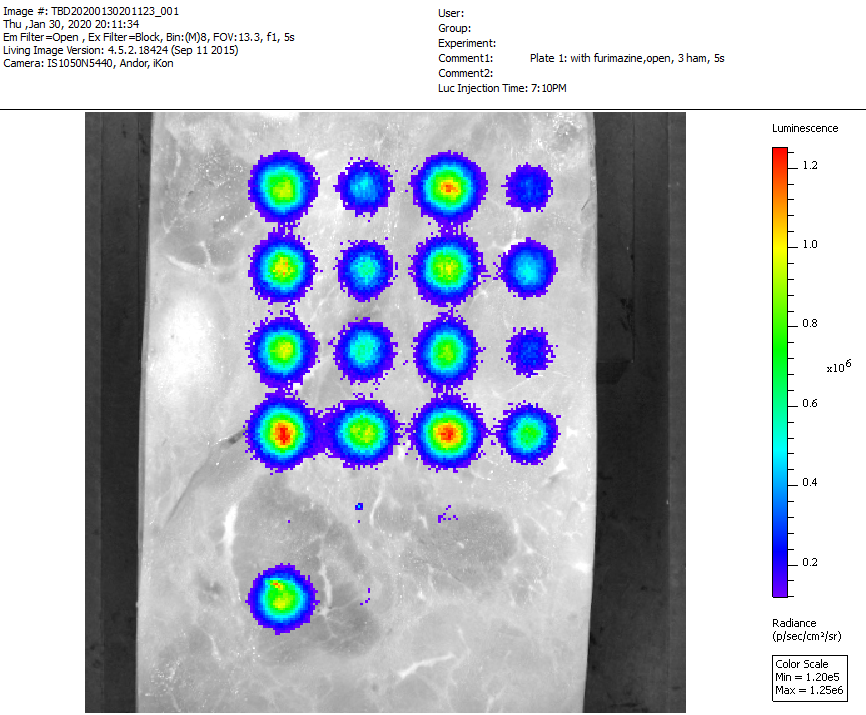

### 1-5s-4ham.png

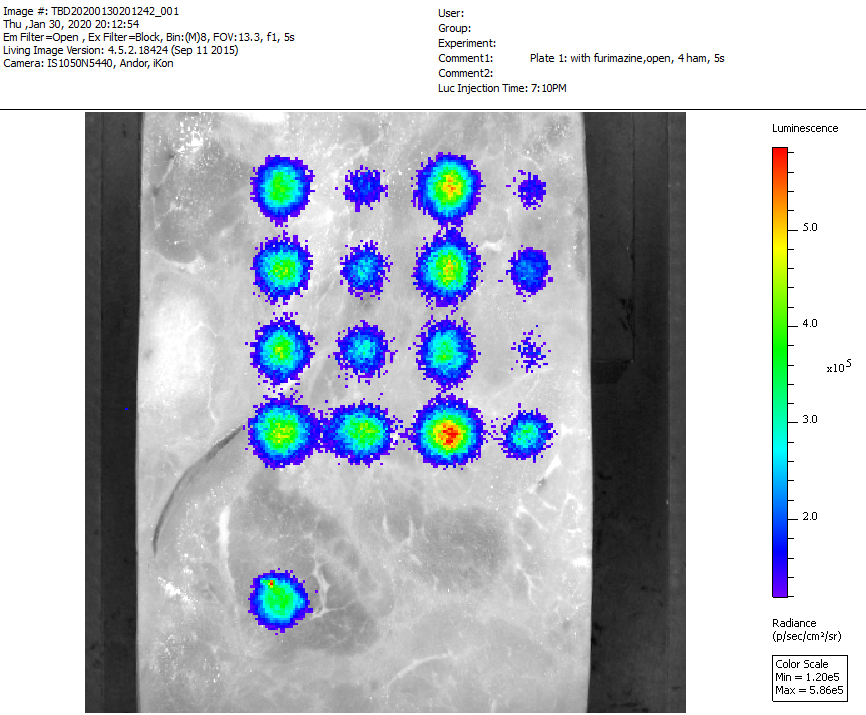

### 1-5s-5ham.png

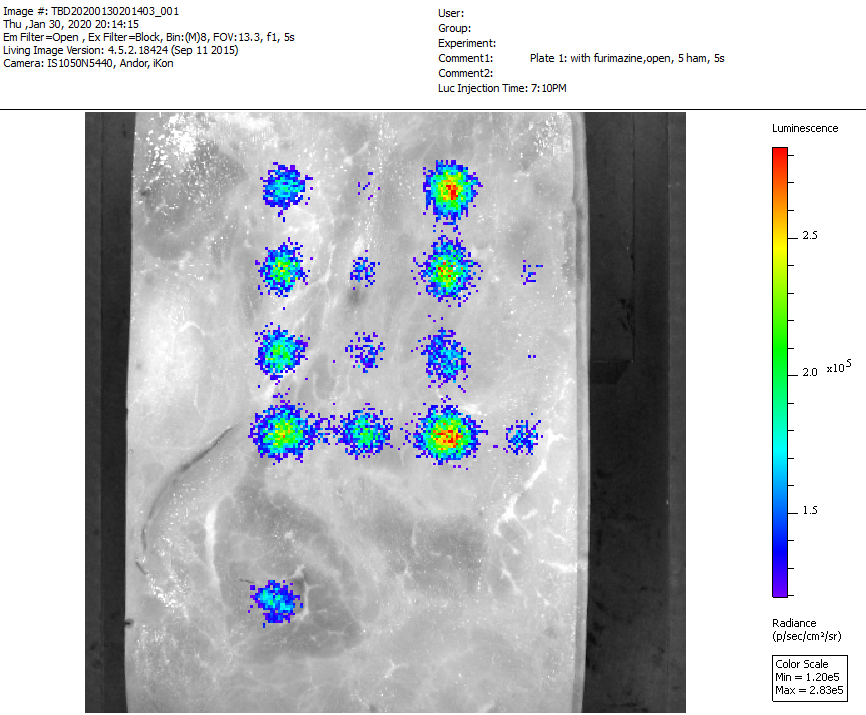

### 1-5s-6ham.png

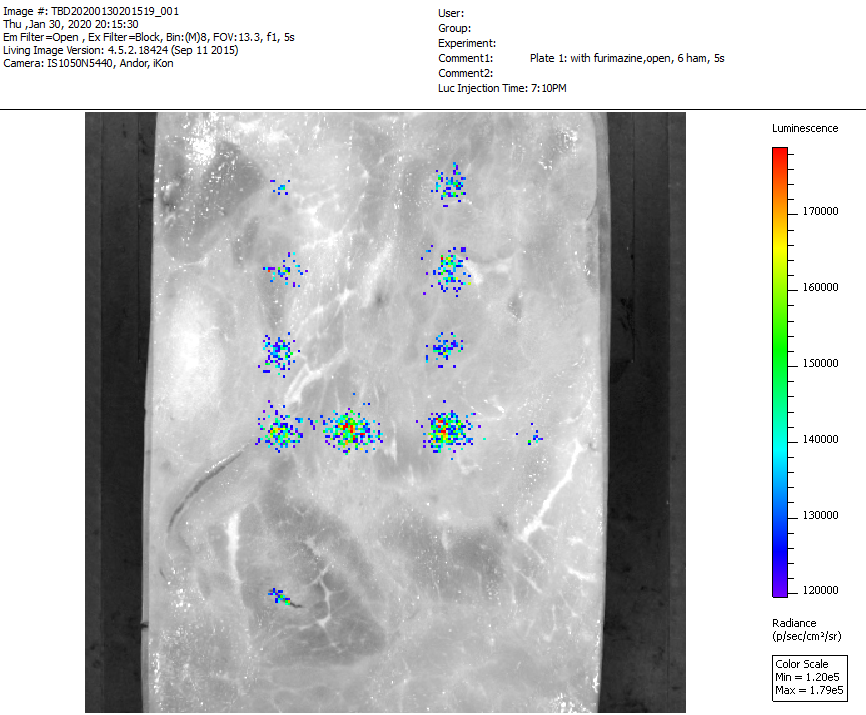

### 1-5s-7ham.png

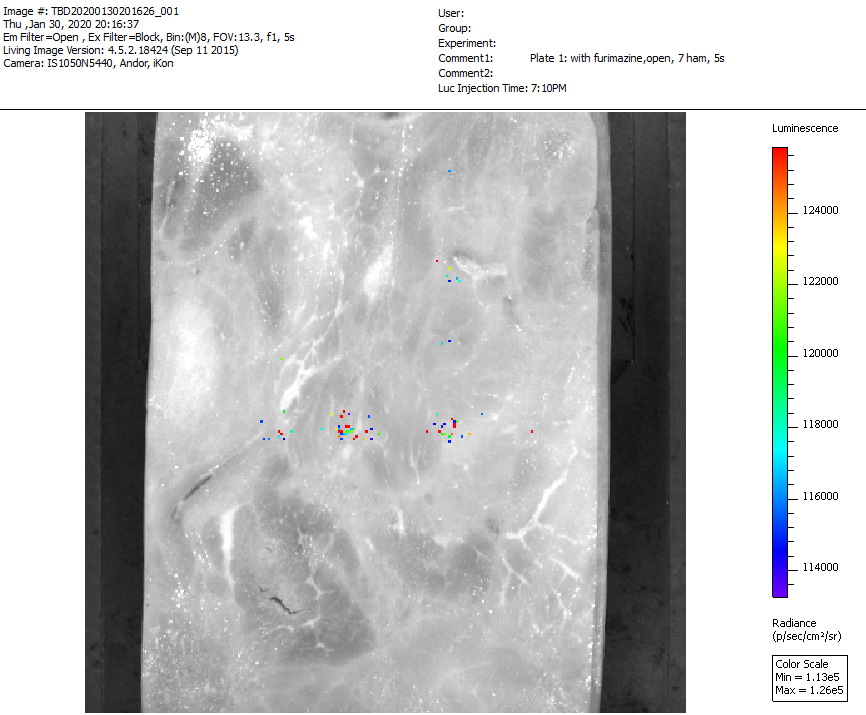

### 2-5s-1ham.png

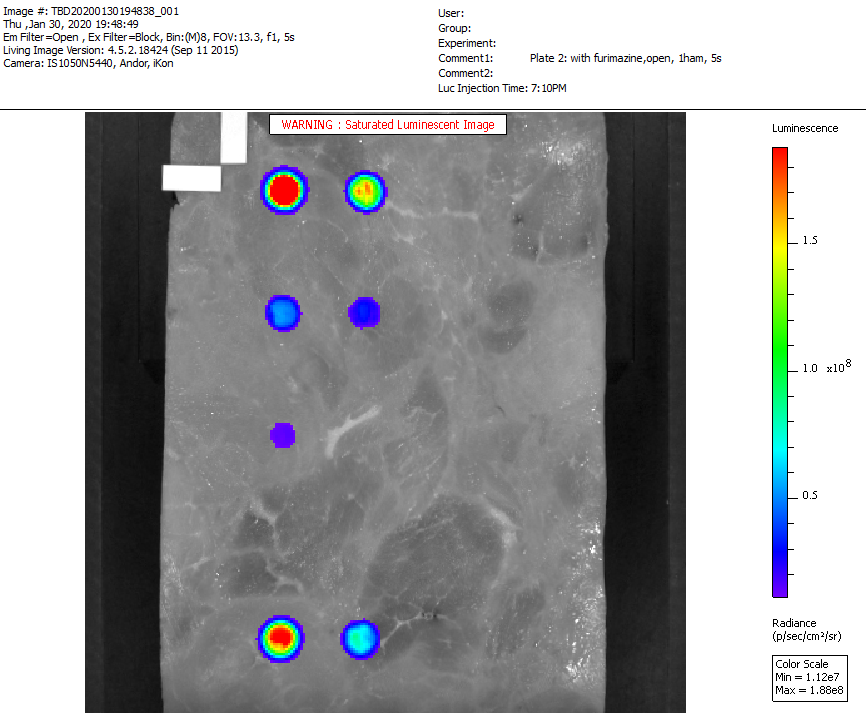

### 2-5s-2ham.png

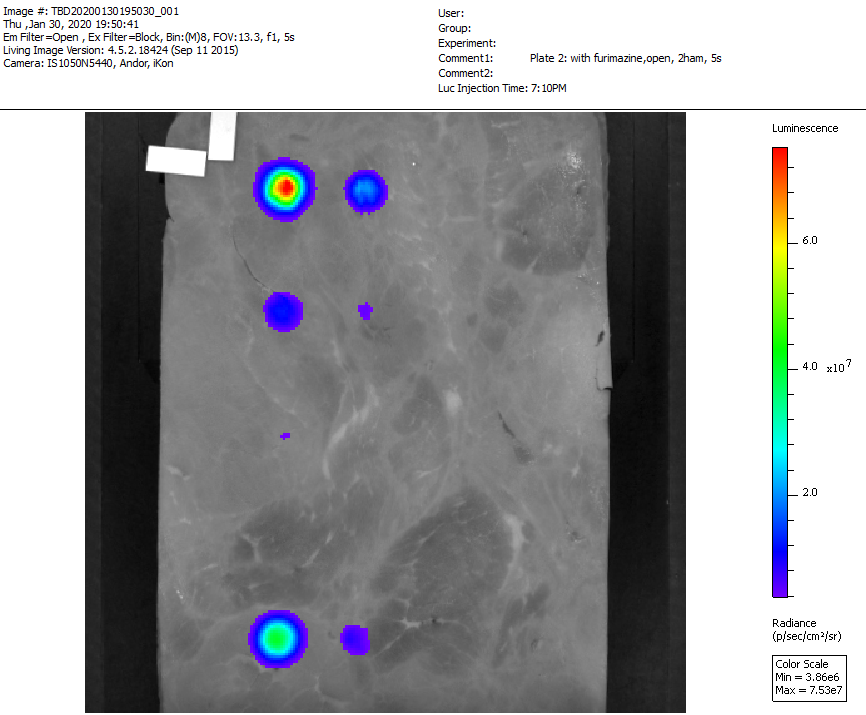

### 2-5s-3ham.png

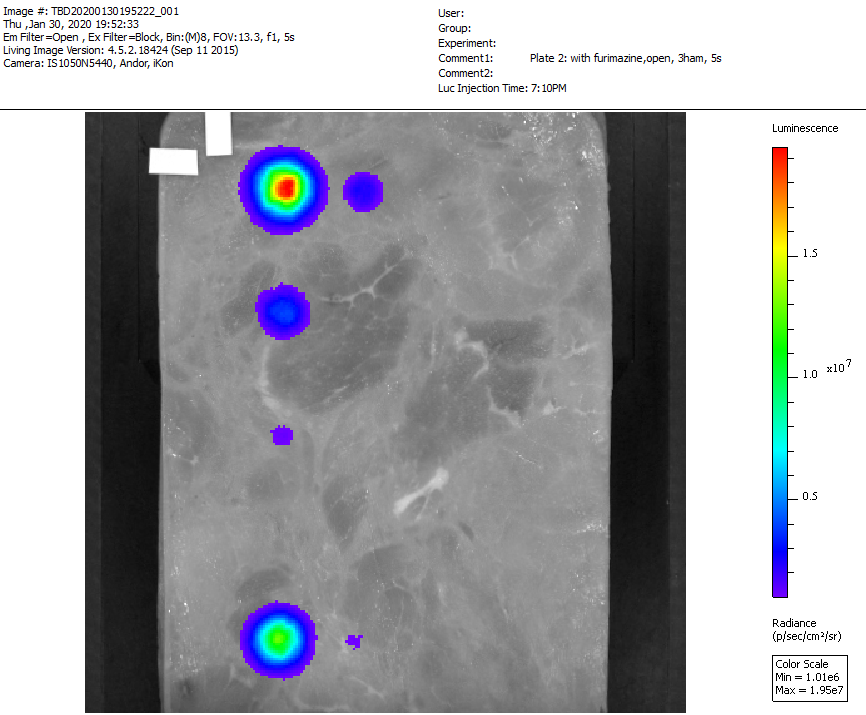

### 2-5s-4ham.png

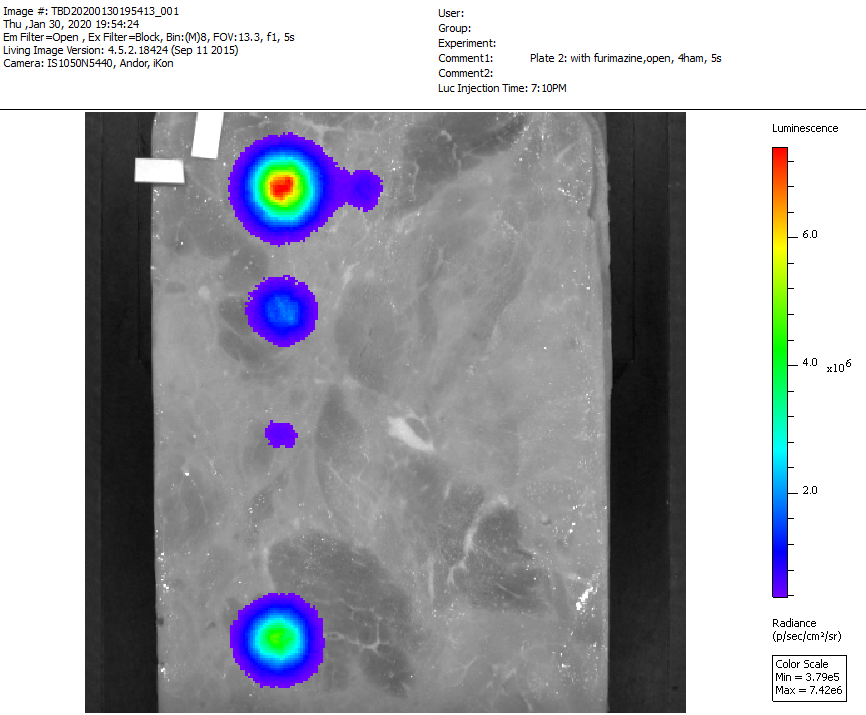

### 2-5s-5ham.png

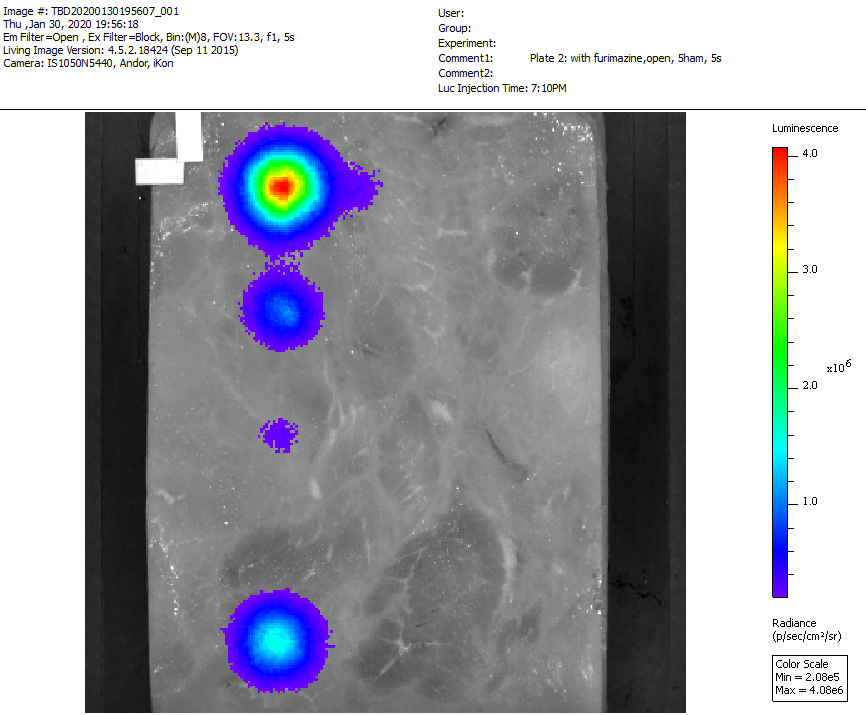

### 8-9-10-11_2m.png

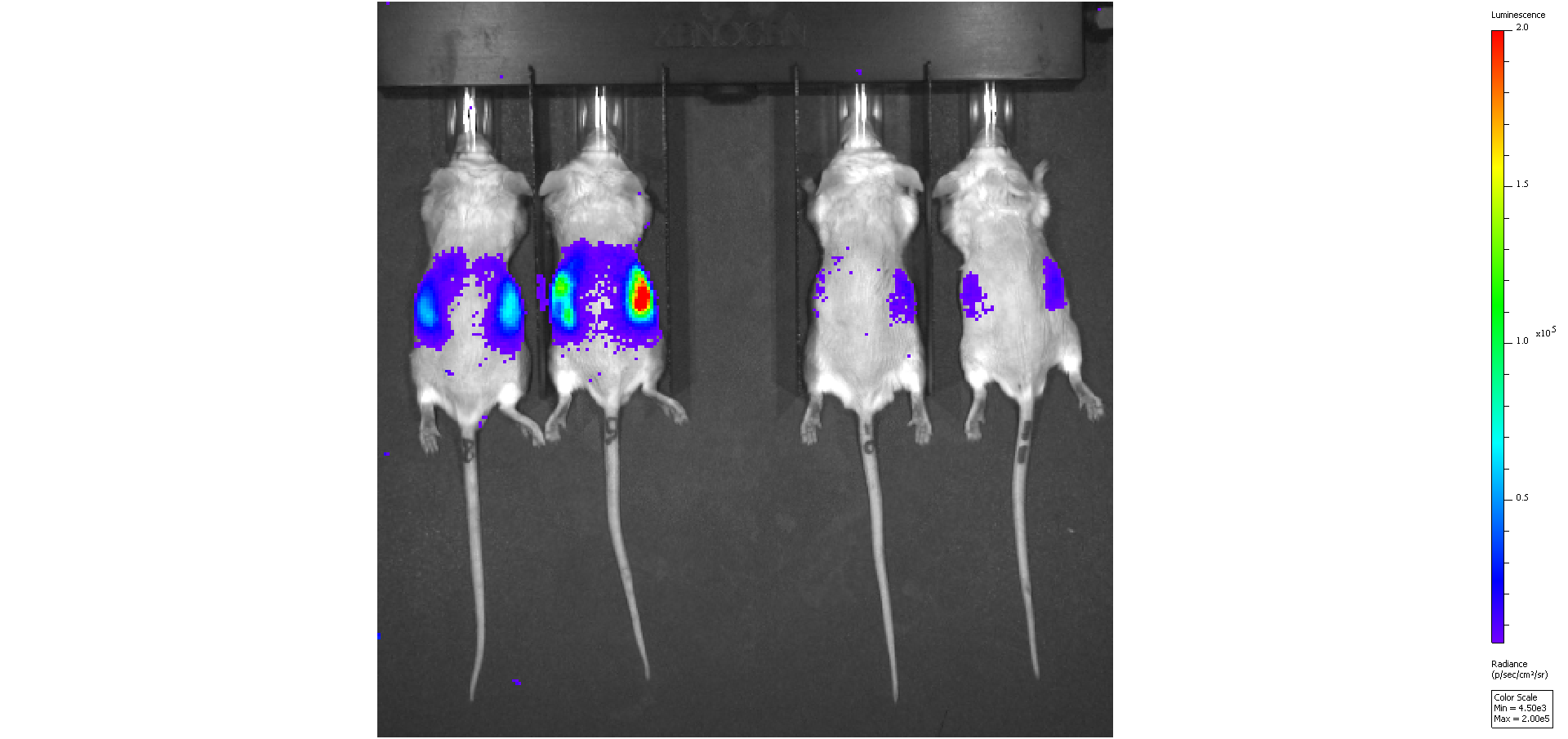

### 8-9-10-11_7m.png

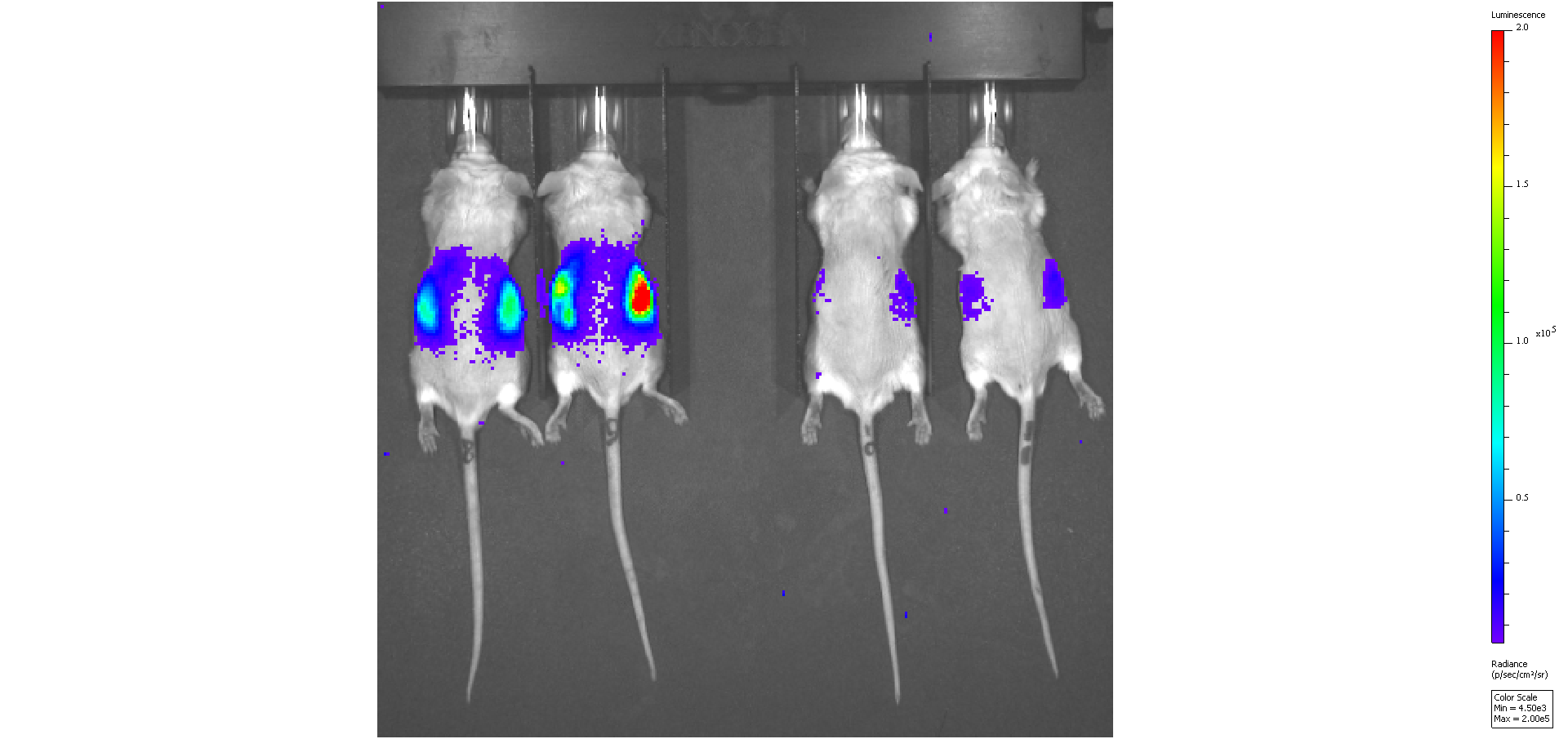

### 8-9-10-11_12m.png

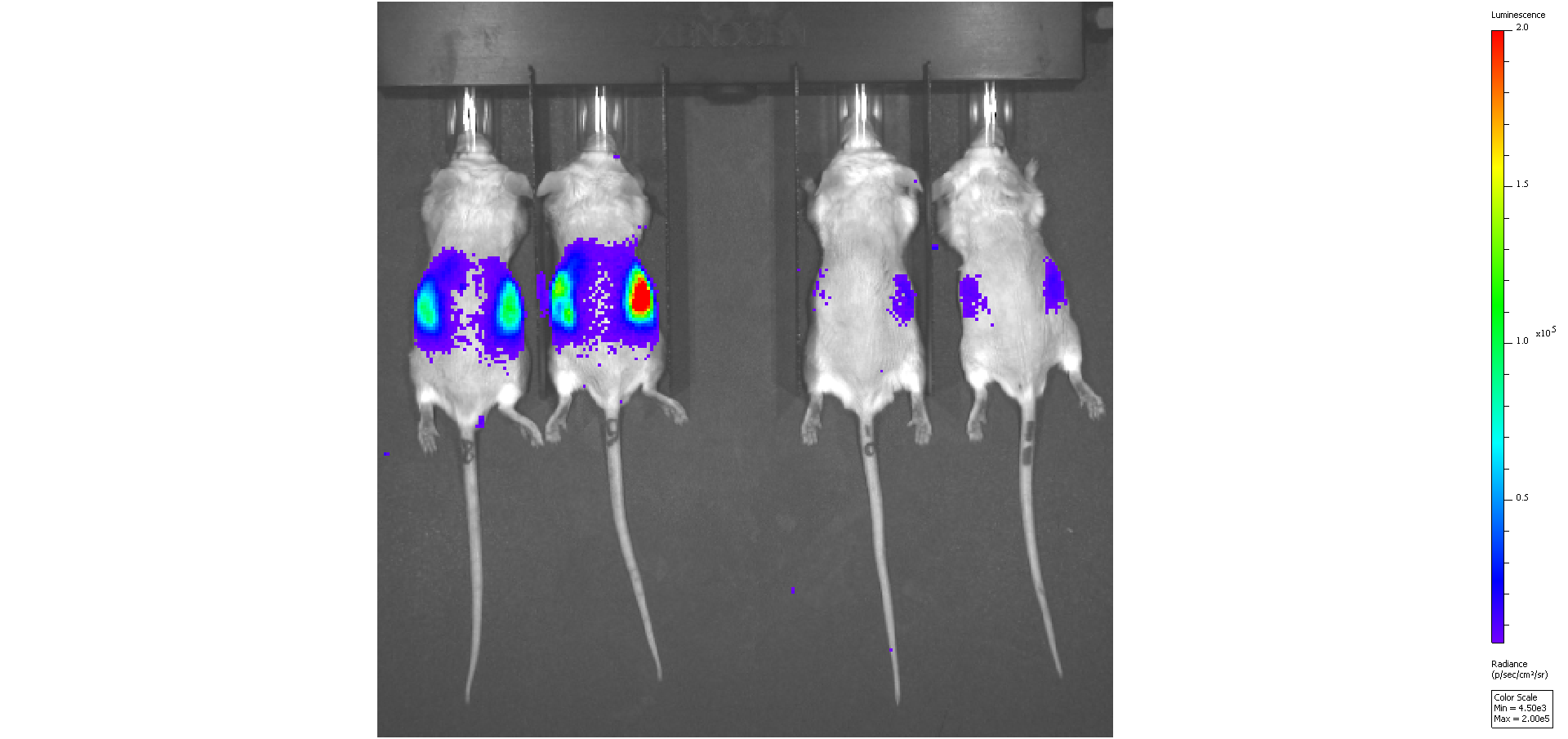

### 8-9-10-11_Baseline.png

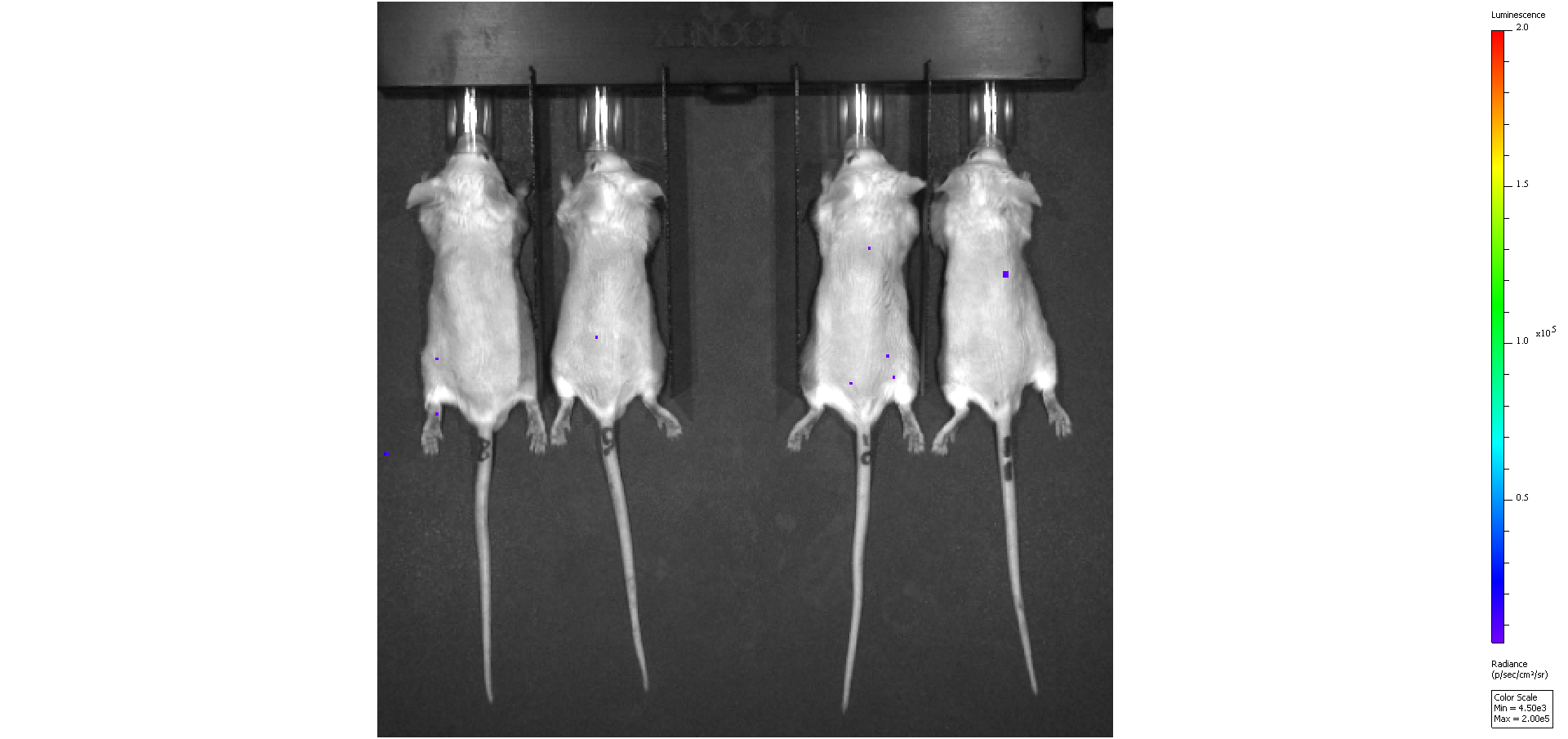

### 12-13-14_2m.png

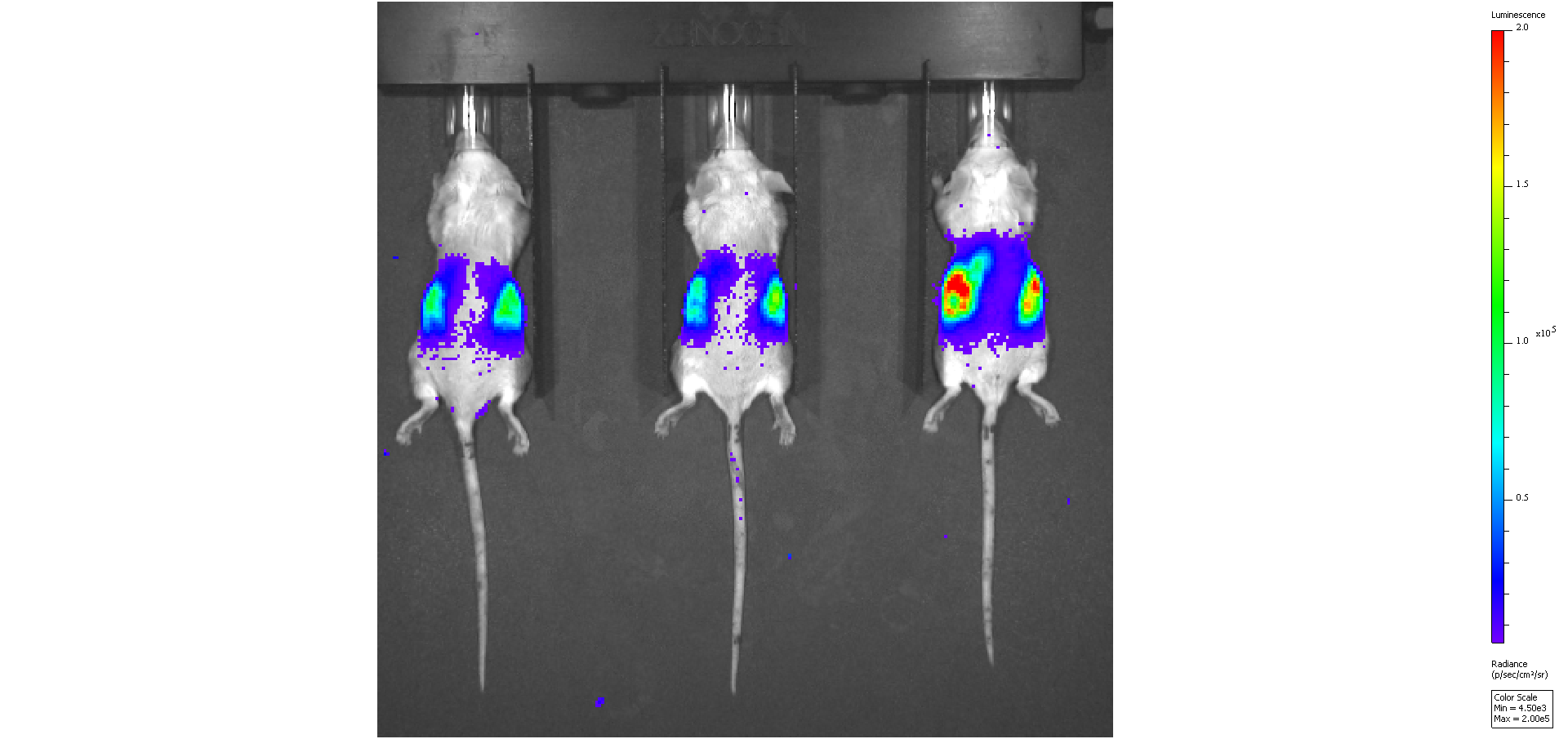

### 12-13-14_7m.png

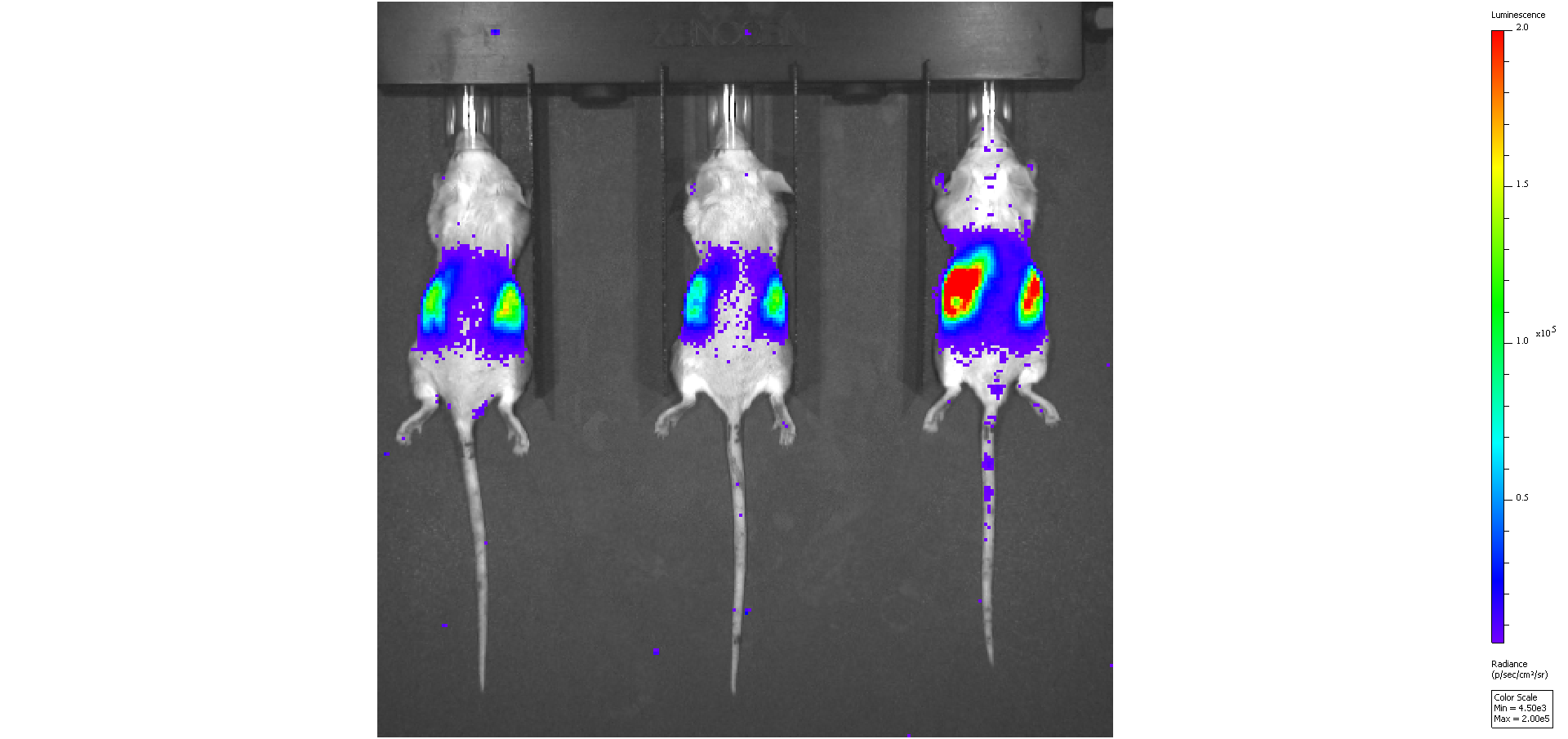

### 12-13-14_12m.png

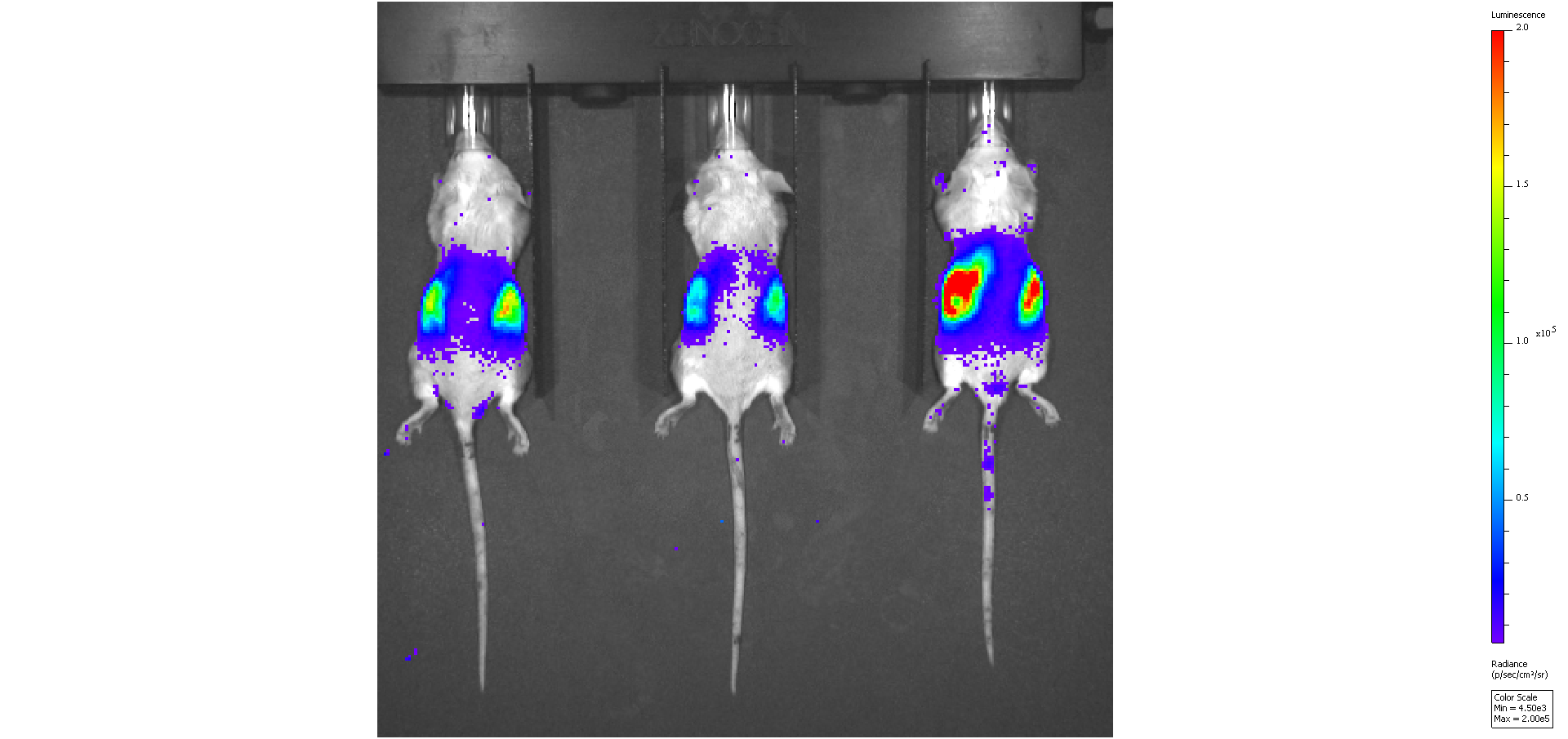

### 12-13-14_Baseline.png

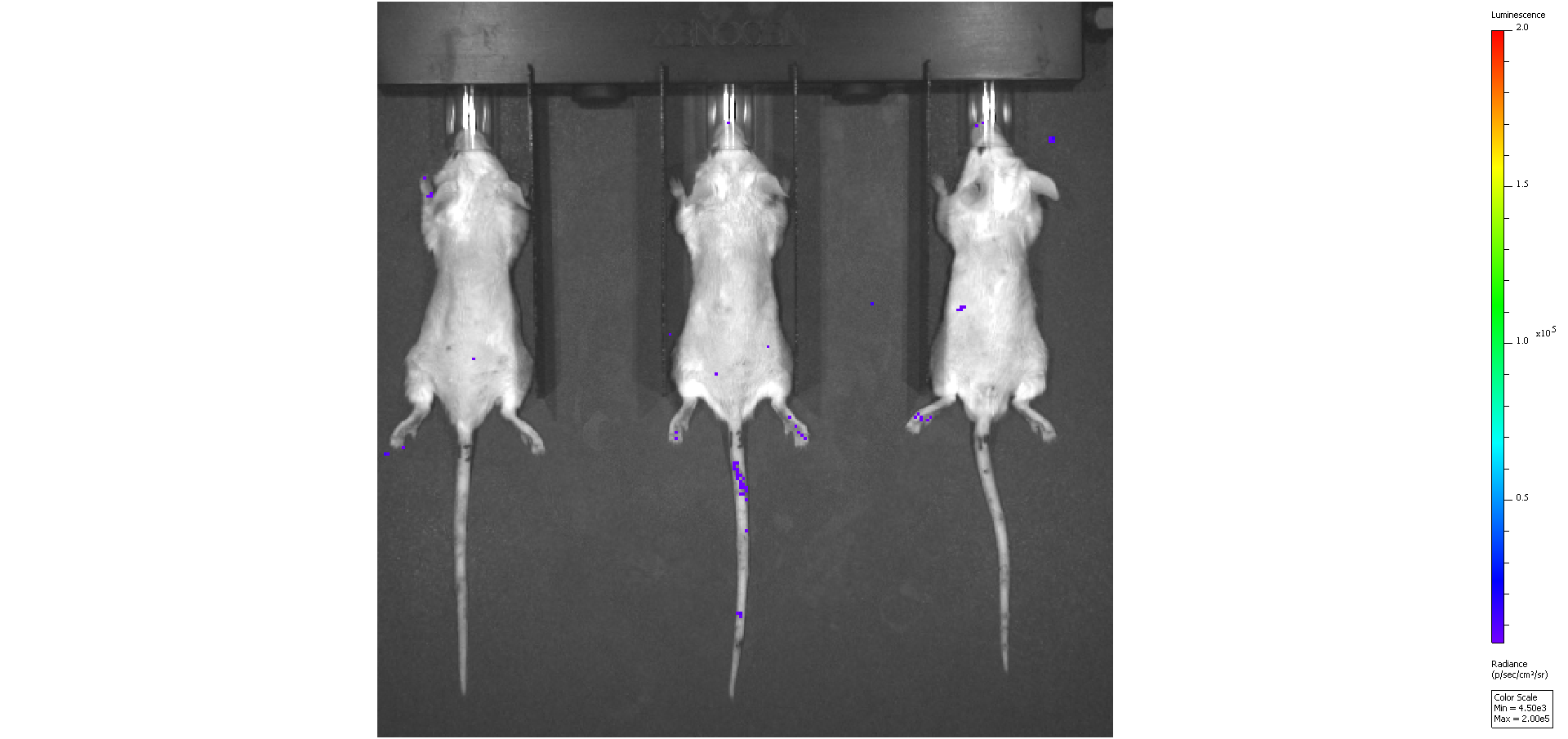

### 15-16-17_2m.png

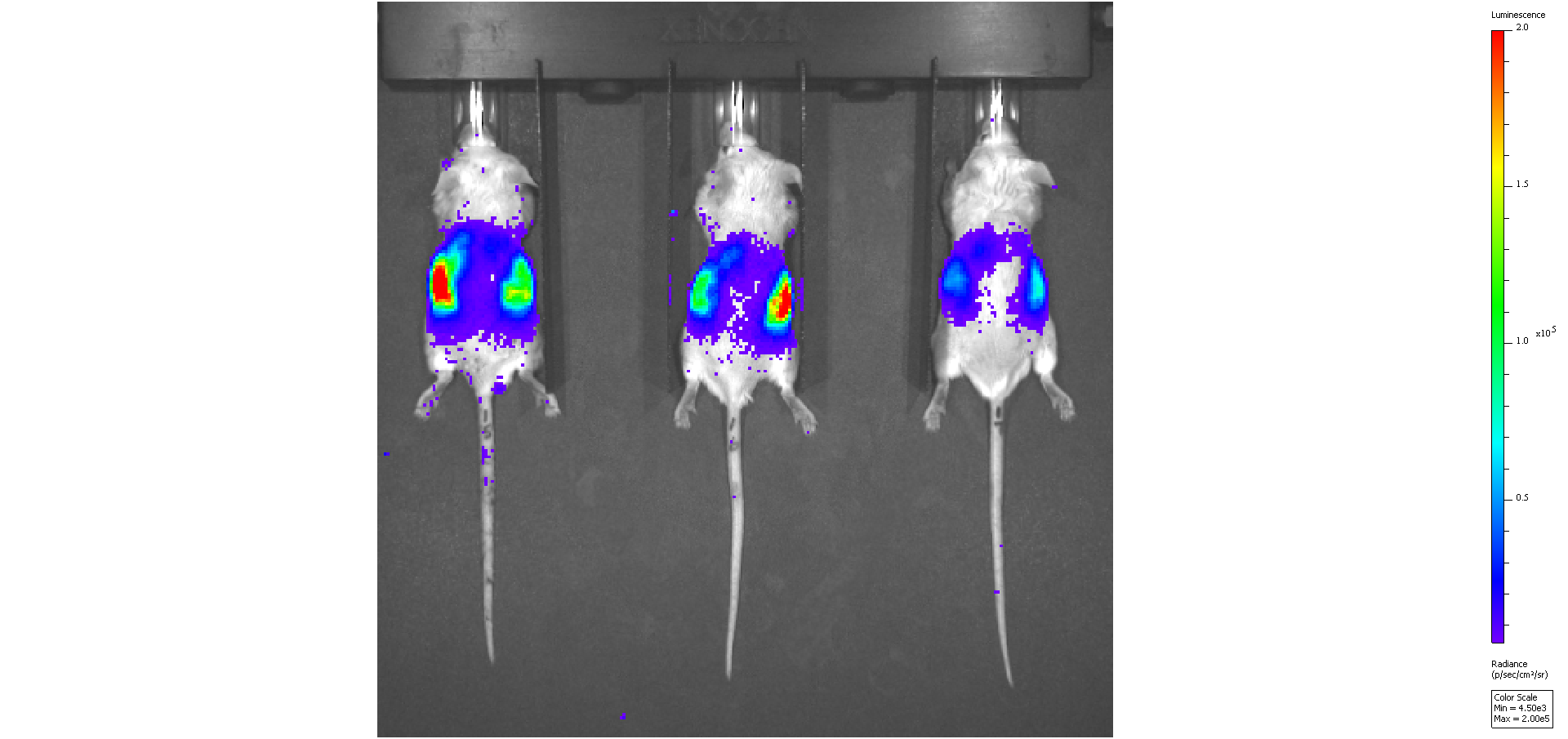

### 15-16-17_7m.png

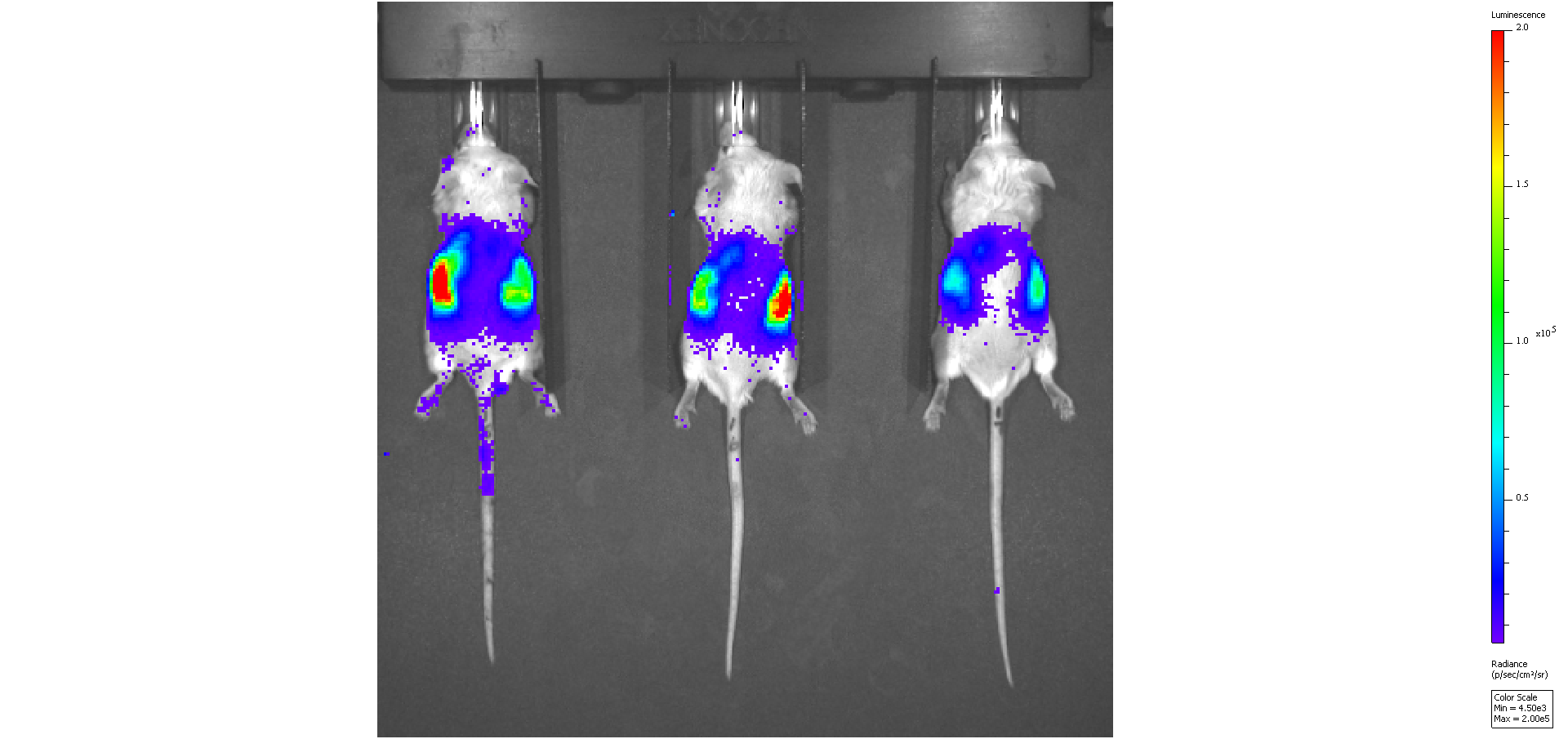

### 15-16-17_12m.png

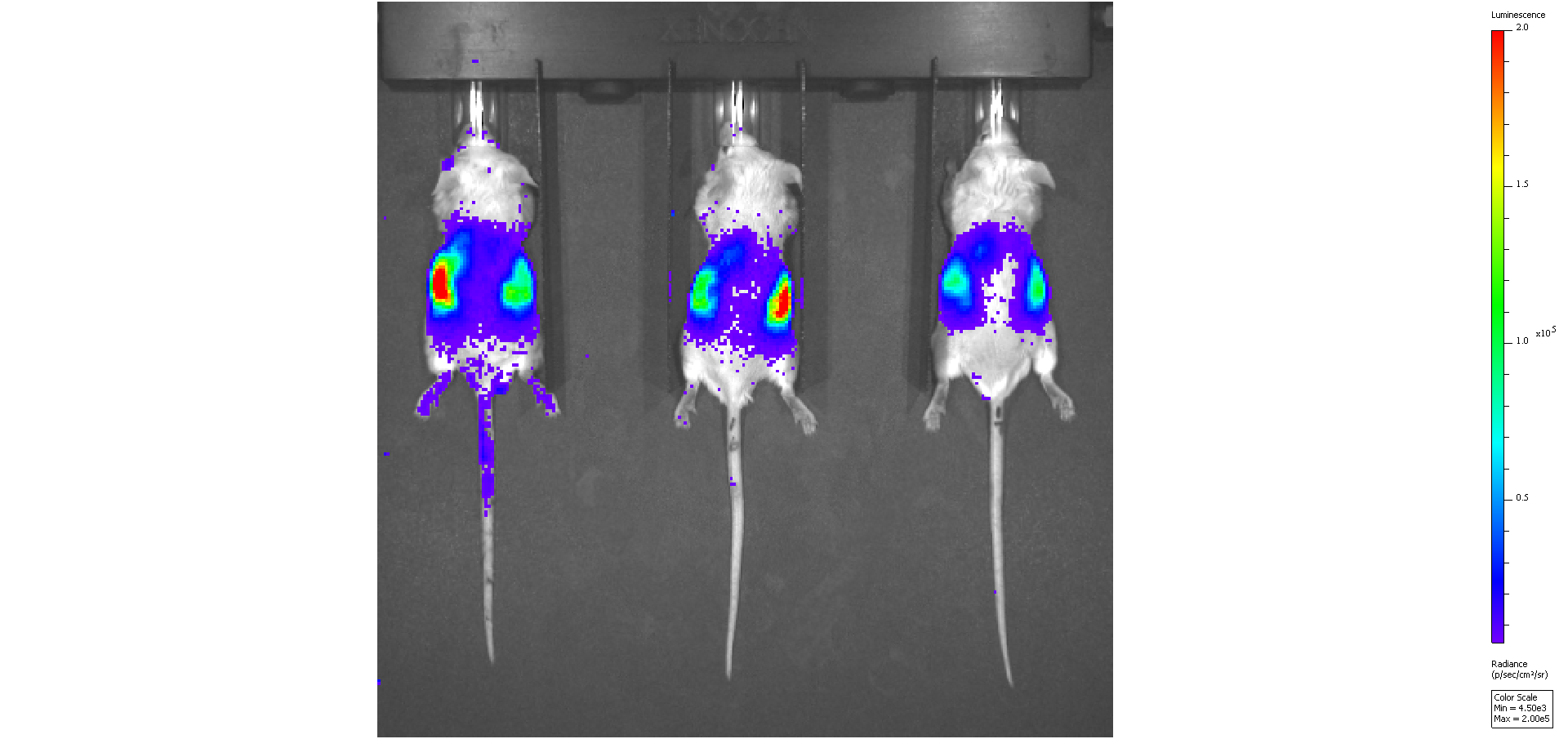

### 15-16-17_Baseline.png

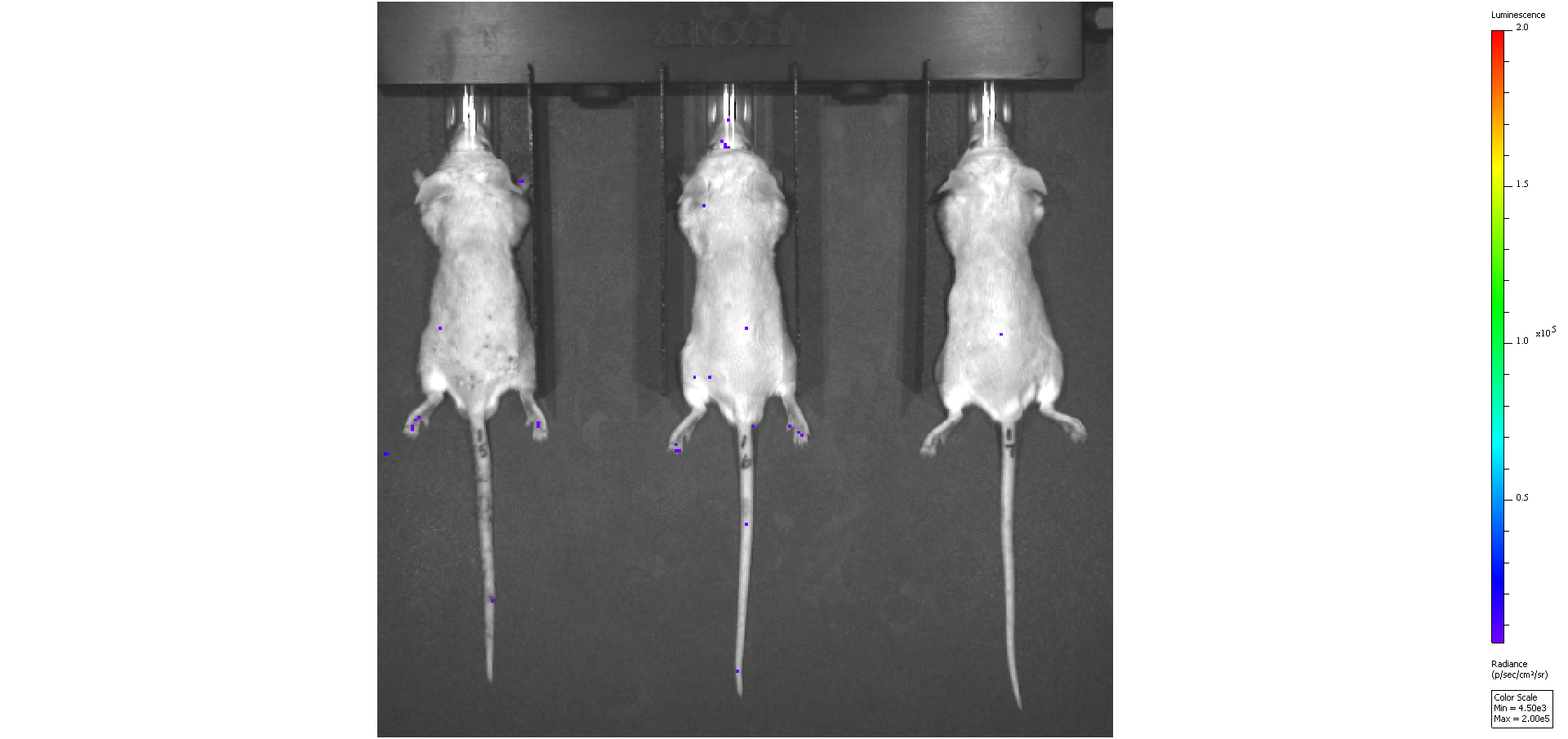

### S5b-Western 3.15.16 b&w.pdf

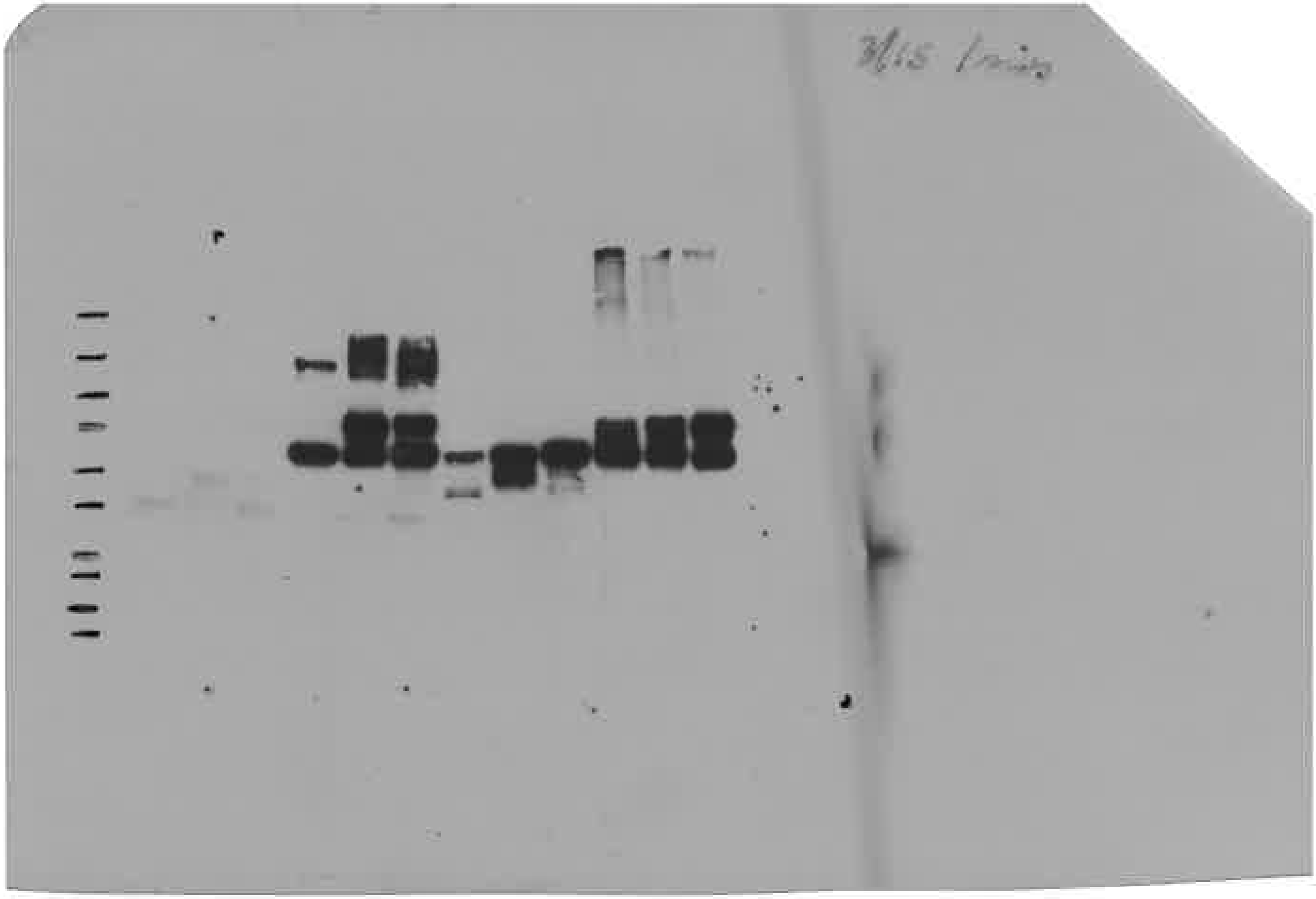

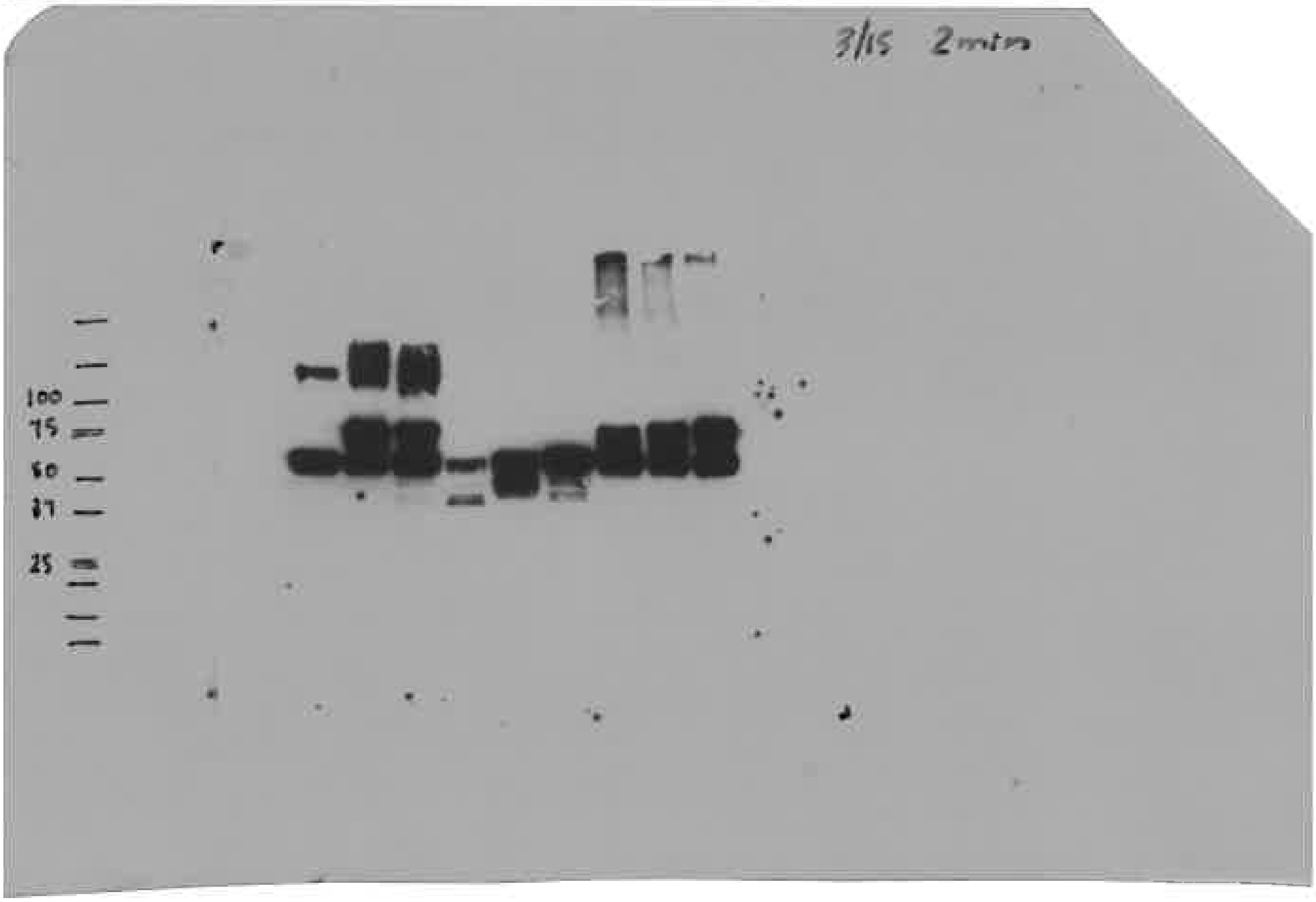

### S5b-Western 3.15.16 color.pdf

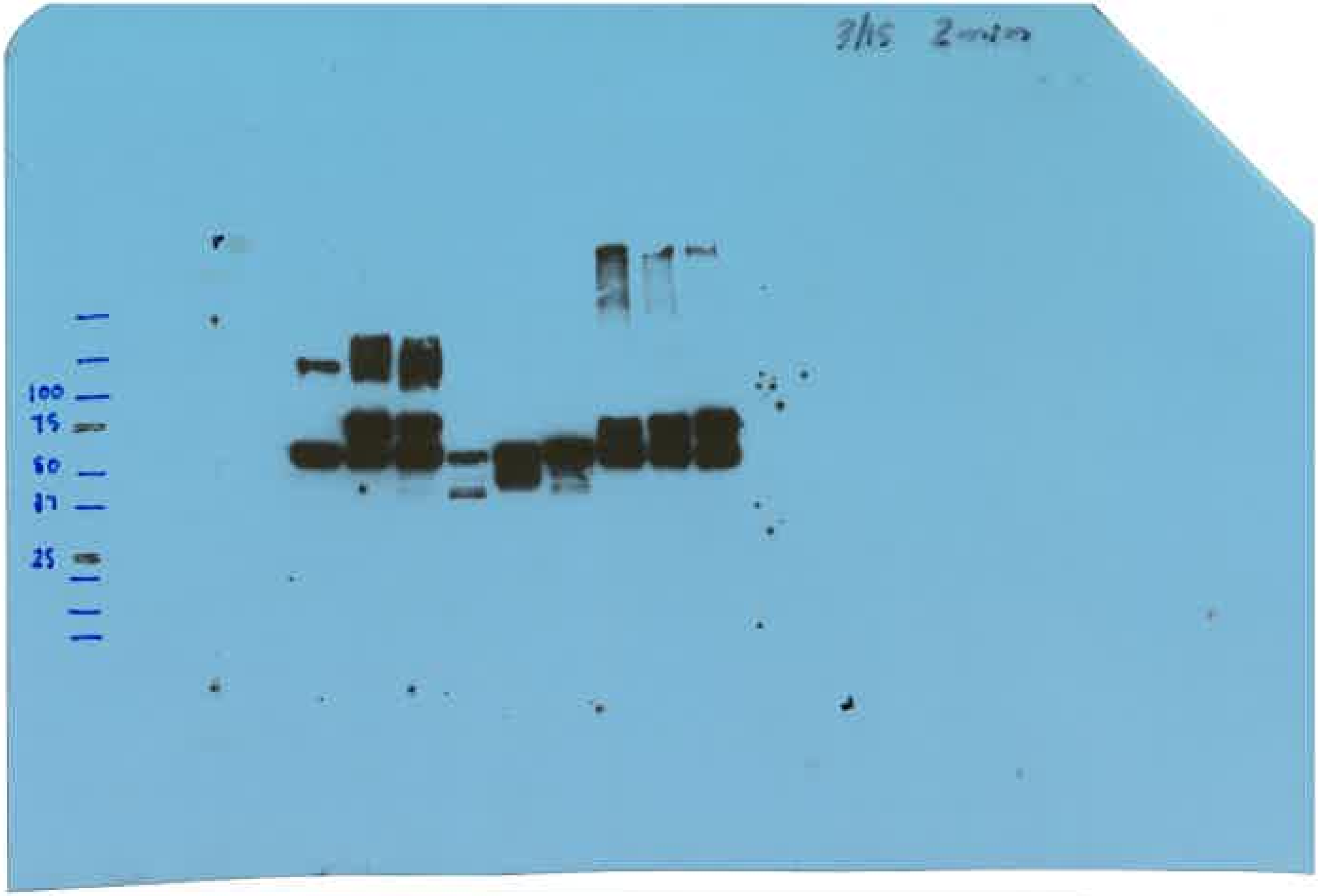

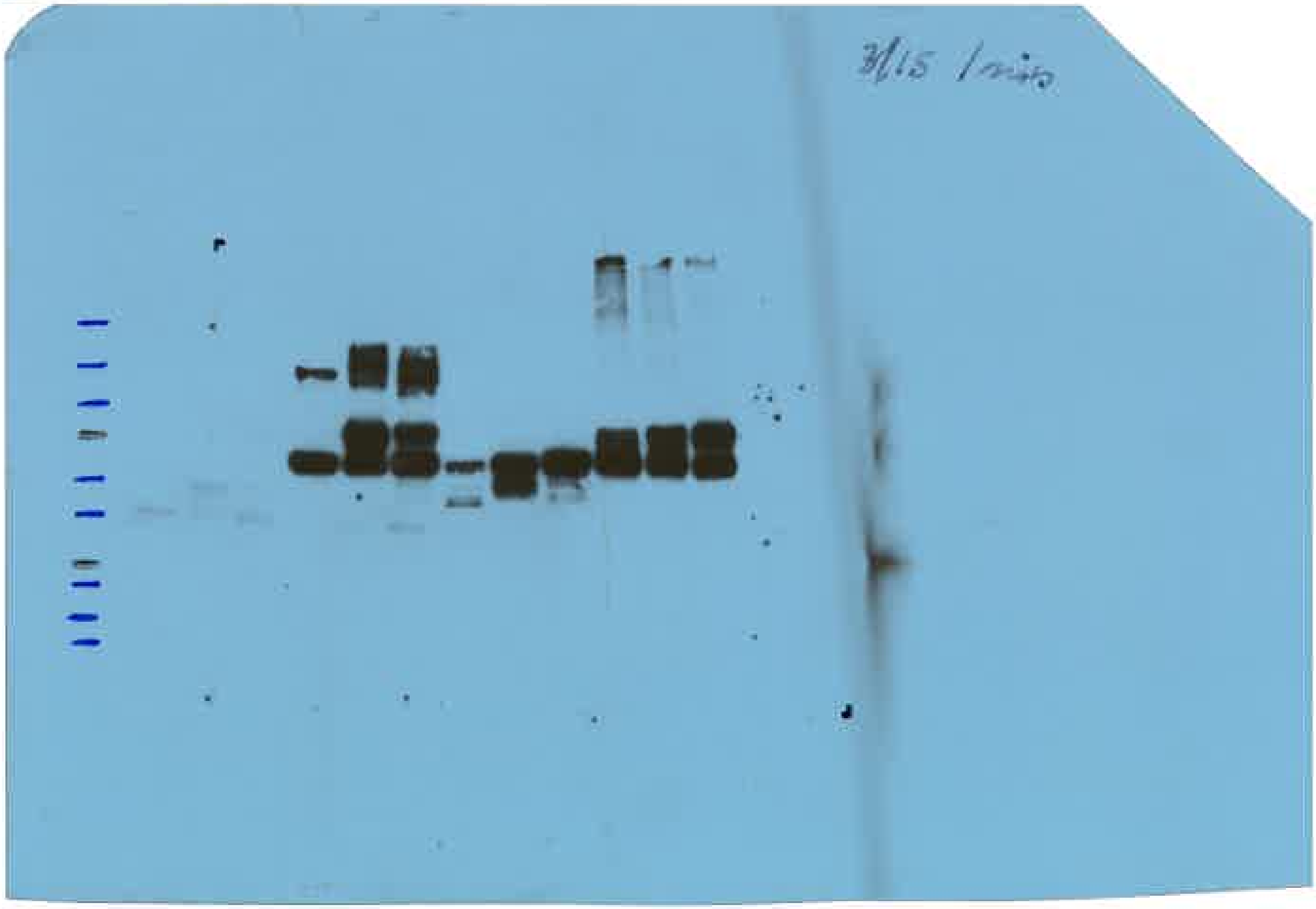

### S9b-Western-S12F5-0316-060654.tif

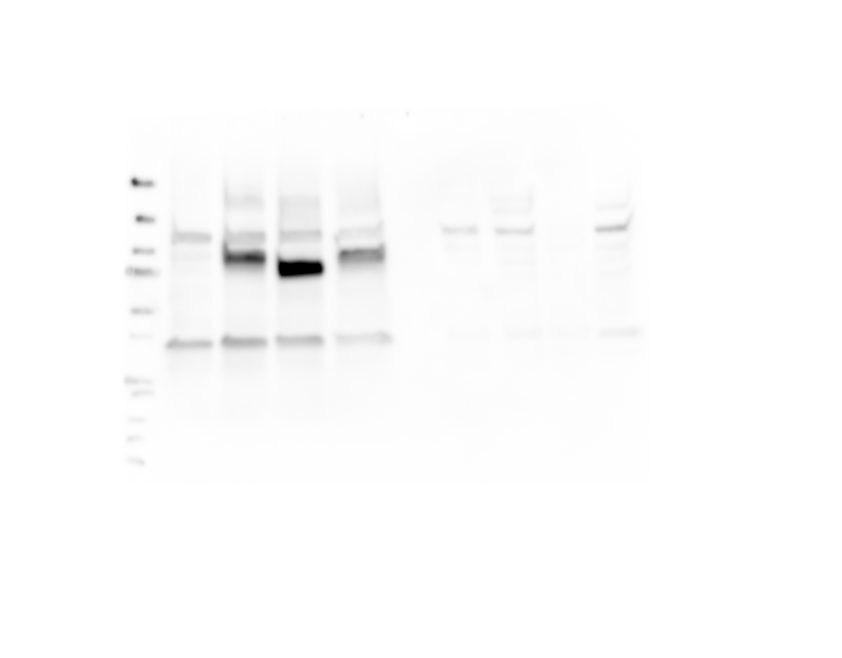

### S9e-0ham.png

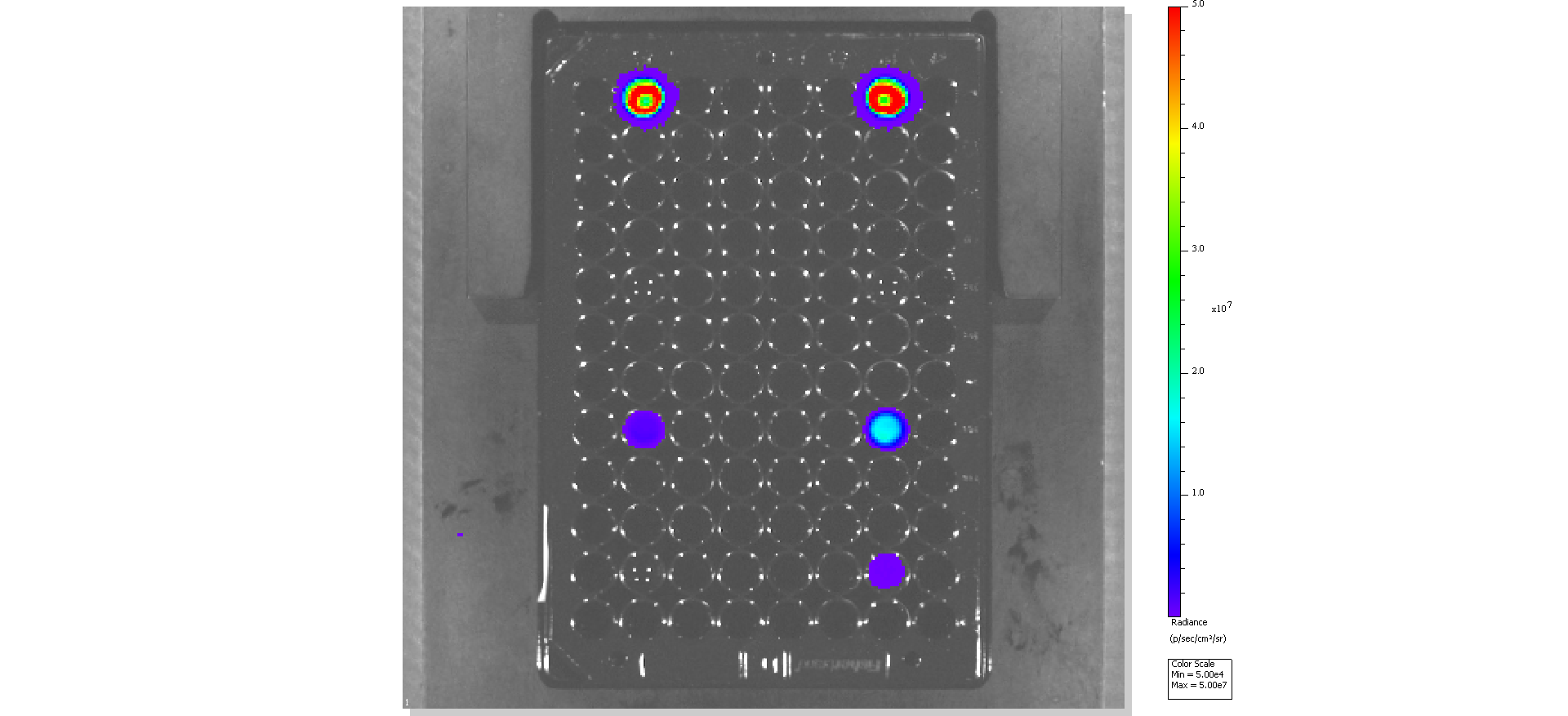

### S10-0ham.png

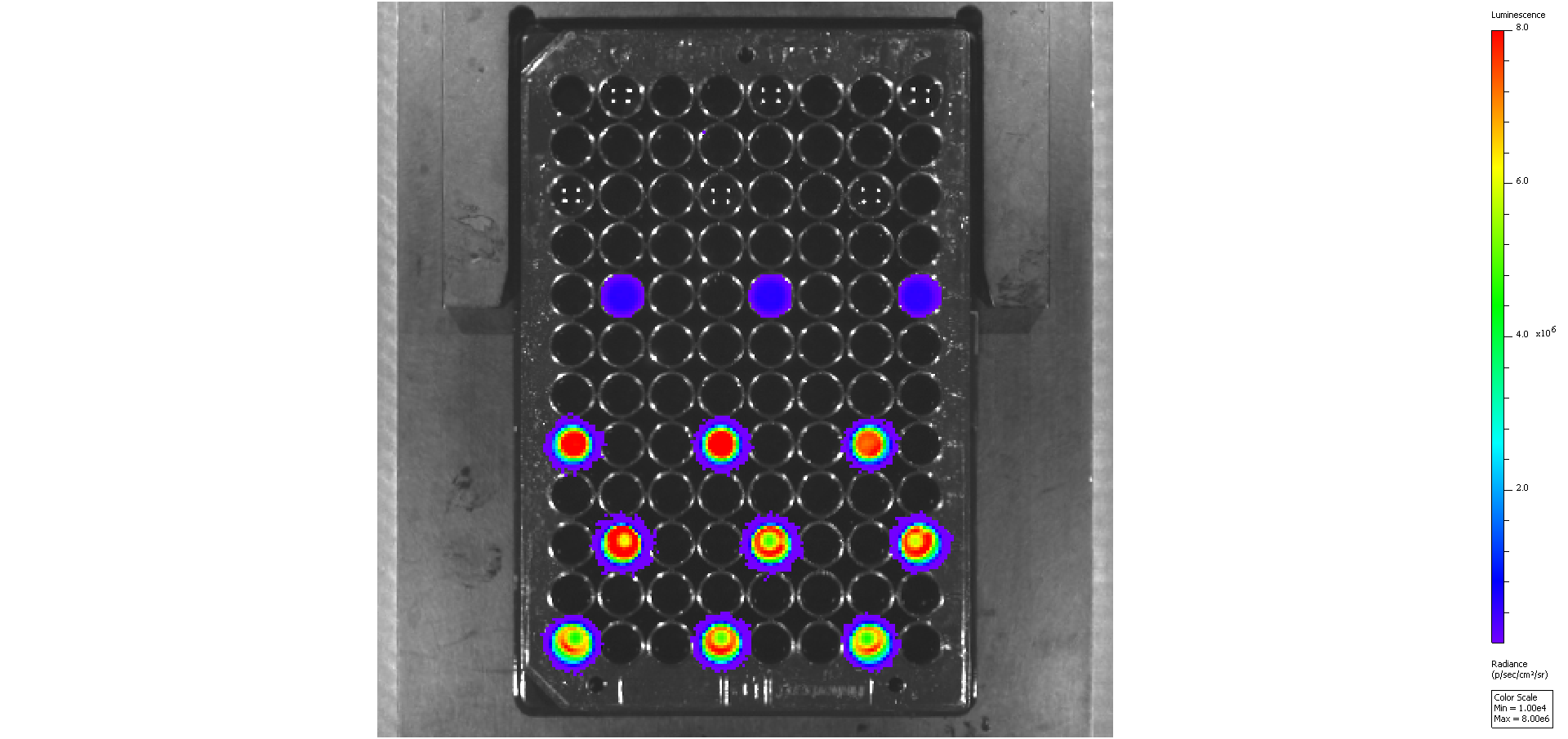

### S10-allham.png

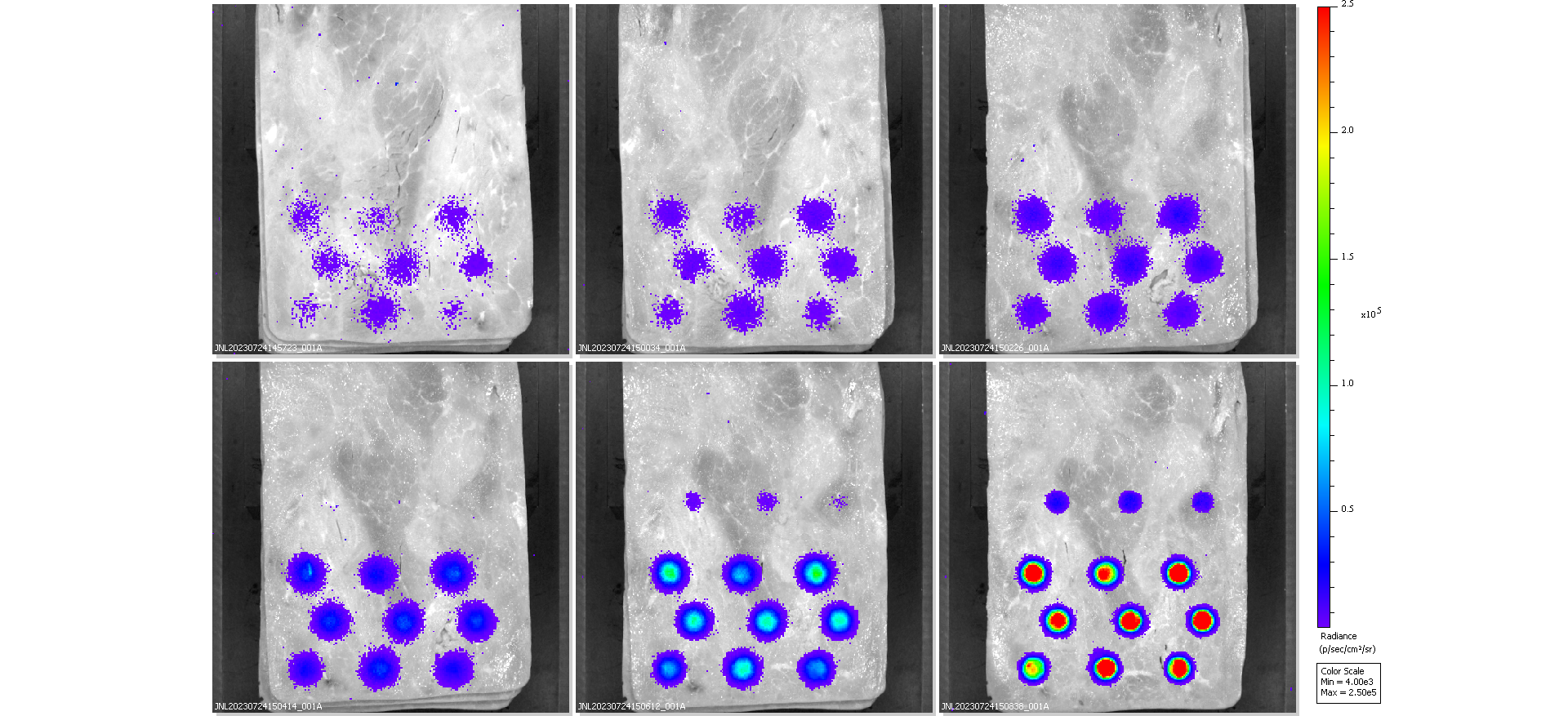

### s12-rep1-0ham.png

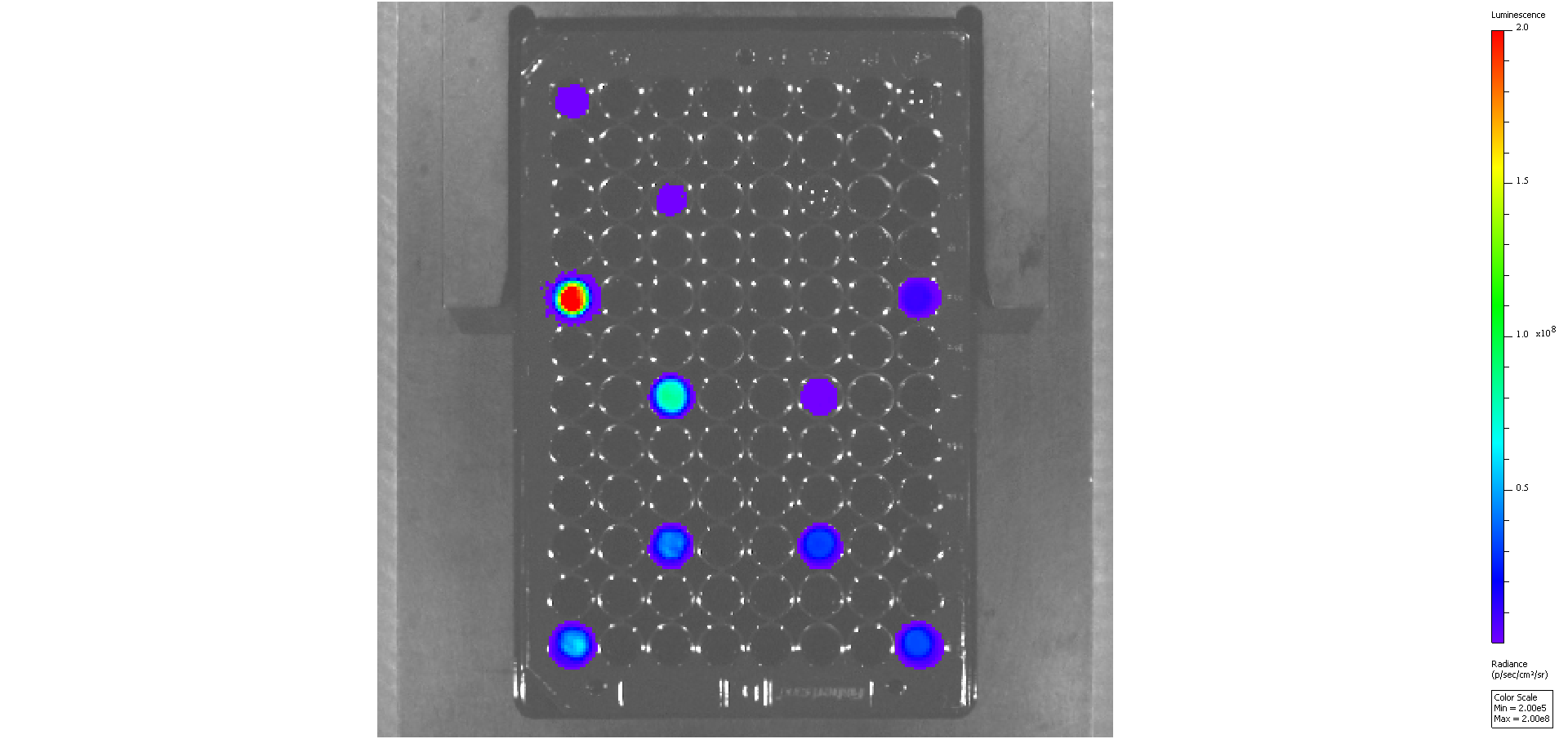

### s12-rep1-allham.png

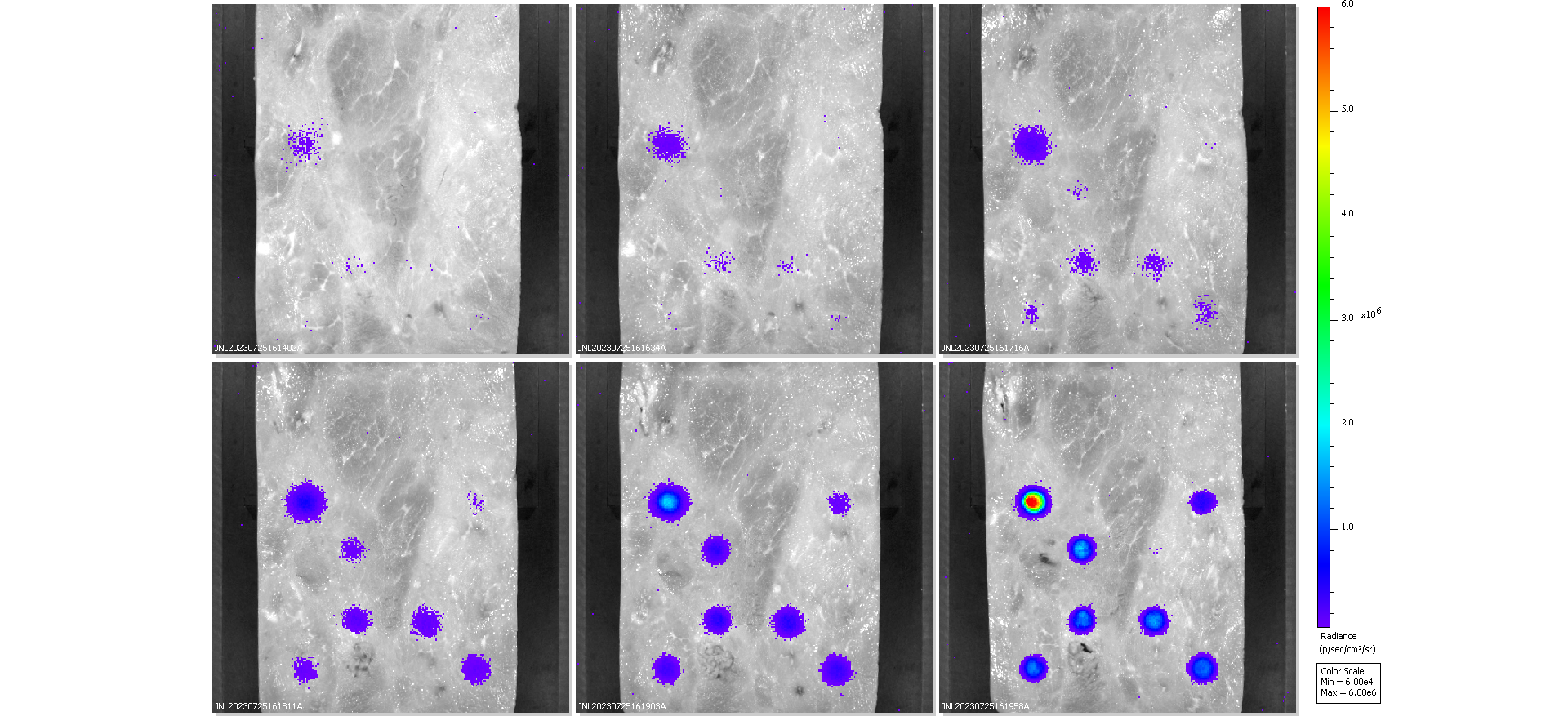

### s12-rep2-0ham.png

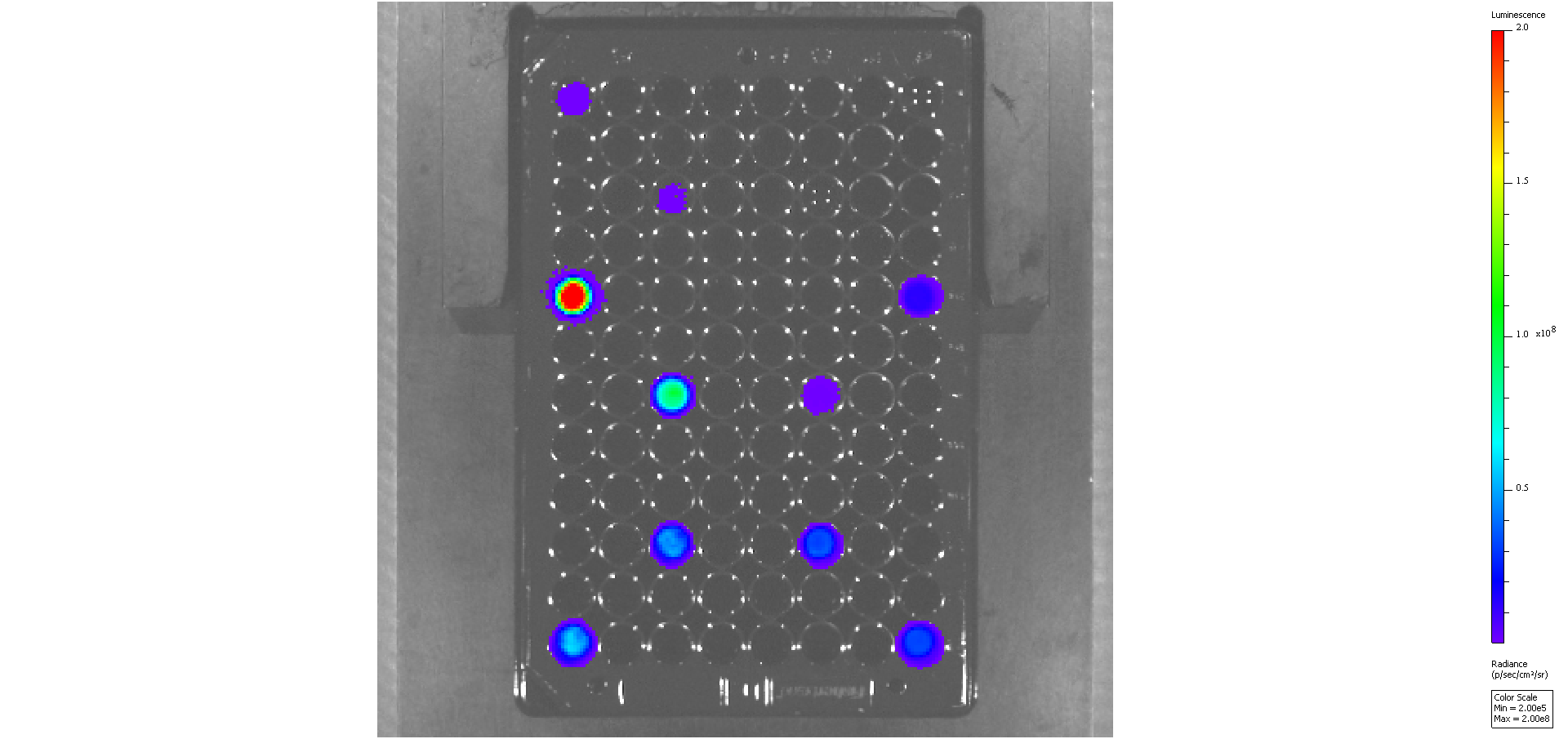

### s12-rep2-allham.png

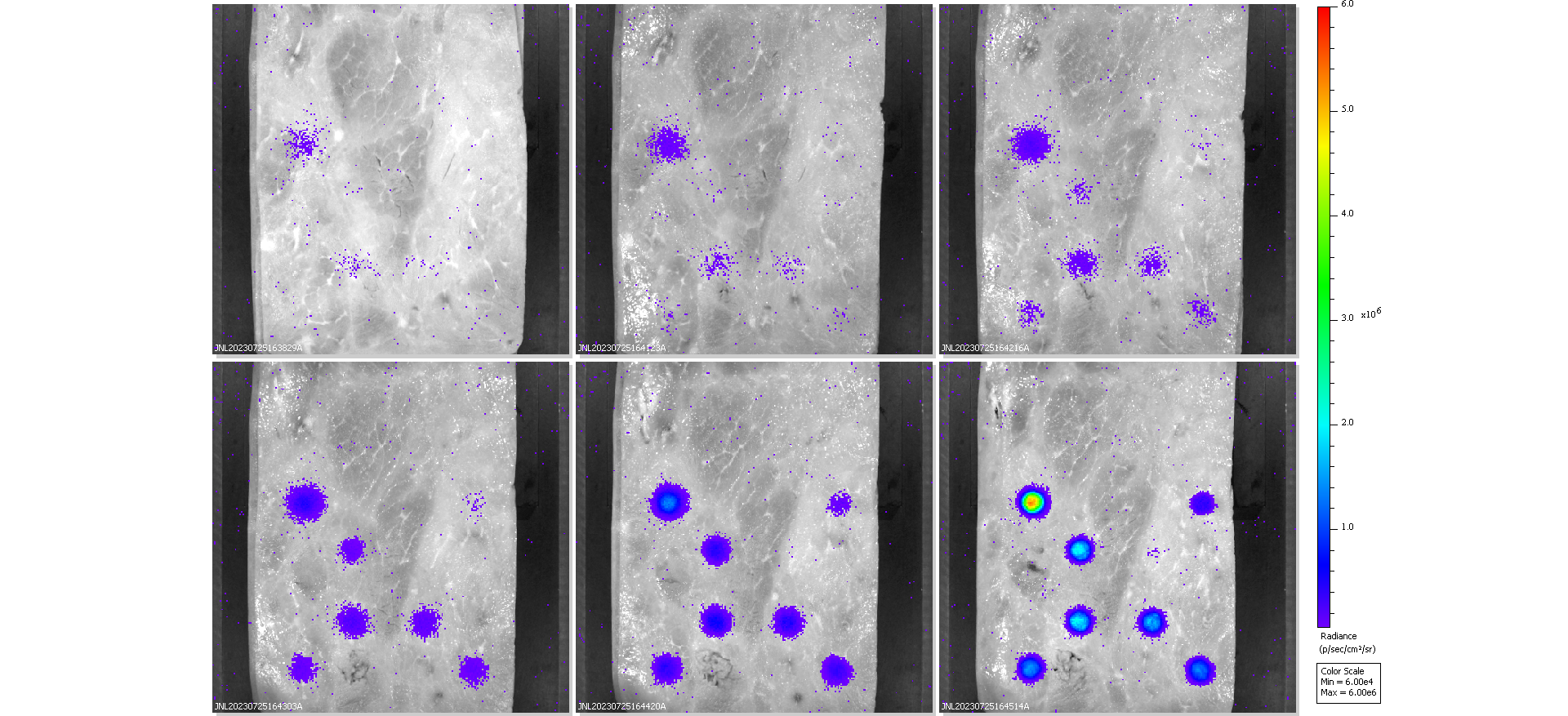

### s12-rep3-0ham.png

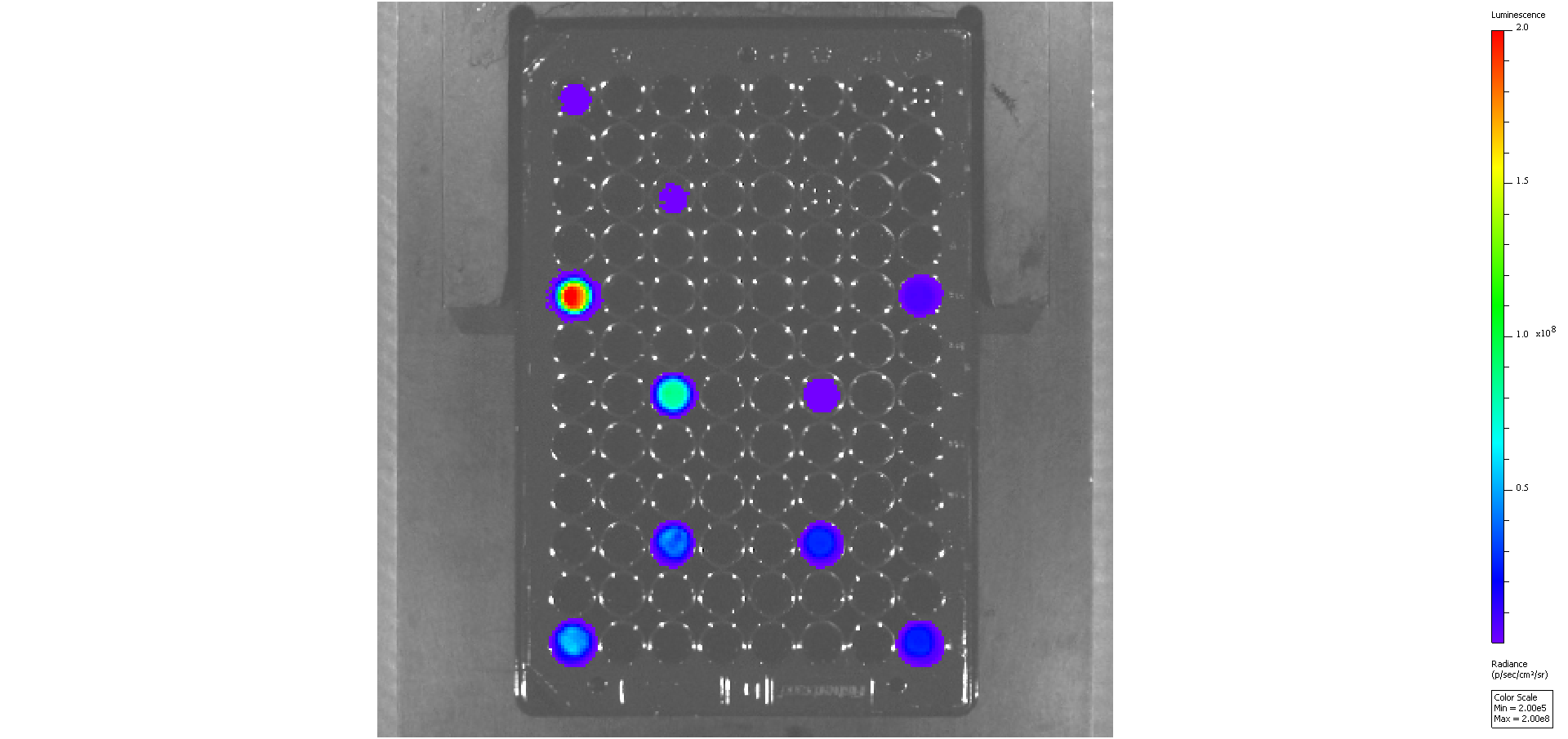

### s12-rep3-allham.png

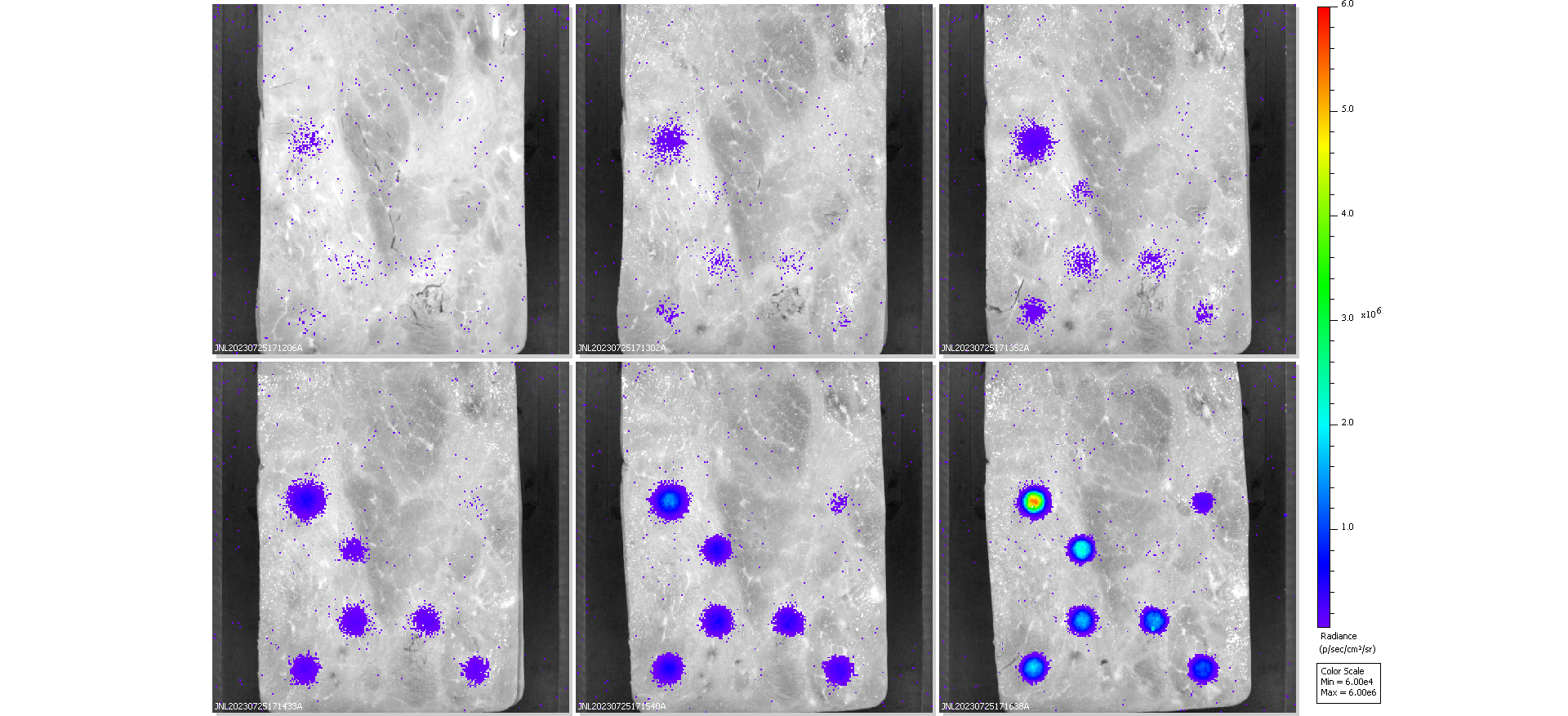
